# Exploratory analysis of multiple traits co-adaptations in the population history

**DOI:** 10.1101/452581

**Authors:** Reiichiro Nakamichi, Shuichi Kitada, Hirohisa Kishino

## Abstract

During the history of range expansion, the populations encounter with variety of environments. They respond to the local environments by modifying the mutually interacting traits. Therefore, to understand the whole life history of the populations, it is ideal to capture the history of their range expansion with reference to the series of surrounding environments and to infer the coadaptation of the multiple traits. Toward this end, we provide an exploratory analysis based on the features of populations: site frequency spectra of populations, population-specific *F*_ST_, association between genes and environments, positive selections on traits mapped on the admixture graph, and GWAS results. Correspondence analysis of genes, environments, and traits provides a bird’s-eye view of the history of population differentiation and range expansion and various types of environmental selections at the times. Principal component analysis of the estimated trait-specific polygenic adaptations mapped on the admixture graph enables to understand the coadaptation of multiple traits. The potential usefulness was confirmed by analyzing a public dataset of wild poplar in northwestern America. In response to the northern cold temperature and longer daylength, the populations increased the photosynthetic activity and nutrient use efficiency at the expense of the risk of pathogen invasion, and in response to warm temperature, they increased the growth. At higher altitude, they shifted the maximum activity to earlier period in spring to reduce the activity in dry summer. The R codes for our representation method and simulations of population colonization used in this study are available as supplementary script.

## 1 INTRODUCTION

Populations adapt to new environments through selection on pre-existing alleles and/or new mutations in adaptation-related loci in the genomes (Barrett & Schluter, 2008). Therefore, adaptation of populations of a species to novel environments changes allele frequencies of loci under selection. Environmental adaptation processes can also create significant differences in phenotypes and traits among populations of a species. When correlated with variation in environmental factors over local subpopulations (hereafter, populations), such variation in traits and phenotypes may reflect phenotypic plasticity or genetic adaptation of the populations. Coop, Witonsky, Di Rienzo, & Pritchard (2010) proposed to detect significant correlations between the SNP allele frequencies and the environmental variables, bypassing the trait variables. Through the annotation of the identified SNPs, it may be possible to characterize the type of adaptation.

Adaptation to environmental factors can change traits and phenotypes of a species, thereby creating population structure underpinned by functional loci. Geographical isolation, which can lead to reproductive isolation and consequent differences in allele frequencies of neutral loci, also contributes to population structuring (Wright, 1965). Divergent selection in an environmental gradient may affect genome-wide population structure (Nosil, Funk, & Ortiz-Barrientos, 2009; Orsini, Vanoverbeke, Swillen, Mergeay, & De Meester, 2013). Empirical studies showed that aridity gradients caused geographically structured populations of Poaceae characterized by cytotype segregation of diploids and allotetraploids (Manzaneda et al., 2012). Geographic distance and habitat differences between populations impacted population structure of marine species (Bradbury & Bentzen, 2007; Jorde et al., 2015; Kitada, Nakamichi, & Kishino, 2017). Therefore, population structure needs to be considered when analyzing correlations among genes, traits, and environmental factors across population samples taken from a wide range of geographical regions.

Genome-wide association studies (GWASs) are widely used to identify associations between genes and traits/environments (Visscher et al., 2017). When data are obtained from a metapopulation exhibiting population structure, the effect of genotypes can be inferred by eliminating population structure effects (Devlin & Roeder, 1999) to avoid spurious associations (Pritchard & Rosenberg, 1999). One representative software program, TASSEL (Yu et al., 2006; Bradbury et al., 2007), performs this type of analysis using a unified mixed model. Alternatively, a structured population can be decomposed into Hardy–Weinberg populations, and the associations tested for each population (Pritchard, Stephens, Rosenberg, & Donnelly, 2000). Future challenges for large-scale GWASs from wild populations (wild GWASs) include development of methods that take population structure into account (Santure & Garant, 2018). Even greater challenge is phenotypic plasticity, which may be identified as a systematic error in the genetic models.

So-called “genome scan methods” consider geographically structured populations and detect SNPs related to environmental variables, traits, and phenotypes (De Mita et al., 2013; De Villemereuil, Frichot, Bazin, François, & Gaggiotti, 2014). For example, BayeScan (Foll & Gaggiotti, 2008) measures the significance of SNP’s locus-specific global *F*_ST_ values, the amount of genetic variations among populations, in Bayesian framework. Genotype-environment associations (GEAs) analyze the allele frequencies of SNPs in sampling locations and test their associations with the environmental variables (Capblancq et al., 2020). Bayenv (Coop, Witonsky, Di Rienzo, & Pritchard, 2010) and the latent factor mixed model (Frichot, Schoville, Bouchard, & François, 2013) can detect SNPs that are highly correlated with environmental factors and traits on the basis of allele frequencies. Notably these methods essentially do not require phenotypic data. Hence, they are valid especially when the life history is complex and cannot be appropriately measured by a few trait variables or the environmental selection on the phenotypes are not characterized (Capblancq et al., 2020). During the evolutionary history of range expansion, the frequencies of existing and derived alleles in a population vary stochastically, and various pressures of environmental selection affect the allele frequencies of related genes and phenotypes. Systematic information on the associations between traits and SNPs in some species such as human (Watanabe et al., 2019) and Arabidopsis (Togninalli et al., 2019) enabled to map adaptive evolution of polygenic traits on the admixture graph (Racimo, Berg, & Pickrel, 2018).

However, the wild populations change their distributions gradually or abruptly generation after generation and encounter with variety of environments. They adapt to the local environments by modifying in balance their multiple traits that are mutually inter-related. For example, populations of sockeye salmon exhibit diversity about life history traits such as spawning time and habitat, and adaptation to local spawning and rearing habitats within complex lake systems (Hilborn, Quinn, Schindler, & Rogers, 2003). Such reproductive traits adapted to specific environment might be controlled by related genes. Gonadotropin-releasing hormone (GnRH) increases in adult salmon brains during homing migration, and controls gonadal maturation during the final phases of upstream migration to spawn (Ueda, 2019). Populations of walking stick insects diverged in body size, shape, host preference, and behavior in parallel with the divergence of their host plant species (Nosil, Crespi, & Sandoval, 2002).

To understand the life history of the populations, it is necessary to overview the history of the range expansion with reference to the newly encountered environments and to capture the coadaptation of the multiple traits under the selections. This paper provides an exploratory approach to characterize the range expansion and the environmental adaptations of the populations through the correspondence analysis of genes, traits, and environments, and the multivariate analysis of traits’ polygenic adaptations mapped on the admixture graph. The R codes for our representation method and simulations of population colonization used in this study are available in the Supporting Information. This approach also accepts SNP genotype data, and reads Genepop format (Raymond & Rousset, 1995; Rousset, 2008).

## 2 MATERIALS AND METHODS

### 2.1 Colored correspondence analysis: history of range expansion and environmental stress

On the basis of the values of the environmental factors, the mean values of the traits, and the allele frequencies at SNPs in each population, correspondence analysis (Benzécri, 1973; Hayashi, 1953) generates a biplot that visualizes the correspondence between the populations and the variables of traits, environments, and genes (SNPs). We had hoped to map the populations with reference to the types of the environment to which they adapt and SNPs with which they adapt. Therefore, as for the allele frequencies at SNPs, we used the frequencies of derived alleles to infer signatures of environmental adaptation. However, it is difficult to know which allele is derived in the actual data without the information on the states in the closely related species. In this paper, we adopted an ad hoc approach of using the frequencies of minor alleles as a substitute, expecting that the frequencies of most of derived alleles tend to be still low. This simple assignment for derived alleles may have errors, but we hoped that it would capture the SNPs that enhanced the allele frequencies to adapt to the local environments. To distinguish the association by positive and negative correlations, we introduced two types of environmental variables: the original environmental value itself, and the sign-reversed value of the original value. Genes and traits that had a positive/negative correlation with the original environmental factors were connected to the original/sign-reversed environmental variables.

To understand the populations in the context of the evolutionary change in the distributional range, we assigned a gradient of colors to the populations. The colors represent population-specific *F*_ST_ (Weir & Goudet, 2017). Population-specific *F*_ST_ estimates the genetic deviation from the ancestral population on the basis of the difference between the heterozygosity of the entire population-pairs and the heterozygosity of each population. The Weir & Goudet’ population-specific *F*_ST_ moment estimator can identify the source population and trace the history of range expansion based on heterozygosity under the assumption that populations closest to the ancestral population have the highest heterozygosity (Kitada, Nakamichi, & Kishino, 2021). We extended the population-specific *F*_ST_ estimator to overall loci as

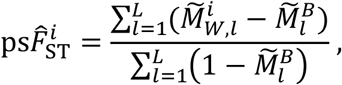

where 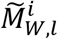 is the unbiased within-population matching of two distinct alleles of locus *l* (*l* = 1 ~ *L*) in population *i* (*i* = 1 ~ *K*), and 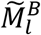 is the between-population-pair matching average over pairs of populations (Buckleton et al., 2016). To interpret the adaptation of the populations, we identified the significant correlations between the genes and the environmental variables (Appendix 1).

### 2.2 PCA of multiple traits polygenic adaptations mapped on the admixture graph

To understand the coadaptation of multiple traits in the life history of the populations, we conducted principal component analysis (PCA) on the outputs of PolyGraph (Racimo, Berg, & Pickrel, 2018). The dynamics of geographical distribution of the populations is first approximated by a set of population differentiation and admixture. Given the allele frequencies of the neutral SNP loci, the pairwise genetic distance among populations can be decomposed into the genetic drifts if we know the history of differentiation and admixture of their ancestral populations. TreeMix (Pickrell & Pritchard, 2012) estimates the structure of this admixture graph by fitting the genetic distances predicted from the scenario of the genetic drifts to the observed genetic distances in the Bayesian framework. Using allele frequencies of SNPs associated with a trait as predictors, PolyGraph estimates the positive selection on the trait occurring along the edges of the admixture graph. The directional changes of their allele frequencies toward the increase/decrease of the trait values are called as positive selection parameters. To obtain the input data required for PolyGraph, we conducted GWAS for each of the traits considered (see Appendix 2). To understand the coadaptation of multiple traits, we made a matrix, by binding the vectors listing the positive selection parameter values for the traits. To see the multiple-traits coadaptation with reference to the environmental variables, we added the among-populations correlations with the environmental variables for each column representing the trait profile. Then, we performed PCA of the estimated positive selection parameters and the environmental factors. Using the factor loading, the positive selection values on the first and the second principal components were calculated as the linear combinations of the trait-specific parameter values and were mapped to the admixture graph.

### 2.3 Simulation of range expansion and adaptation

To illustrate how the overview generated by the exploratory analysis can be interpreted, we conducted a simulation of the colonization and strong environmental selection (Austerlitz, Jung-Muller, Godelle, & Gouyon, 1997) and included *K* (= 25) linearly arrayed populations. Starting from population 1 located at the edge, populations were successively colonized in every 10 generations and exposed to the local environments (Supplementary Figure S1). The allele frequencies of the SNPs in each population varied stochastically by genetic drift, colonization of its ancestral population, and selection pressure of the environments. The environmental factor had two states, *severe* and *normal*, and the values 1 and 0 were assigned to the states. The environmental factor had the value of 0 in most populations. Only populations 9 and 15 were exposed to *severe* environments. The environmental factor does not affect the allele frequencies at the neutral SNPs but affects the allele frequencies of the SNPs that contribute the traits. Adaptation to an environmental stress is often accompanied by the cost of reduced activity in the *normal* environment (e.g., Baucom & Mauricio, 2004); therefore, the derived alleles can adapt to the severe environment at the expense of cost in the *normal* environment (see Appendix 3 for details). The sample size was set to 50 for each of the 25 populations. We simulated the case of strong selection and cost of adaptation.

In this study, we needed to simulate the dynamics of phenotypic traits and the allele frequencies at the loci that contribute to the traits. It was not computationally practical to specify the selection coefficients on the loci directly and to carry out individual-based simulation. As a result, we simulated the dynamics of the **population** allele frequencies and mean traits, assuming that the traits are polygenic and in the form of sum of monogenic latent traits. While the pre-existing alleles as well as the de novo mutations contribute to environmental adaptations, in this simulation, we assumed, for simplicity, that the relevant pre-existing loci were already monomorphic in the ancestral population. Two traits under the selection of environments were polygenic but generated by summing up five latent traits that are monogenic and affected by the environments. The derived allele at the genetic locus contributing to each latent trait had the selection coefficient of *s* = 0.1 in the severe environment and −0.1 in the *normal* environment (see equations A4 and A5 in Appendix 3).

### 2.4 Application to wild poplar data in North America

As an empirical example, we analyzed publicly available data that included genetic and trait information of 441 individuals of wild poplar (*Populus trichocarpa*), which were collected from various regions over a range of 2,500 km near the Canadian–US border at a latitude of 44’ to 59’ N, a longitude of 121’ to 138’ W, and an altitude of 0–800 m (McKown et al., 2014a; McKown et al., 2014b; Geraldes et al., 2013). The data included geographical information of sampling locations, genotypes of 34,131 SNPs (3,516 genes), and values of stomatal anatomy, leaf tannin, ecophysiology, morphology, and disease. These individuals consists of 25 drainages (populations) (Geraldes *et al*. 2014): 9 in northern British Colombia (NBC), 12 in southern British Colombia (SBC), 2 in inland British Colombia (IBC), and 2 in Oregon (ORE). We calculated the averages of the environmental values at the sampling locations and phenotypic values of the individuals, and we considered them representative values of the trait and environment for each population. We plotted the longitudes and latitudes for the individuals on the map (Figure 1). Because our major concern was identifying correlations between among-population differentiations of genes, traits, and environmental factors, we selected the SNP with the highest global *F*_ST_ value over 25 populations from each of the 3,516 gene regions. Here, we focused on the 45 trait variables (Supplementary Table S1; McKown et al., 2014a; McKown et al., 2014b), namely, adaxial stomata density (ADd), abaxial stomata density (ABd), average of two measurements of leaf rust disease morbidity (DP), 14 phenology traits, 12 biomass traits and 16 ecophysiology traits (see Supplementary Table S1). Each sampling location of a population was described by nine environmental/geographical variables: altitude (ALT), longest yearly daylength (photoperiod) (DAY), frost-free days (FFD), mean annual temperature (MAT), mean warmest month temperature (MWMT), mean annual precipitation (MAP), mean summer precipitation (MSP), annual heat–moisture index (AHM, ~MAT/MAP, an indicator of drought), and summer heat–moisture index (SHM, ~MWMT/MSP). The day length and temperature have a north-south cline, while temperature, rainfall, and drought have an east-west (coastal to inland) cline (Geraldes *et al*. 2014). In addition, 18 soil conditions, namely, the ratio of clay, silt, sand, and gravel, soil depth, bulk density, cation exchange capacity, organic carbon, pH, each which were observed in topsoil and subsoil, were obtained from The Unified North American Soil Map (Liu et al., 2013) and used as environmental values of the sampling locations (see Supplementary Table S2).

**Figure 1.**
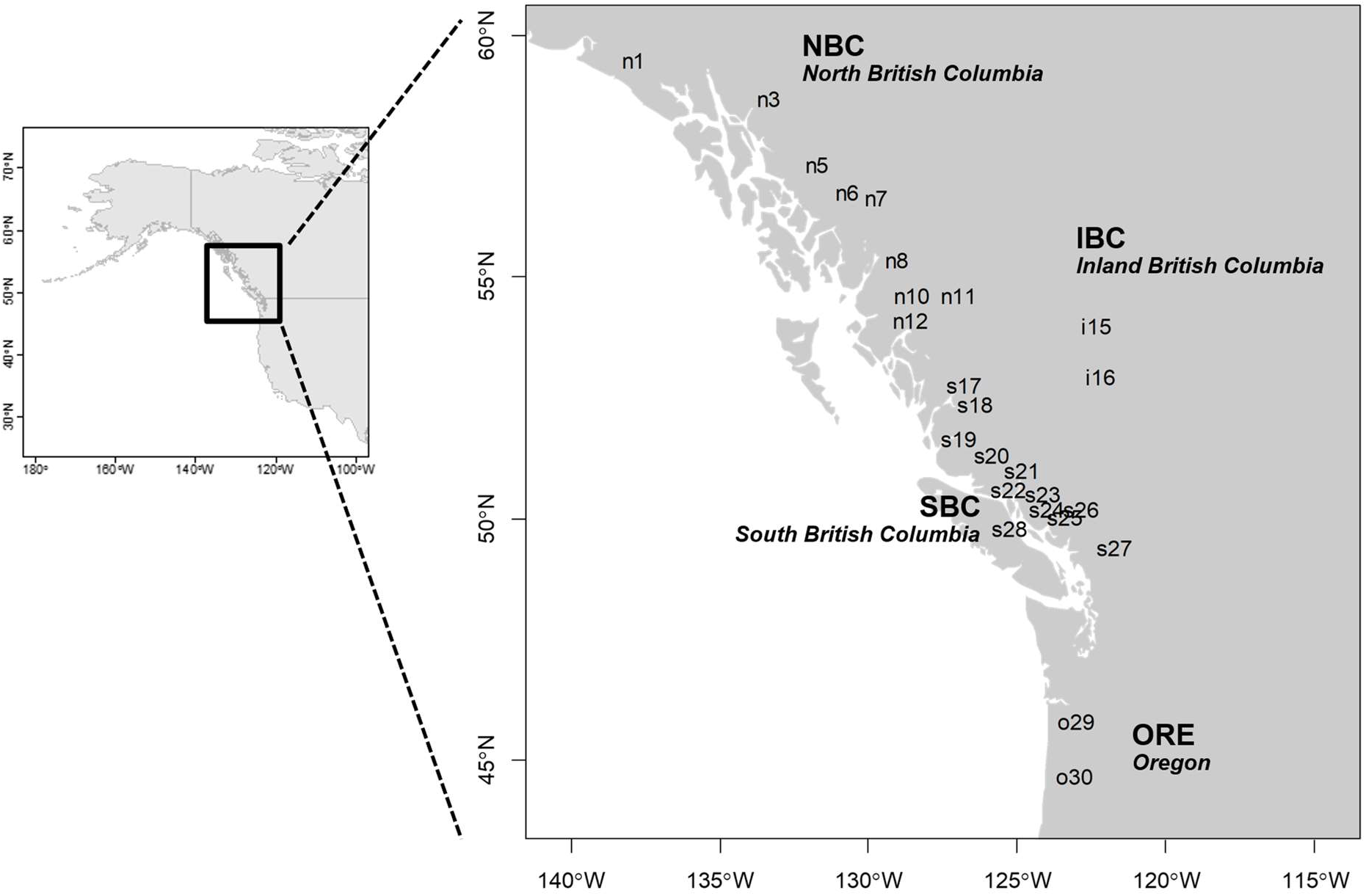
Wild poplar populations in North America. The sampled individuals were classified into 25 populations. Populations were grouped into the regions: North British Columbia (NBC), South British Columbia (SBC), Inland British Columbia (IBC) and Oregon (ORE). The populations were labeled by the first characters of the region names and the id numbers. Data are from Geraldes et al. (2013) and McKown et al. (2014a, 2014b).

## 3 RESULTS

### 3.1 Analysis of simulated data and performance of our method

Figure 2 shows the results of our gene-trait-environment association analysis using simulated data, consisted of the correspondence analysis, population-specific *F*_ST_ values, admixture graph estimated by TreeMix, polygenic trait adaptation estimated by PolyGraph and PCA of estimated positive selection parameters and environmental value. Figure 2a shows the biplot of the correspondence analysis for the genetic–environmental parameter *s* = 0.1; that is, the homozygote of the derived allele is 1.1 times more advantageous than the homozygote of the ancestral allele under the *severe* environmental condition (*E* = 1) and 1.1 times less advantageous under the *normal* environmental condition (*E* = 0) (see Appendix 3 for details). Even adaptive mutations, with fixation probability ~1 – *e*^−2*s*^, have little chance to become dominant. Because of the cost of adaptation, the adaptive mutations were most likely to be deleted immediately, unless they occurred in pop9, pop15, or their immediate colonizers (data not shown).

**Figure 2.**
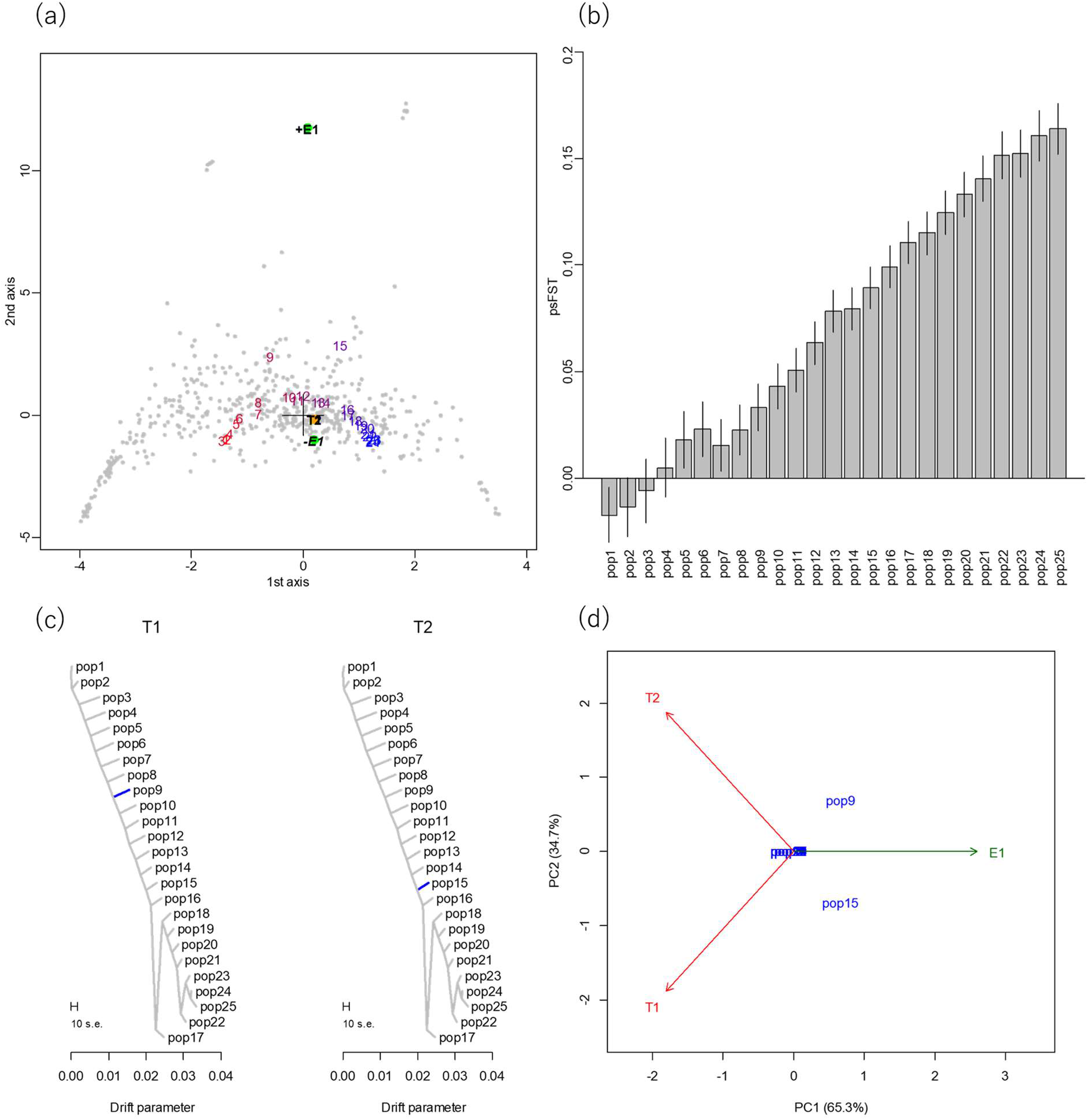
Performance of the exploratory data analysis; simulated data. (a) Correspondence analysis of simulated colonization and adaptation. Green nodes represent environmental factor *E*. The plus sign (+E) indicates the original environmental value, whereas the minus sign (−*E*) represents the sign-reversed environmental value. Gray nodes are neutral and environmentally adaptive loci. The orange node is the observed traits *T*1 and *T*2. Each population label is colored by its population-specific *F*_ST_ value. Numbers from 1 to 25 represent populations, and the color gradients on population labels represent the standardized magnitude of a population-specific *F*_ST_ value at the sampling point, with colors between blue (for the largest *F*_ST_, which represents the youngest population) and red (smallest *F*_ST_, which represents the oldest population). (b) Estimated population-specific *F*_ST_ values. The order of the population-specific *F*_ST_ estimates was stable in 100 simulations, and the point estimates from the first run were plotted with their asymptotic standard errors. (c) Estimated admixture graph and adaptation of traits *T*1 and *T*2. Blue color indicates the decrease of trait values. (d) Principal component analysis (PCA) of estimated positive selection parameters and environments.

The gradient of the colors assigned to the populations in the biplot indicates the history of range expansion on the basis of population-specific *F*_ST_ values and the environmental stresses that the population experienced during the course of range expansion. The environmental factor and environmentally adaptive genes showed positive correlations (connection between green node +*E* and purple nodes). However, the environmental factor and trait showed negative correlation (connection between green node −*E* and the orange node). Most of the populations were characterized by neutral loci; however, pop9 and pop15 were located near the environmental factor. If the populations can be distinguished through neutral loci, this is a sign of isolation-by-adaptation, which requires strong fitness costs to immigration within the population, and thus signs very strong selection. Population-specific *F*_ST_ values and admixture graph (Figure 2b and 2c) reveal that the history that population expansion started around pop1 and extended through pop25.

The estimated positive selection parameters plotted on admixture graph (Figure 2c) shows that the population pop9 decreased the trait T1 when they diverged from pop8 and was exposed to the *severe* environment. Pop15 decreased the trait T2 when they diverged from pop14 and was exposed to the *sever* environment. PCA of positive selection parameters and environment (Figure 2d) represents the pattern of coadaptation of the two traits in response to the environmental selection pressure. The 1st principal component is the opposing axis between the traits and the environment, which explained 65.3% of the variance of the selection parameters and environmental variables. Pop9 and Pop15 decreased the traits under the *sever* state of the environment. The 2nd principal component explain the difference between the two traits, *T*_1_ – *T*_2_, and explained 34.7% variance. The difference was large in pop15 and small in pop9.

### 3.2 Analysis of wild poplar data

#### 3.2.1 Correspondence Analysis

Our generated 2D plot of correspondence and correlation analysis identified the global structure of genetic differentiation and adaptation (Figure 3a). The placement of populations and coloration by population-specific *F*_ST_ provide an interpretation of habitat expansion of three directions, from the inland to the coast and to northern and southern areas (Figure 3b, population s27, which has the lowest value of population-specific *F*_ST_, may be better labeled as “si27” because it is located inland).

**Figure 3.**
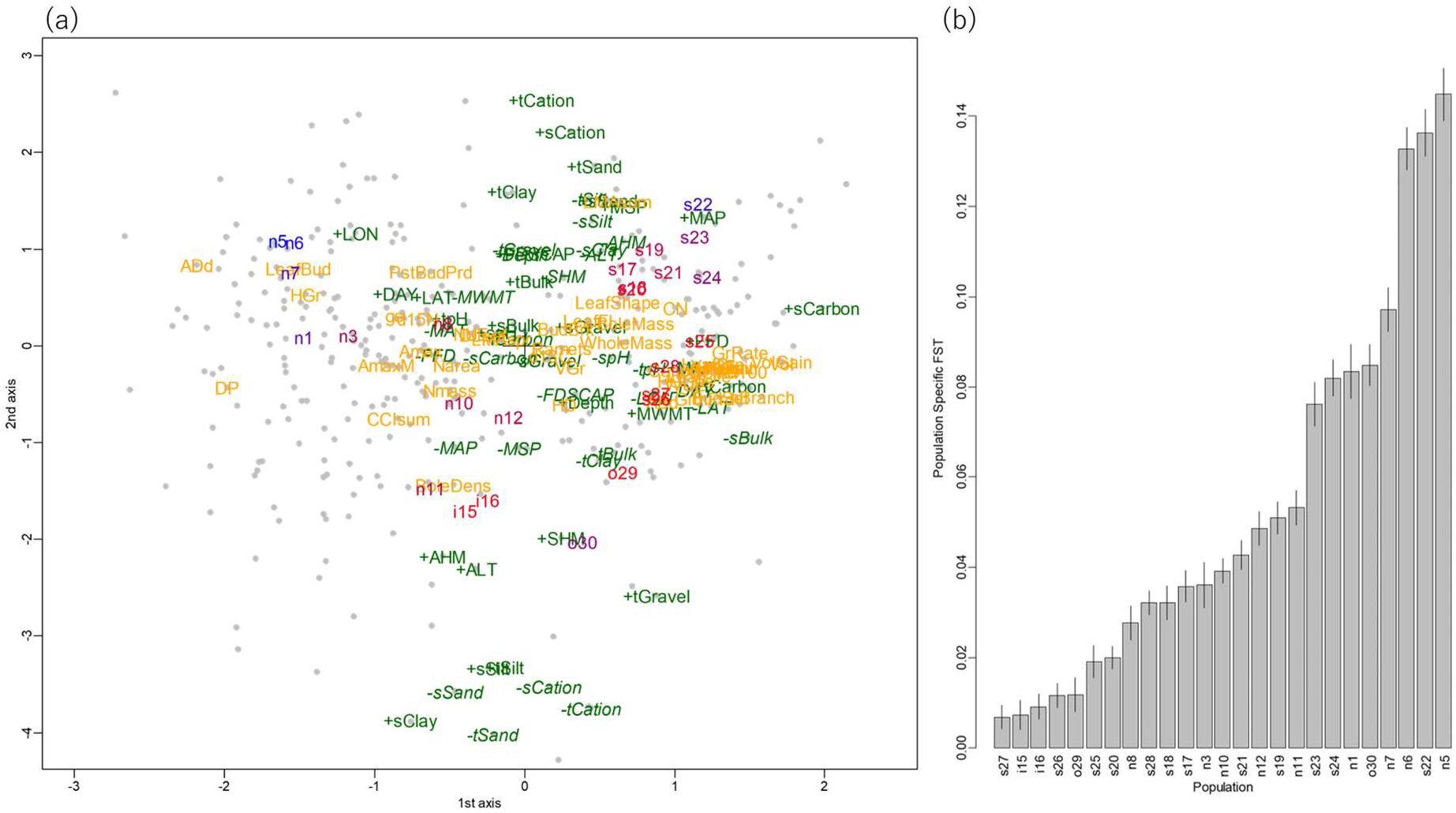
Perspective of the genes, environments, and traits and the life history of the western North American wild poplar populations. (a) Correspondence analysis. Populations are indicated in the cloud of genes (SNPs, marked as dots), environments (colored green), and traits (colored yellow). Environment labels with plus signs represent the original environmental values, whereas environment labels with minus signs (in italic) represent the sign-reversed environmental values. The colors of the populations represent low (red) and high (blue) population-specific *F*_ST_ values. (b) Population-specific *F*_ST_ values of the 25 populations. The heading population labels, “n”, “s”, “i”, and “o” represent populations in the regions of Northern British Columbia, Southern British Columbia, Inland British Columbia, and Oregon respectively. Supplementary Table S1 and S2 list the 45 trait variables and 29 environmental variables shown in the figure.

The color gradient of population-specific *F*_ST_ values on the population labels indicated that the ancestral population might inhabit the inland area (SBC s27 and IBC, i15, i16), which is characterized by high altitude (+ALT) and dry conditions year-round (+AHM, -*MAP*’), as shown in the center of Figure 3a. Dry conditions (+AHM) were correlated with genes associated with drought and osmotic regulation (see Appendix 1, Supplementary Table S3): CBF4 (response to drought and cold stress; Haake et al., 2002, Hussain et al., 2018), XERICO (response to osmotic stress, response to salt stress; Ko, Yang, & Han, 2006), SAL1 (response to water deprivation and salt stress; Wilson et al., 2009), MYB85 (cell wall biogenesis responding water deprivation and salinity; Winter et al., 2007), and APX1 (water deficit; Zandalinas et al. 2016). This indicates that poplar was initially adapted to the dry and cold uplands.

Slightly larger population-specific *F*_ST_ values than those of IBC (Figure 3a,b) indicated that the population expansion might have then occurred to the coastal area (SBC), which was characterized by relatively short day length in summer (-*DAY*); this means that the seasonal variation of day length is small in the southern area, with mild temperatures (+MAT, +FFD) and wet conditions year-round (-*AHM*, +MAP), as plotted in lower left of Figure 3a. The small seasonal variation of day length (-*DAY*) was correlated with abaxial stomata density, which indicated that strong southern sunlight stimulates photosynthesis and requires many stomata. Mild temperatures (+MAT, +FFD) were correlated with genes associated with body growth: GH3.9 (root growth; Khan & Stone, 2007), GSL12 (signaling during growth and development; Yadav et al., 2014), and iqd2 (leaf growth regulator; Nikonorova et al., 2018). The year-round wet environment (-*AHM*) was correlated with a gene related to water conditions, HRA1 (response to hypoxia, Giuntoli et al., 2014). The results indicated that populations in SBC were adapted to warmth and oxygen deprivation due to excessive water. After adaptation in SBC, wild poplar might have expanded to the southern area (ORE), which is characterized by warm and dry conditions, particularly in the summer (+SHM). Dry summer (+SHM) was correlated with stress response in the gene NAC090 (salt and drought tolerance; Zang et al., 2019), which revealed that the population adapted to hot and dry summer conditions. In such dry environment particularly in ORE, the soil consisted of sub soil clay and top soil gravel with little sand to maintain water retention and root growth. Contrarily, the top soil consisted of soil sand and clay with little sub soil silt in SBC to control excessive water to maintain good drainage to prevent root rot with cation absorption (tCation and sCation).

Populations in NBC had large population-specific *F*_ST_ values and small genetic diversity, suggesting that they are young, and that wild poplar expanded to the northern area. This area is characterized by long day length in summer (+DAY), which means that day length varies greatly from season to season, and low temperatures (-*MAT*, -*MWMT*, -*FFD*), as described in Figure 3a. These variables were correlated with adaxial stomata density and leaf rust disease (DP). This finding supports the preceding knowledge that the adaxial stomata compensates for reduced photosynthetic efficiency in the northern area; however, there is a risk of pathogen invasion (Melotto et al., 2006). DAY was correlated with genes associated with light conditions: ACT7 (response to light stimulus; McDowell et al., 1996), PRR7 (circadian rhythm; Alabadí et al., 2001), PRR5 (response to long day condition; Nakamichi et al., 2005) and GA3OX1 (response to red light and gibberellin; Nelson et al., 2010). These results indicated that the population adapted to the light conditions, which vary greatly among seasons.

#### 3.2.2 PCA and the multiple-traits coadaptation mapped on the admixture graph

To interpret the correlations suggested from the correspondence analysis in the context of the life history of the populations, we first estimated the history of population differentiation and admixture by applying TreeMix to the population-specific genotype frequencies of the 34,131 SNPs. Out of the 45 traits, significant associations with genes were detected for 25 traits. For each of the 25 traits, we mapped the positive selection parameters on the admixture graph by utilizing the associated-SNPs allele frequencies (see Appendix 2, Supplementary Table S4) in the populations using PolyGraph (Supplementary Figure S2). Then, we performed PCA of positive selection parameters obtained from PolyGraph and environmental variables (see Methods, Supplementary Table S5).

The 1st principal component explains the coadaptation of the populations during the north-south extension of their distribution. They encountered with change in day length, temperature and chemical content of soil (Figure 4a). Northward extension (higher latitude (LAT)) was accompanied by longer day length (DAY). To adapt to such an environmental change, the populations increased photosynthetic activity (Amax), the rate of gas change (gs), and the efficiency of the nutrient use efficiency (NUE). The increased photosynthetic activity with increased stomatal numbers increased the risk of invasion of pathogen (DP). The southward expansion with higher temperature (mean annual temperature (MAT) and mean warmest month temperature (MWMT)) with more frost-free days (FFD), they had a higher growth (GrRate, HGain), in height (Height) and got more branches (nBranch). Correspondingly, they had longer period of growth (GrthPrd), longer time to bud set (BudSet), and longer lifespan of leaves (LeafLife, Yel) and canopy (CanoDr). Figure 4b summarizes such the predicted increase/decrease of these traits on the admixture graph.

**Figure 4.**
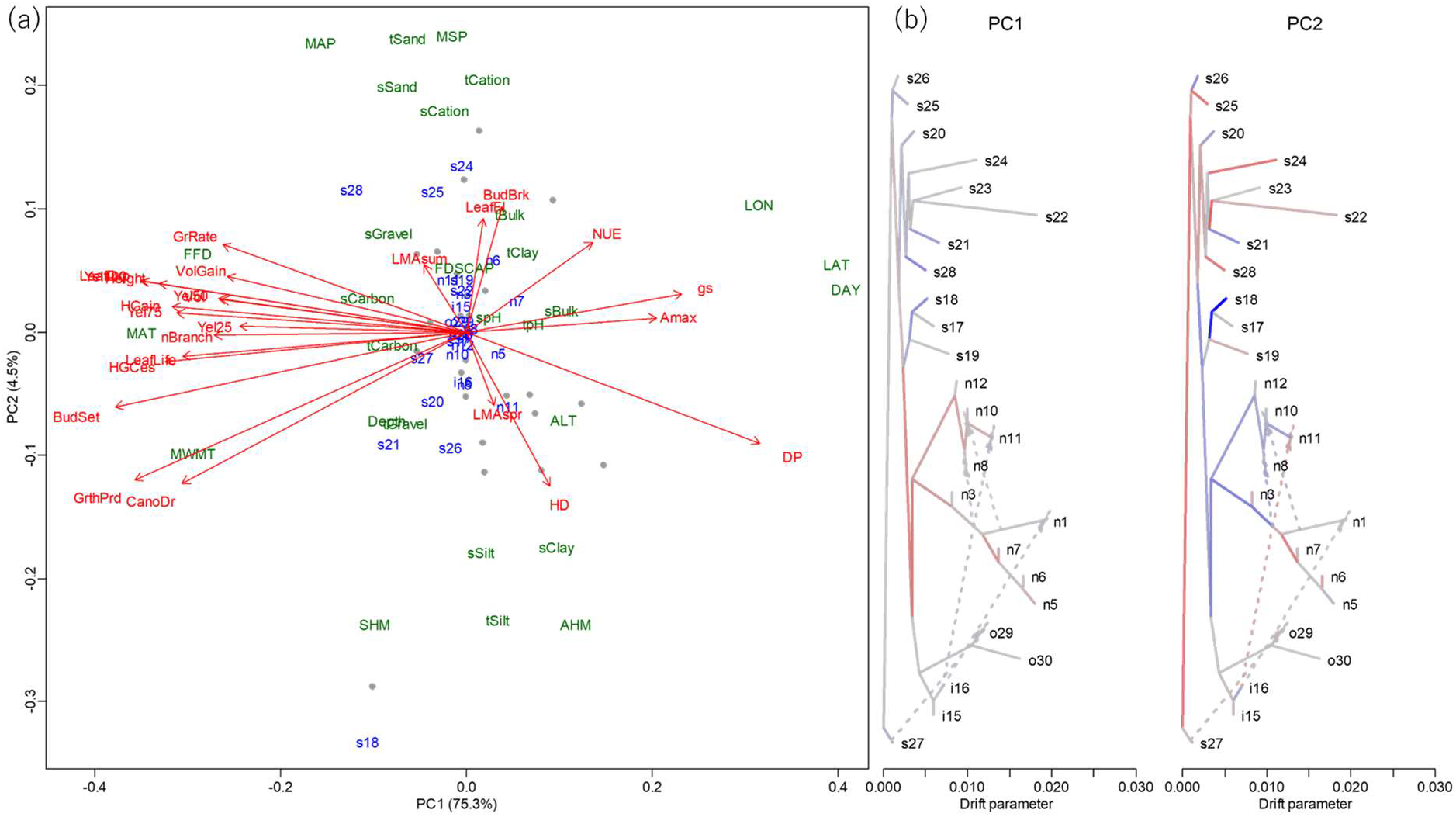
Multiple traits coadaptation in the western North American wild poplar population. (a) Principal component analysis. For each trait, the vector of positive selection parameters that characterize the predicted increase/decrease of the trait value on the admixture graph was estimated using PolyGraph. To understand the coadaptation with reference to the local environments, the among-populations correlation with the environmental variables were added to the vector. The expanded vectors were combined, generating a multivariate data of traits with profiles of positive selection and the correlation with environments (see Materials and Methods). The labels colored blue (populations) and green (environments) represent edges of the admixture graph and the environments. The labels colored red represent traits. Only the terminal edges leading to the current populations are shown and the other internal edges are shown as points. (b) Positive selections of the “principal component traits”. For each principal component, the vector of the positive selection parameter values was obtained as linear combination of the trait-specific selection parameter values with the weight of the factor loadings and mapped on the admixture graph. Red/blue colors represent the selection toward the increase/decrease of the “principal component trait” values.

The 2nd principal component shows the adaptation to the difference of water availability. The environment of larger mean annual precipitation (MAP) and mean summer precipitation (MSP) generated the top and sub soil sandy (tSand and sSand) and of cation (tCation and sCation). Shortage of water with large heat-moisture ratio in summer (SHM) and annually (AHM) at high altitude (ALT) generated fine-grained soil: the top and sub soil of silt (tSilt and sSilt) and clayish top soil (tClay). The populations with less water availability shifted the timing of the growth to earlier period with shortened time to bud break (BudBrk) and leaf flush (LeafFl) by increased activity with large leaf mass per area in spring (LMAspr) and reduced the activity in summer: smaller leaf mass per area in summer (LMAsum). As a result, height/diameter ratio (HD) increased.

## 4 DISCUSSION

A population evolves in space and time and responds to variable environments. From its birth, a population may continuously change its distribution range, and the initial localities may have occasionally been exposed to unprecedented environmental stress. In these localities, individuals and populations can acclimate to such environmental stresses by phenotypic plasticity in a short term, and in a long term, the populations can adapt by changing its geographical distribution or genomes. We focused on the latter and attempted to understand the whole life history of the populations. Toward this end, we conducted an exploratory analysis of multiple-traits coadaptation. The whole scheme of the analysis is multiple layered. In the first layer, we generated the features that are used as inputs for the second layer analysis. They are population-wise SNPs site frequency spectra, genome-wide association with traits and environments, polygenic adaptations mapped on the admixture graph, and obtained by the ever-evolving population genetic and quantitative genetic procedures. The exploratory analysis is the second layer analysis of the features generated in the first layer analysis.

To overview the whole scenario of the historical change of geographical distribution and the adaptation to the new environments at the times, we conducted correspondence analysis that locates populations in relation with the SNPs, environmental variables, and trait variables. From the biplot, the history of range expansion and differentiation were inferred by the values of the population-specific *F*_ST_. To interpret the adaptations, we referred the first layer analysis of association study searching for the genes that are associated with the environmental variables surrounding the characteristic groups of populations.

To understand the coadaptation of multiple traits, we analyzed the correlations among the estimated history of positive selections increasing/decreasing the traits values. They were obtained by the first layer analysis of constructing the admixture graph representing the history of population differentiation and admixture and of mapping the positive selection parameters on the admixture graph based on the spatial allele frequencies of the SNPs associated with the traits.

We attempted to show, through the numerical simulation and an analysis of an empirical data, that the complexity of populations’ life history can be interpreted well solely by integrating the information of among-populations genetic difference, genome-wide association with multiple environments and multiple traits. Multiple layered approach may be a practical choice. Our approach is still in its infancy. One direction of future study is to include latent variables that are interpreted as key elements of environmental selection and adaptation. Another direction is an attempt to quantify the pattern of adaptation. A natural framework is a Bayesian approach that use the features provided by the first layer analysis as the prior information.

As a final remark, we note that our analysis uses populations as units of the analysis. However, populations are often defined by post-stratification of the sample. In the case of wild poplar data, we adopted the population assignment provided by the original dataset. The power of our exploratory approach depends on the accuracy of the assignment of the individuals to local populations. Individual-level analysis deserves consideration.

## Supporting information

Supplementary Script

## ACKNOWLEDGEMENTS

We appreciate the essential comments made by the reviewers that significantly improved the manuscript. This study was supported by Japan Society for the Promotion of Science Grants-in-Aid for Scientific Research KAKENHI nos. 16H02788 and 19H04070 to H.K. We thank Mallory Eckstut, PhD, from Edanz Group for editing a draft of this manuscript.

## DATA ACCESSIBILITY STATEMENT

The authors affirm that all data necessary for confirming the conclusions of the article are present within the article, figures, and supplementary information. The R codes to perform our representation method and simulations of population colonization are available in the Supporting Information.

## AUTHOR CONTRIBUTIONS

H.K. designed the study and developed the theory. R.N. and H.K. analyzed data. R.N. created R package for implementing the method and performed simulations. S.K., H.K., and R.N. provided critical feedback, helped shape the research, and wrote the manuscript.

## Appendix 1 Gene–environment correlation

To interpret the adaptation to each type of the local environments, we identified the significant correlations between the environment and genes (SNPs). We accounted for the correlation structure of the residuals. At each locus, the variance matrix of the observed allele frequencies reflects the genetic drift and gene flow and the sampling variance (Nicholson, Smith, Jónsson, Gústafsson, & Stefánsson, 2002; Coop, Witonsky, Di Rienzo, & Pritchard, 2010). Here, we adopted the frequentist approach to choose significant pairs given a value of false discovery rate (FDR) (1% for the simulation and 5% for the poplar data).

We consider *K* populations derived from a common ancestral population and *L* loci of biallelic neutral markers. Let 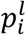 and 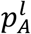 be the derived allele frequency of marker *l* (*l* = 1 − *L*) in population *i* (*i* = 1 − *K*) and the (unobserved) ancestral population. Given the samples from the populations, the allele frequencies are estimated by the observed counts as 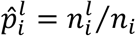, where *n_i_* is the number of the samples (twice of the number of individuals) in population *i*, and 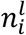 is the count of derived allele at marker *l* in population *i*.

The among-population mean allele frequencies vary largely among neutral loci. Therefore, we incorporate the contribution of variable allele frequencies in the ancestral population to estimate the among-population correlation that is shared among loci. Given the allele frequency in the ancestral population, the variance–covariance matrix of the allele frequencies 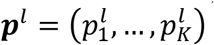 at locus *l* is formulated as

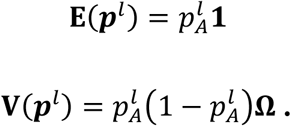

*ν_ij_* = Ω_*ij*_ (*i,j* = 1,…, *K*) represents the among-population covariance (Weir & Hill, 2002; Coop, Witonsky, Di Rienzo, & Pritchard, 2010). The variance and covariance of the observed allele frequencies are

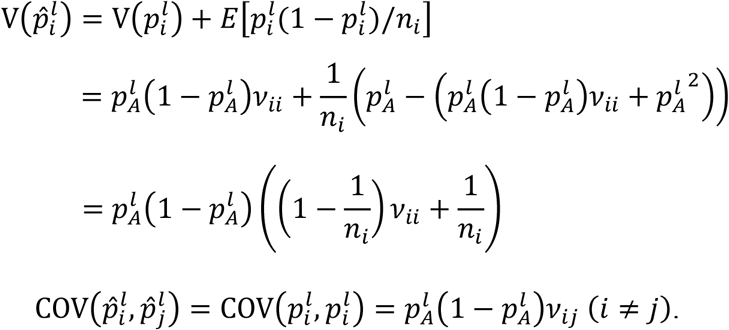

From this, we obtained the moment estimator Ω as

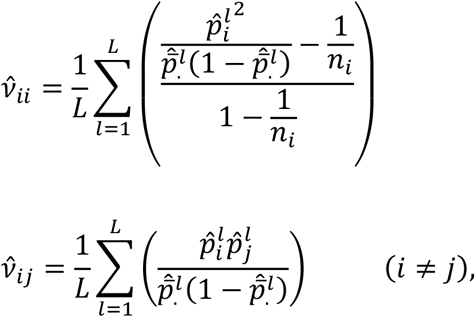

where 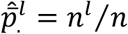. Because most SNPs rarely have alleles in equilibrium (Wright, 1931), these estimates are accurate when many neutral loci are available. With this estimated variance–covariance matrix of the allele frequencies, we obtained the variance–covariance matrix of the observed counts, 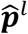, as

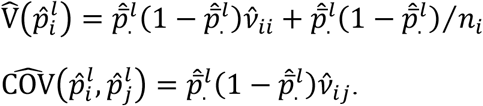

Assuming the normality of the estimated regression coefficient, the p-value was calculated by contrasting the coefficient with the standard error based on the standard generalized least squares (GLS) method. Out of environment–gene (SNP) pairs and environment–trait pairs, we selected the significant pairs with an FDR of 0.05 using the Benjamini–Hochberg procedure (Benjamini & Hochberg, 1995).

To improve the power of detecting associations between genes and environments, we focused on the SNPs that were over-differentiated among populations compared with the level of differentiation of neutral loci. First, we obtained the maximum likelihood estimates of the locus-specific global *F*_ST_ values (Beaumont & Bolding, 2004) using R package `FinePop2` in CRAN. We fitted a gamma distribution to the distribution of these locus-specific global *F*_ST_ values by maximum likelihood procedure by using the function. The *F*_ST_ values were far below 1, at least in the simulation and in the real data analysis (see Supplementary Figure S3). We assumed that most of the SNPs were neutral, and that the fitted distribution approximates the distribution of locus-specific global *F*_ST_ values of neutral sites. As a set of over-differentiated SNPs, we collected the SNPs with *F*_ST_ values with upper p-value < 0.1 in this gamma distribution.

## Appendix 2 Kernel-based GWAS and estimation of the effects

In order to make the dataset for PolyGraph analysis, we conducted GWAS for each of the traits using genome-wide gene-based analysis by considering genes as testing units (Deng et al. 2020). For a given gene, joint effect of multiple SNPs within the gene is obtained by Gaussian kernel function. Association between a trait and the candidate kernel function of a gene is evaluated by the generalized association test based on *U*-statistics which uses environmental factors as the fixed effects to control the correlation structure of the populations. Genes which have significant association with the trait were selected by 5% of false discovery rate. We adopted these genes as the explanatory variables which explain the population adaptation in the PolyGraph model. Finally, for each trait, we performed simple linear regression on each of the SNPs on the significant genes on the trait, and estimated regression coefficient of the gene. Then, we adopted the sign (+1 or −1) of the coefficient as the selective pressure in the PolyGraph analysis. For the analysis of poplar, we performed this association test between 45 traits and 3,516 genes and obtained 22 traits which had significant genes (see Supplementary Table S3).

## Appendix 3 Simulation scenario

### A3.1 Populations, SNPs and traits

In this study, we needed to simulate the dynamics of phenotypic traits and the allele frequencies at the loci that contribute to the traits. As a result, it was not practical to specify the selection coefficients on the loci directly. Because of the computational burden, we simulated the dynamics of the population allele frequencies and mean traits based on the expected randomness and survival probability of the alleles in 1-d stepping-stone system of *K* = 25 linearly arrayed populations (see Supplementary Figure S1). We assumed that the traits are polygenic and in the form of sum of monogenic latent traits. While the pre-existing alleles as well as the de novo mutations contribute to environmental adaptations, we assumed, for simplicity, that the relevant pre-existing loci were already monomorphic in the ancestral population. We simulated the case of strong selection and cost of adaptation.

Population 1 accommodated an ancestral population of *N_e_* = 10^5^. In every generation, the populations exchanged 1% of *N_e_* individuals with adjacent populations. Once in 10 generations, habitat expansion occurred, and 1% of *N_e_* immigrated to the adjacent vacant population and increased the population size to the capacity *N_e_* in one generation. We introduced an environmental factor that had a *severe* (1) state in populations 9 and 15, and *normal* (0) state in the other populations:

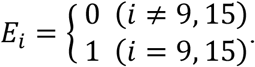

This did not affect the allele frequencies of neutral alleles but affected the survival of non-neutral alleles.

For the initial population, we generated 10,000 polymorphic loci whose alleles were neutral against the environment. Their allele frequencies were set to the theoretical equilibrium distribution, *f*(*q*) ∝ *q*^−1^(1 – *q*)^−1^ (Wright, 1931). Then, additional polymorphisms of 50 neutral loci and 10 environmentally adaptive loci were introduced to the existing populations in each generation. The current genetic diversity reflects the genetic drift of polymorphic loci in the ancestral populations and that of de novo mutations that occurred in the history of populations. To include the latter effect, we generated neutral mutations as well. For computational reason, the initial frequency of the derived alleles at the newly generated loci was set to 0.01 in the populations where mutations occur and 0 in the other populations; they mimicked new mutations that survived the initial phase after their birth. Population allele frequencies varied with random drift under a binomial distribution. Each of the ten environmentally adaptive loci contributed to a latent monogenic trait whose survival was affected by the state of the environment (see Appendix A3.2). Two polygenic traits *T*1 and *T*2 were formed as a sum of five of the ten latent traits respectively.

After 260 generations, we obtained loci that retained their polymorphism. As a simplified procedure that mimic SNP discovery process, we randomly selected a prespecified number of SNPs. In this simulation, we selected 5,000 initial neutral loci, 50 newly derived neutral loci, and two sets of five (totally 10) newly derived environmentally adaptive loci. Then, we generated the allele frequencies of the sample consisting of 50 individuals for each population.

### A3.2 Environmental selection and fitness of derived alleles contributing to the latent traits

In a neutral gene, the allele frequencies are changed by random drift. However, in an environmentally adaptive gene, derived alleles have advantages/disadvantages in *severe*/*normal* conditions compared with the ancestral allele. Therefore, derived allele frequency increases/decreases in *severe*/*normal* conditions by natural selection. We inferred the environmental adaptation and the cost on the basis of the survival probability of the relevant trait.

The genotype *G* = 0,1,2 of the environmental adaptation locus contributes to a latent trait *T*(*G,E*) with the interaction of genotype *G* and environmental factor *E*:

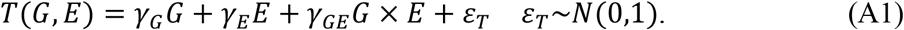

The survival probability *S*(*T*) of the trait value *T* is described as the probability that *T* is positive:

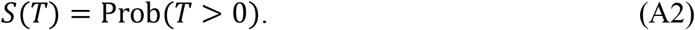

The larger the trait, the greater chance of survival. The survival probability of a genotype *G* under environmental condition *E, S*(*G|E*), is given as *S*(*G|E*) = *S*(*T*(*G,E*)). In population *i*, given the frequency 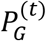 of genotype *G* at generation *t*, the allele frequency at the next generation is obtained as 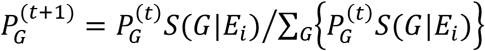. We note that only the relative values of survival probabilities are relevant for the population genetic dynamics.

The derived allele is assumed to be advantageous over the ancestral allele under the *severe* environmental condition *E* = 1, whereas it is disadvantageous under the *normal* environmental condition *E* = 0. Therefore, we consider the case where *γ_G_* ≤ 0, *γ_E_* ≤ 0, and *γ_GE_* ≥ 0. We set a simulation parameter *r* and *s* to control environmental effect *γ_E_*, genetic effect *γ_G_*, and gene-environment interaction *γ_GE_*. The parameter *r* (0 < *r* < 1) was introduced to represent the stress of *severe* environmental condition *E* = 1 and was defined by the ratio of the survival probabilities between the two conditions:

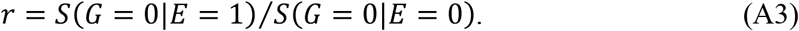

The parameter *s* (≥ 0) represents the fitness of the derived allele in the *severe* environmental condition *E* = 1:

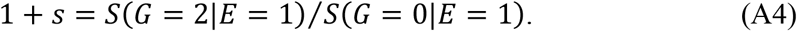

We assumed the cost of adaptation by reversing the fitness in the *normal* environmental condition *E* = 0:

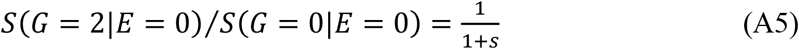

The coefficients *γ_E_*, *γ_G_*, and *γ_GE_* were obtained from *r* and *s*. First, we noted that, from equations (A1) and (A2), 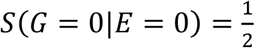 and *S*(*G* = 0|*E* = 1) = Φ(*γ_E_*). Φ is the cumulative distribution of the standard normal distribution. Hence, from equation (A3), we obtained 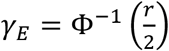. Similarly, we obtained 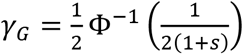 from equation (A5). Finally, we obtained 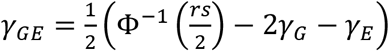 from equation (A4).

## REFERENCES

Alabadí, D., Oyama, T., Yanovsky, M. J., Harmon, F. G., Más, P., & Kay, S. A. (2001). Reciprocal regulation between TOC1 and LHY/CCA1 within the Arabidopsis circadian clock. Science, 293, 880–883. https://doi.org/10.1126/science.1061320

Austerlitz, F., Jung-Muller, B., Godelle, B., & Gouyon, P.-H. (1997). Evolution of coalescence times, genetic diversity and structure during colonization. Theoretical Population Biology, 51, 148–164. https://doi.org/10.1006/tpbi.1997.1302

Barrett, R.D., & Schluter, D. (2008). Adaptation from standing genetic variation. Trends in Ecology and Evolution, 23, 38–44. https://doi.org/10.1016/j.tree.2007.09.008

Baucom, R. S. & Mauricio, R. (2004). Fitness costs and benefits of novel herbicide tolerance in a noxious weed. Proceedings of the National Academy of Sciences, USA, 101, 13386–13390. https://doi.org/10.1073/pnas.0404306101

Beaumont, M. A., & Balding, D. J. (2004). Identifying adaptive genetic divergence among populations from genome scans. Molecular Ecology, 13, 969–980. https://doi.org/10.1111/j.1365-294X.2004.02125.x

Benjamini, Y., & Hochberg, Y. (1995). Controlling the false discovery rate: a practical and powerful approach to multiple testing. Journal of the Royal Statistical Society: Series B (Statistical Methodology), 57, 289–300. https://doi.org/10.1111/j.2517-6161.1995.tb02031.x

Benzécri, J. P. (1973). Data Analyses. Volume II. Correspondence Analysis. Paris, France: Dunod.

Bradbury, I. R. & Bentzen, P. (2007). Non-linear genetic isolation by distance: implications for dispersal estimation in anadromous and marine fish populations. Marine Ecology Progress Series, 340, 245–257. doi:10.3354/meps340245

Bradbury, P. J., Zhang, Z., Kroon, D. E., Casstevens, T. M., Ramdoss, Y., & Buckler, E. S. (2007). TASSEL: software for association mapping of complex traits in diverse samples. Bioinformatics, 23, 2633–2635. https://doi.org/10.1093/bioinformatics/btm308

Buckleton, J., Curran, J., Goudet, J., Taylor, D., Thiery, A., & Weir, B. S. (2016). Population-specific *F*_ST_ values for forensic STR markers: A worldwide survey. Forensic Science International: Genetics, 23, 91–100. https://doi.org/10.1016/j.fsigen.2016.03.004

Capblancq, T., Fitzpatrick, M. C., Bay, R. A., Exposito-Alonso, M., & Keller, S. R. (2020). Genomic prediction of (mal) adaptation across current and future climatic landscapes. Annual Review of Ecology, Evolution, and Systematics, 51, 245–269. https://doi.org/10.1146/annurev-ecolsys-020720-042553

Coop, G. D., Witonsky, D., Di Rienzo, A., & Pritchard, K. J. (2010). Using environmental correlations to identify loci underlying local adaptation. Genetics, 185, 1411–1423. https://doi.org/10.1534/genetics.110.1148

De Mita, S., Thuillet, A. C., Gay, L., Ahmadi, N., Manel, S., Ronfort, J., & Vigouroux, Y. (2013). Detecting selection along environmental gradients: analysis of eight methods and their effectiveness for outbreeding and selfing populations. Molecular Ecology, 22, 1383–1399. https://doi.org/10.1111/mec.12182

Deng, Y., He, T., Fang, R., Li, S., Cao, H., & Cui, Y. (2020). Genome-wide gene-based multi-trait analysis. Frontiers in Genetics, 11, 437. doi:10.3389/fgene.2020.00437

De Villemereuil, P., Frichot, É., Bazin, É., François, O., & Gaggiotti, O. E. (2014). Genome scan methods against more complex models: when and how much should we trust them? Molecular Ecology, 23, 2006–2019. https://doi.org/10.1111/mec.12705

Devlin, B., & Roeder, K. (1999). Genomic control for association studies. Biometrics, 55, 997–1004. https://doi.org/10.1111/j.0006-341X.1999.00997.x

Foll, M., & Gaggiotti, O. (2008). A genome scan method to identify selected loci appropriate for both dominant and codominant markers: a Bayesian perspective. Genetics, 180, 977–993. https://doi.org/10.1534/genetics.108.092221

Fraley, C., & Raftery, A. E. (2016). Model-based clustering, discriminant analysis and density estimation. Journal of the American statistical Association, 97, 611–631. https://doi.org/10.1198/016214502760047131

Frichot, E., Schoville, S. D., Bouchard, G., & François, O. (2013). Testing for associations between loci and environmental gradients using latent factor mixed models. Molecular Biology and Evolution, 30, 1687–1699. https://doi.org/10.1093/molbev/mst063

Geraldes, A., Difazio, S. P., Slavov, G. T., Ranjan, P., Muchero, W., … & Tuskana, G. A. (2013). 34K SNP genotyping array for Populus trichocarpa: Design, application to the study of natural populations and transferability to other Populus species. Molecular Ecology Resources, 13, 306–323. https://doi.org/10.1111/1755-0998.12056

Geraldes, A., Farzaneh, N., Grassa, C. J., McKown, A. D., Guy, R. D., Mansfield, S. D., … & Cronk, Q. C. (2014). Landscape genomics of Populus trichocarpa: the role of hybridization, limited gene flow, and natural selection in shaping patterns of population structure. Evolution, 68, 3260–3280. https://doi.org/10.1111/evo.12497

Giuntoli, B., Lee, S. C., Licausi, F., Kosmacz, M., Oosumi, T., … & Perata, P. (2014). A trihelix DNA binding protein counterbalances hypoxia-responsive transcriptional activation in Arabidopsis. PLOS Biology, 12, e1001950. https://doi.org/10.1371/journal.pbio.1001950

Haake, V., Cook, D., Riechmann, J. L., Pineda, O., Thomashow, M. F., & Zhang, J. Z. (2002). Transcription factor CBF4 is a regulator of drought adaptation in Arabidopsis. Plant Physiology, 130, 639–648. https://doi.org/10.1104/pp.006478

Hayashi, C. (1953). Multidimensional quantification: With applications to analysis of social phenomena. Annals of the Institute of Statistical Mathematics, 5, 121–143. https://www.ism.ac.jp/editsec/aism/pdf/005_2_0121.pdf

Hilborn, R., Quinn, T. P., Schindler, D. E., & Rogers, D. E. (2003). Biocomplexity and fisheries sustainability. Proceedings of the National Academy of Sciences, USA, 100, 6564–6568. https://doi.org/10.1073/pnas.1037274100

Hussain, H.A., Hussain, S., Khaliq, A., Ashraf, U., Anjum, S.A., Men, S., & Wang, L. (2018). Chilling and Drought Stresses in Crop Plants: Implications, Cross Talk, and Potential Management Opportunities. Frontiers in Plant Science, 9, 393. https://doi.org/10.3389/fpls.2018.00393

Jorde, P. E., Søvik, G., Westgaard, J. I., Albretsen, J., André, C., …. & Jørstad, K. E. (2015). Genetically distinct populations of northern shrimp, *Pandalus borealis*, in the North Atlantic: adaptation to different temperatures as an isolation factor. Molecular Ecology, 24, 1742–1757. https://doi.org/10.1111/mec.13158

Khan, S. & Stone, J. M. (2016). Arabidopsis thaliana GH3.9 influences primary root growth. Planta, 226, 21–34. https://doi.org/10.1007/s00425-006-0462-2

Kitada, S., Nakamichi, R., & Kishino, H. (2017). The empirical Bayes estimators of fine-scale population structure in high gene flow species. Molecular Ecology Resources, 17, 1210–1222. https://doi.org/10.1111/1755-0998.12663

Kitada, S., Nakamichi, R., & Kishino, H. (2021). Understanding population structure in an evolutionary context: population-specific *F*_ST_ and pairwise *F*_ST_. G3 Genes|Genomes|Genetics, in press. https://doi.org/10.1093/g3journal/jkab316

Ko, J. H., Yang, S. H., & Han, K. H. (2006). Upregulation of an Arabidopsis RING-H2 gene, XERICO, confers drought tolerance through increased abscisic acid biosynthesis. Plant Journal, 47, 343–355. https://doi.org/10.1111/j.1365-313X.2006.02782.x

Liu, S., Wei, Y., Post, W. M., Cook, R. B., Schaefer, K., & Thornton, M. M. (2013). The Unified North American Soil Map and its implication on the soil organic carbon stock in North America. Biogeosciences, 10, 2915–2930. https://doi.org/10.5194/bg-10-2915-2013

Manzaneda, A. J., Rey, P. J., Bastida, J. M., Weiss-Lehman, C., Raskin, E., & Mitchell-Olds, T. (2012). Environmental aridity is associated with cytotype segregation and polyploidy occurrence in Brachypodium distachyon (Poaceae). New Phytologist, 193, 797–805. https://doi.org/10.1111/j.1469-8137.2011.03988.x

McDowell, L. M., An, Y., Huang, S., McKinney, E. C., & R. B. Meagher, R. B. (1996). The Arabidopsis ACT7 actin gene is expressed in rapidly developing tissues and responds to several external stimuli. Plant Physiology, 111, 699–711. https://doi.org/10.1104/pp.111.3.699

McKown, A. D., Guy, R. D., Klápště, J., Geraldes, A., Friedmann, M., … & Douglas, C. J. (2014a). Geographical and environmental gradients shape phenotypic trait variation and genetic structure in *Populus trichocarpa*. New Phytologist, 201, 1263–1276. https://doi.org/10.1111/nph.12601

McKown. A. D., Guy, R. D., Quamme, L., Klápště, J., La Mantia, J., … & Azam M. S. (2014b). Association genetics, geography and ecophysiology link stomatal patterning in *Populus trichocarpa* with carbon gain and disease resistance trade-offs. Molecular Ecology, 23, 5771–5790. https://doi.org/10.1111/mec.12969

Melotto, M., Underwood, W., Koczan, J., Nomura, K., & He, S. Y. (2006). Plant stomata function in innate immunity against bacterial invasion. Cell, 126, 969–980. https://doi.org/10.1016/j.cell.2006.06.054

Nakamichi, N., Kita, M., Ito, S., Sato, E., Yamashino, T., & Mizuno, T. (2005). The Arabidopsis Pseudo-response Regulators, PRR5 and PRR7, Coordinately Play Essential Roles for Circadian Clock Function. Plant and Cell Physiology, 46, 609–619. https://doi.org/10.1093/pcp/pci061

Nelson, D. C., Flematti, G. R., Riseborough, J.-A., Ghisalberti, E. L., Dixon, K. W., & Smith, S. M. (2010). Karrikins enhance light responses during germination and seedling development in Arabidopsis thaliana. Proceedings of the National Academy of Sciences, USA, 107, 7095–7100. https://doi.org/10.1073/pnas.0911635107

Nicholson, G., Smith, A. V., Jónsson, F., Gústafsson, O., Stefánsson, K., & Donnelly, P. (2002). Assessing population differentiation and isolation from single-nucleotide polymorphism data. Journal of the Royal Statistical Society: Series B (Statistical Methodology), 64, 695–715. https://doi.org/10.1111/1467-9868.00357

Nikonorova, N., Van den Broeck, L., Zhu, S., van de Cotte, B., Dubois, M., … & De Smet, I. (2018). Early mannitol-triggered changes in the Arabidopsis leaf (phospho) proteome reveal growth regulators. Journal of Experimental Botany, 69, 4591–4607. https://doi.org/10.1093/jxb/ery261

Nosil, P., Crespi, B. J., & Sandoval, C. P. (2002). Host-plant adaptation drives the parallel evolution of reproductive isolation. Nature, 417, 440–443. https://doi.org/10.1038/417440a

Nosil, P., Funk, D. J., & Ortiz-Barrientos, D. (2009). Divergent selection and heterogeneous genomic divergence. Molecular Ecology, 18, 375–402. https://doi.org/10.1111/j.1365-294X.2008.03946.x

Orsini, L., Vanoverbeke, J., Swillen, I., Mergeay, J., & De Meester, L. (2013). Drivers of population genetic differentiation in the wild: isolation by dispersal limitation, isolation by adaptation and isolation by colonization. Molecular Ecology, 22, 5983–5999. https://doi.org/10.1111/mec.12561

Pickrell, J., & Pritchard J. K. (2012). Inference of population splits and mixtures from genome-wide allele frequency data. PLoS Genetics 8, e1002967. https://doi.org/10.1371/journal.pgen.1002967

Pritchard, J. K., Stephens, M., Rosenberg, N. A., & Donnelly, P. (2000). Association mapping in structured populations. American Journal of Human Genetics, 67, 170–181. https://doi.org/10.1086/302959

Pritchard, J. K., & Rosenberg, N. A. (1999). Use of unlinked genetic markers to detect population stratification in association studies. American Journal of Human Genetics, 65, 220–228. https://doi.org/10.1086/302449

Racimo, F., Berg, F. J., and Pickrel, J. K. (2018). Detecting polygenic adaptation in admixture graphs. Genetics, 208, 1565–1584. https://doi.org/10.1534/genetics.117.300489

Raymond, M., & Rousset, F. (1995). GENEPOP (version 1.2): population genetics software for exact tests and ecumenicism. Journal of Heredity, 86, 248–249.

Rousset, F. (2008). Genepop’007: a complete reimplementation of the Genepop software for Windows and Linux. Molecular Ecology Resources, 8, 103–106. https://doi.org/10.1111/j.1471-8286.2007.01931.x

Santure, A. W., & Garant, D. (2018). Wild GWAS-association mapping in natural populations. Molecular Ecology Resources, 18, 729–738. https://doi.org/10.1111/1755-0998.12901

Togninalli, M., Seren, Ü., Freudenthal, J. A., Monroe, J. G, Meng, D., Nordborg, M., Weigel, D., Borgwardt, K., Korte, A., & Grimm, D. G. (2019). AraPheno and the AraGWAS Catalog 2020: a major database update including RNA-Seq and knockout mutation data for Arabidopsis thaliana. Nucleic Acids Research, 48, D1063–D1068. doi:10.1093/nar/gkz925

Ueda, H. (2019). Sensory mechanisms of natal stream imprinting and homing in Oncorhynchus spp. Journal of Fish Biology, 95, 293–303. https://doi.org/10.1111/jfb.13775

Visscher, P. M., Wray, N. R., Zhang, Q., Sklar, P., McCarthy, M. I., … & Yang, J. (2017). 10 years of GWAS discovery: biology, function, and translation. American Journal of Human Genetics, 101, 5–22. https://doi.org/10.1016/j.ajhg.2017.06.005

Watanabe, K., Stringer, S., Frei, O., Mirkov, M. U., de Leeuw, C., Polderman, T. J. C., van der Sluis, S., Andreassen, O. A., Neale, B. M., & Posthuma, D. (2019). A global overview of pleiotropy and genetic architecture in complex traits. Nature Genetics, 51, 1339–1348. https://doi.org/10.1038/s41588-019-0481-0

Weir, B. S., & Goudet, J. (2017). A unified characterization of population structure and relatedness. Genetics, 206, 2085–2103. https://doi.org/10.1534/genetics.116.198424

Weir, B. S, & Hill, W. G. (2002). Estimating F-statistics. Annual Review of Genetics, 36, 721– https://doi.org/750.10.1146/annurev.genet.36.050802.093940

Wilkinson, S., & Davies, W. J. (2002). ABA-based chemical signalling: the co-ordination of responses to stress in plants. Plant, Cell and Environment, 25, 195–210. https://doi.org/10.1046/j.0016-8025.2001.00824.x

Wilson, P. B., Estavillo, G. M., Field, K. J., Pornsiriwong, W., Carroll, A. J., …. & Pogson, B.J. (2009). The nucleotidase/phosphatase SAL1 is a negative regulator of drought tolerance in Arabidopsis, The Plant Journal, 58, 299–317. https://doi.org/10.1111/j.1365-313X.2008.03780.x

Winter D., Vinegar B., Nahal, H., Ammar, R., Wilson, G. V., & Provart, N. J. (2007). An “electronic fluorescent pictograph” browser for exploring and analyzing large-scale biological data sets. PLoS One 2, e718. https://doi.org/10.1371/journal.pone.0000718

Wright, S. (1931). Evolution in Mendelian population. Genetics, 16, 97–159. PMID: 17246615

Wright, S. (1965). The interpretation of population structure by F-statistics with special regard to systems of mating. Evolution, 19, 395–420. https://www.jstor.org/stable/2406450

Yu, J., Pressoir, G., Briggs, W. H., Bi, I. V., Yamasaki, M., Doebley, J. F., … & Kresovich, S. (2006). A unified mixed-model method for association mapping that accounts for multiple levels of relatedness. Nature Genetics, 38, 203–208. https://doi.org/10.1038/ng1702

Zandalinas, S. I., Balfagón, D., Arbona, V., Gómez-Cadenas, A., Inupakutika, M. A., & Mittler, R. (2016). ABA is required for the accumulation of APX1 and MBF1c during a combination of water deficit and heat stress. Journal of Experimental Botany, 67, erw299. https://doi.org/10.1093/jxb/erw299

Zang, D., Wang, j., Zhang, X., Liu, Z., & Wang, Y. (2019). Arabidopsis heat shock transcription factor HSFA7b positively mediates salt stress tolerance by binding to an E-box-like motif to regulate gene expression. Journal of Experimental Botany, 70, 5355–5374. https://doi.org/10.1093/jxb/erz261

