## Supplementary Figures for "Exploratory analysis of multiple traits co-adaptations in the population history"

### Exploratory analysis of multiple trait coadaptation in the population history

This pdf file includes:
Supplementary Figures S1–S3
Supplementary Tables S1–S5 are presented as an Excel file.

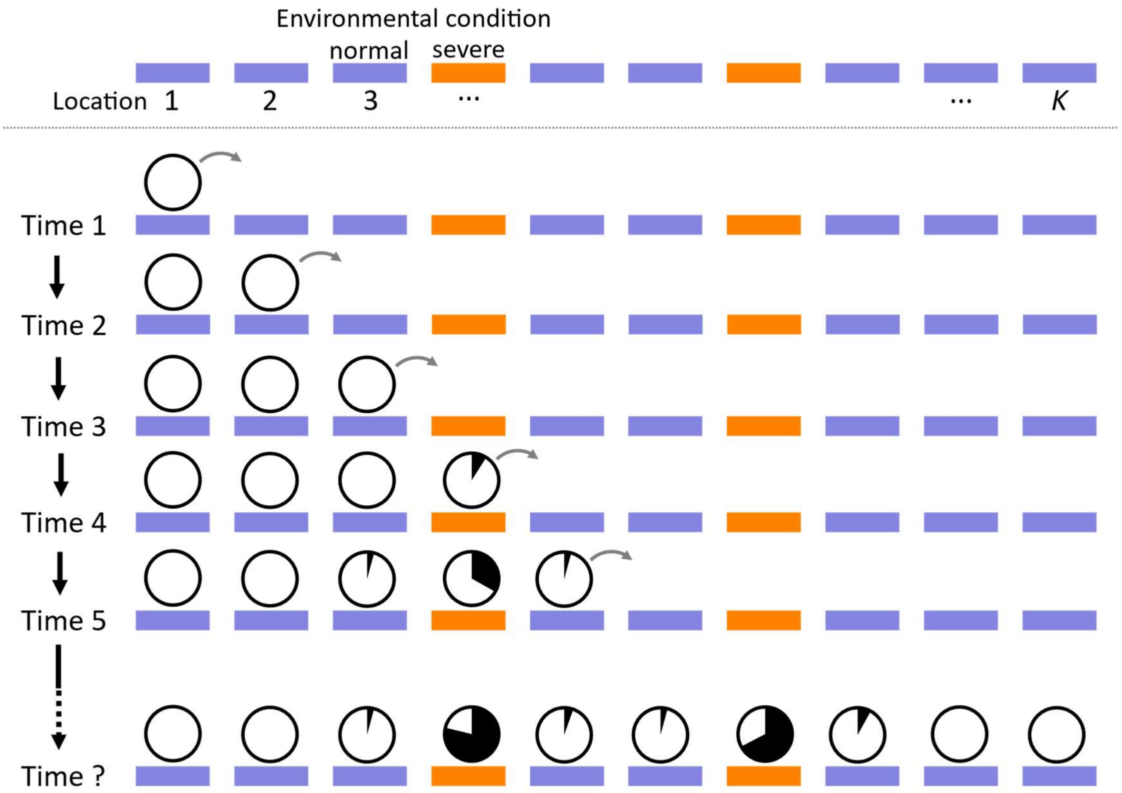


**Figure S1 Population expansion and environmental adaptation.** Each population has a binary environmental condition of *severe* (orange) or *normal* (blue). The population colonization started from population 1 and expanded to population *K*. Pie charts denote frequencies of ancestral (white) and derived (black) alleles. Populations were successively colonized in every 10 generations, and 1% of $N_e$ individuals migrate to adjacent vacant location, as indicated by the arrows (see the text). Migrated individuals increase to $N_e$ in one generation, and thereafter follow the genetic drift.

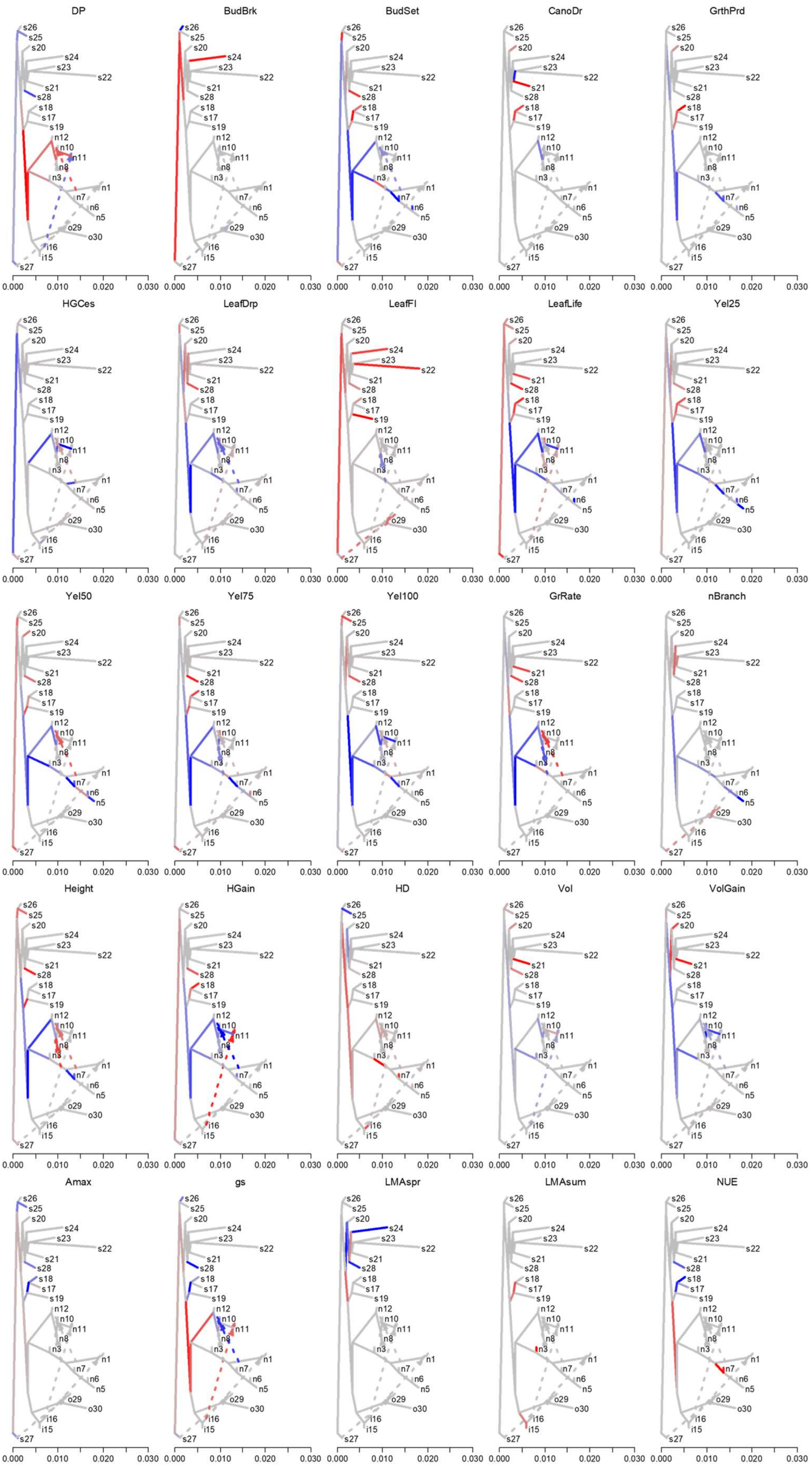
DP
BudBrk
BudSet
CanoDr
GrthPrd
HGCes
LeafDrp
LeafFl
LeafLife
Yel25
Yel50
Yel75
Yel100
GrRate
nBranch
Height
HGain
HD
Vol
VolGain
Amax
gs
LMAspr
LMAsum
NUE
s26
s25
s20
s24
s23
s22
s21
s28
s18
s17
s19
n12
n10
n11
n8
n3
n1
n7
n6
n5
o29
o30
i16
i15
s27
0.000
0.010
0.020
0.030

**Figure S2 Positive selection parameters of 25 traits mapped on admixture graphs.** For each trait, the vector of positive selection parameters that characterize the predicted increase/decrease of the trait value on the admixture graph was estimated using PolyGraph and mapped on the admixture graphs estimated by TREEMIX. Red/blue colors represent selection toward increase/decrease of the trait values.

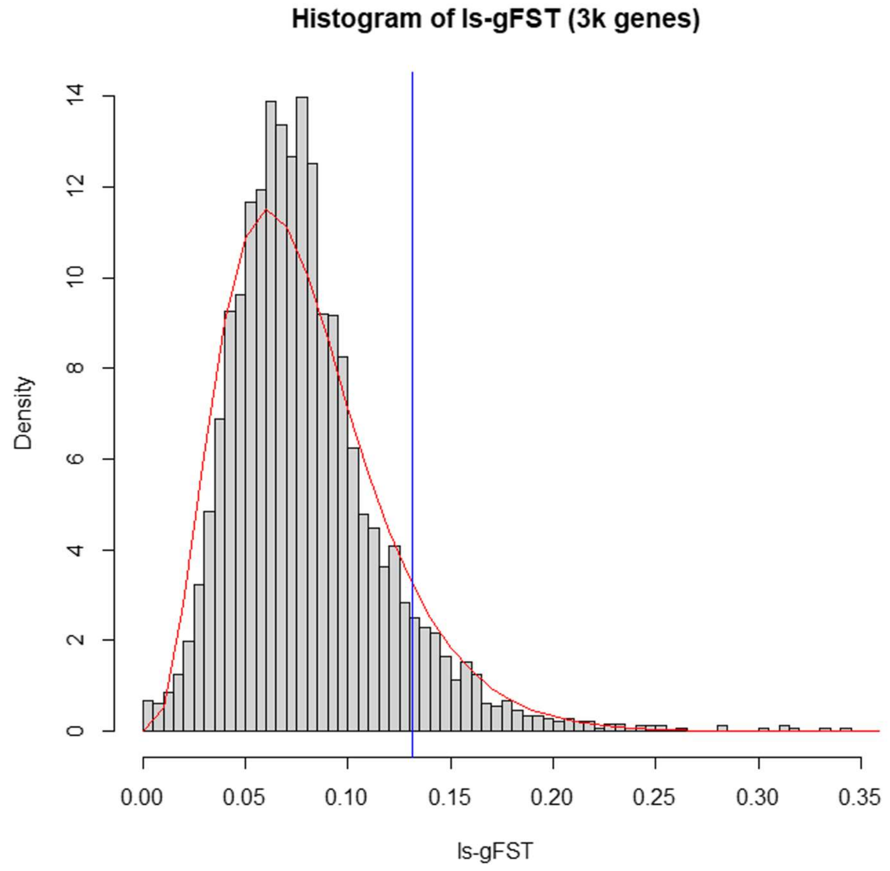


**Figure S3 Distribution of the locus-specific global $F_{ST}$ values and fitted gamma distribution.** Gray histogram is the distribution of locus-specific global $F_{ST}$ values. Red line is the fitted gamma distribution. Blue line is the threshold for upper p-value of 0.1 of the gamma distribution. SNPs in the right side of the blue line were used as adaptive set for association analysis.
