## Supplementary Script for "Exploratory analysis of multiple traits co-adaptations in the population history": admixturegraphFunction20210927.R

```
# Originally implemented by T. Mailund et al. (2016) R package "admixturegraph" (https://cran.r-project.org/package=admixturegraph) 
# Modified by R. Nakamichi

agraph_parents <- function(nodes, parent_edges) {
  n <- length(nodes)
  parents <- matrix(FALSE, n, n)
  rownames(parents) <- colnames(parents) <- nodes
  for (row in 1:nrow(parent_edges)) {
    if (!(parent_edges[row, 1] %in% nodes)) {
      stop(paste("The node", parent_edges[row, 1], 
                 "is used in the edges but is not specified as a node."))
    }
    if (!(parent_edges[row, 2] %in% nodes)) {
      stop(paste("The node", parent_edges[row, 2], 
                 "is used in the edges but is not specified as a node."))
    }
    parents[parent_edges[row, 1], parent_edges[row, 2]] <- TRUE
  }
  parents
}

agraph_children <- function(nodes, parent_edges) {
  n <- length(nodes)
  children <- matrix(FALSE, n, n)
  rownames(children) <- colnames(children) <- nodes
  for (row in 1:nrow(parent_edges)) {
    if (!(parent_edges[row,1] %in% nodes)) {
      stop(paste("The node", parent_edges[row,1], 
                 "is used in the edges but is not specified as a node."))
    }
    if (!(parent_edges[row,2] %in% nodes)) {
      stop(paste("The node", parent_edges[row,2], 
                 "is used in the edges but is not specified as a node."))
    }
    children[parent_edges[row,2], parent_edges[row,1]] <- TRUE
  }
  children
}

agraph_weights <- function(nodes, admixture_weights, parents) {
  n <- length(nodes)
  weights <- matrix("", n, n)
  rownames(weights) <- colnames(weights) <- nodes
  if (!is.null(admixture_weights)) {
    for (row in 1:nrow(admixture_weights)) {
      child <- admixture_weights[row, 1]
      parent <- admixture_weights[row, 2]
      admix_weight <- admixture_weights[row, 3]
      
      if (! parents[child,parent]) {
        stop(paste("There is no edge in the graph from", child, "to", parent))
      }
      
      weights[child, parent] <- admix_weight
      weights[parent, child] <- admix_weight
    }
  }
  weights
}

edge <- function(child, parent) c(child, parent, NA)

admixture_edge <- function(child, parent1, parent2, prop = NA) {
  if (is.na(prop)) {
    other_prop <- NA
  } else {
    other_prop <- paste0("(1 - ", prop, ")")
  }
  c(child, parent1, prop, child, parent2, other_prop)
}

admix_props <- function(child, parent1, parent2, prop)
  c(child, parent1, prop, child, parent2, paste0("(1 - ", prop, ")"))

extract_admixture_proportion_parameters <- function(parent_edges) {
  result <- parent_edges[!is.na(parent_edges[,3]),]
  if(nrow(result) > 0) result else NULL
}

parent_edges <- function(edges) matrix(ncol = 3, byrow = TRUE, data = edges)

admixture_proportions <- function(admix_props) matrix(ncol = 3, byrow = TRUE, data = admix_props)

agraph <- function(leaves, inner_nodes, parent_edges, 
                   admixture_proportions = extract_admixture_proportion_parameters(parent_edges)) {
  nodes <- c(leaves, inner_nodes)
  parents <- agraph_parents(nodes, parent_edges)
  children <- agraph_children(nodes, parent_edges)
  admixture_probs <- agraph_weights(nodes, admixture_proportions, parents)
  structure(list(leaves = leaves, inner_nodes = inner_nodes,
                 nodes = nodes,
                 parents = parents,
                 probs = admixture_probs,
                 children = children),
            class = "agraph")
}
```
