## Supplementary Script for "Exploratory analysis of multiple traits co-adaptations in the population history": FinePop2Ext20211002.R

```
GLS <- function(model, data, omega=NULL){
  n <- nrow(data)
  model <- terms(model, data=data)
  y.name <- as.character(model[[2]])
  x.name <- attr(model,"term")

  if(is.null(omega)){omega <- diag(nrow(data))}

  y <- data[, y.name, drop=FALSE]
  x <- data[, x.name, drop=FALSE]
  ydev <- as.matrix(y - colMeans(y))
  xdev <- as.matrix(x - colMeans(x))
  #xdev <- as.matrix(apply(x, 2, function(x){x-mean(x)}))

  numP <- ncol(x)

  somega <- solve(omega)
  b_hat <- solve((t(xdev) %*% somega %*% xdev)) %*% (t(xdev) %*% somega %*% ydev)  
  resid0 <- ydev - xdev %*% b_hat
  sigma2 <- as.numeric(1/(n-numP-1) * (t(resid0) %*% somega %*% resid0))
  var_b <- sigma2 * solve(t(xdev) %*% somega %*% xdev)

  se <- sqrt(diag(var_b))
  zv <- t(b_hat) / se
  pvz <- pnorm(abs(zv), lower.tail=FALSE)*2 
  #tv <- zv * sqrt((n-1)/n)
  #pvt <- pt(abs(tv), df=n-1, lower.tail=FALSE)*2 
  
  logL <- -n/2 * log(sigma2) - 1/2*log(det(omega)) - n/2

  #tv <- as.vector(tv) 
  #pvt <- as.vector(pvt)
  zv <- as.vector(zv)
  pvz <- as.vector(pvz)
  b_hat <- as.vector(b_hat) 
  names(b_hat) <- x.name
  return(list(
    coefficients=data.frame(
      estimate=b_hat,
      std.error=se,
      #t.value=tv,
      #p.value.t=pvt,
      Z.value=zv,
      p.value=pvz
    ),
    variance=var_b,
    logL=logL
  ))
}


minor.allele.freq <- function(popdata){
  maf <- array(NA, dim=c(popdata$num_pop, popdata$num_loci),
               dimnames=list(popdata$pop_names, popdata$loci_names))
  for(loci.id in 1:popdata$num_loci){
    caf <- popdata$allele_freq[[loci.id]][1,]
    caf <- if(mean(caf,na.rm=T)<0.5){caf}else{1-caf}
    maf[,loci.id] <- caf
  }
  return(maf)
}
minor.allele.count <- function(popdata){
  mac <- array(NA, dim=c(popdata$num_pop, popdata$num_loci),
               dimnames=list(popdata$pop_names, popdata$loci_names))
  for(loci.id in 1:popdata$num_loci){
    cac <- popdata$allele_count[[loci.id]][1,]
    cind <- popdata$ind_count[[loci.id]]
    cac <- if(sum(cac,na.rm=T)<sum(cind,na.rm=T)){cac}else{cind*2-cac}
    mac[,loci.id] <- cac
  }
  return(mac)
}

GET.data <- function(popdata, environment, trait=NULL){
  maf <- minor.allele.freq(popdata)
  na.check <- !is.na(colSums(maf))
  maf <- maf[,na.check]

  if(is.null(colnames(environment))){colnames(environment) <- paste0("env",1:ncol(environment))}
  if((!is.null(trait))&is.null(colnames(trait))){colnames(trait) <- paste0("trait",1:ncol(trait))}

  get.data <- cbind(environment, trait, maf)
  rownames(get.data) <- popdata$pop_names

  get.index <- c(rep("environment", ncol(environment)),
                 rep("trait", ncol(trait)),
                 rep("gene", ncol(maf)))
  get.index <- factor(get.index, levels=c("environment","trait","gene"))

  return(list(data=get.data, index=get.index))
}

GET.data.attribute <- function(data.get, label=NULL,
                               col.env="darkgreen", col.trait="orange", col.gene="black",
                               size.env=1.5, size.trait=1.5, size.gene=0.7){
  col.data <- c(col.env,col.trait,col.gene)
  names(col.data) <- c("environment","trait","gene")

  size.data <- c(size.env, size.trait, size.gene)
  names(size.data) <- c("environment","trait","gene")

  if(is.null(label)){label <- colnames(data.get$data)}

  index <- data.get$index
  attr <- data.frame(label=label, type=data.get$index,
                     col=col.data[index], size=size.data[index])
  rownames(attr) <- colnames(data.get$data)
  return(attr)
}


GET.GLS <- function(data.get, omega.list=NULL){
  env <- data.get$data[,data.get$index=="environment", drop=F]
  trait <- data.get$data[,data.get$index=="trait", drop=F]
  gene <- data.get$data[,data.get$index=="gene", drop=F]

  gt <- scale(cbind(trait,gene))
  if.gene <- c(rep(F, ncol(trait)), rep(T,ncol(gene)))
  env <- scale(env)

  n.e <- ncol(env)
  n.gt <- ncol(gt)
  n.p <- nrow(env)
 
  num.edge <- n.e*n.gt
  zvec <- data.frame(environment=rep(colnames(env),each=n.gt),
                     trait_gene=rep(colnames(gt), n.e), z=rep(0, num.edge), p=rep(0, num.edge))
  Imat <- diag(n.p)
  omega <- Imat
  cedge <- 1
  for(ce in 1:n.e){
    ctext <- paste0(cedge, "/", num.edge)
    message(ctext, appendLF=F);flush.console()
    x <- env[,ce]
    x <- x - mean(x) 
  for(cg in 1:n.gt){
    y.name <- colnames(gt)[cg]
    y <- gt[,cg]
    y <- y - mean(y)
    Z <- NA; pv <- NA
    comega <- if(if.gene[cg]){omega.list[,,y.name]}else{Imat}
    resGLS <- GLS(y~x, data.frame(y,x), comega)
    Z <- resGLS$coefficients["x","Z.value"]
    pv <- resGLS$coefficients["x","p.value"]
    zvec[cedge,"z"] <- Z
    zvec[cedge,"p"] <- pv
    cedge <- cedge+1
  }
    message(rep("\b",nchar(ctext)), appendLF=F)
  }
  rownames(zvec) <- NULL
  
  return(zvec)
}

GET.FDR <- function(result.gls, fdr){
  env <- unique(as.character(unlist(result.gls[,"environment"])))
  edge.signif <- rep(F, nrow(result.gls))
  for(cenv in env){
    cedge <- which(result.gls[,"environment"]==cenv) 
    cp <- result.gls[cedge,"p"]
    cq <- p.adjust(cp, method="BH")
    cedge.i <- cq < fdr
    edge.signif[cedge] <- cedge.i
  }
  edge.signif <- which(edge.signif)
  result.fdr <- result.gls[edge.signif,]
  rownames(result.fdr) <- NULL
  return(result.fdr)
}

GET.FDR.node <- function(result.fdr){
  unique(as.character(unlist(result.fdr[,c("environment","trait_gene")])))
}

GET.corresp <- function(data.get){
  env <- data.get$data[,data.get$index=="environment", drop=F]
  trait <- data.get$data[,data.get$index=="trait", drop=F]
  gene <- data.get$data[,data.get$index=="gene", drop=F]

  nenv <- -env
  colnames(nenv) <- paste0("-",colnames(nenv))
  env <- apply(cbind(env,nenv), 2, function(x){(x-min(x))/(max(x)-min(x))})
  rm(nenv)

  trait <- apply(trait, 2, function(x){(x-min(x))/(max(x)-min(x))})

  data.crp <- cbind(env,trait,gene)
  res.crp <- corresp(data.crp, nf=2)
  return(res.crp)
}

plot.GET.corresp <- function(result.crp, result.fdr, data.get.attr, pop.label=NULL, pop.color=NULL){
  lim.xy <- rbind(result.crp$rscore,result.crp$cscore)
  lim.xy <- rbind(apply(lim.xy, 2, min),apply(lim.xy, 2, max))
  lim.xy <- lim.xy*1.1
  dimnames(lim.xy) <- list(c("min","max"),c("x","y"))

  node.fdr <- unique(as.character(unlist(result.fdr[,c("environment","trait_gene")])))
  #if(is.null(node.label)){node.label <- node.fdr}

  attr <- data.get.attr[node.fdr,]
  node.fdr.env <- unique(as.character(result.fdr[,c("environment")]))
  node.fdr.gt <- unique(as.character(result.fdr[,c("trait_gene")]))
  attr.e <- attr[node.fdr.env,]
  attr.ne <- attr.e
  attr.e[,"label"] <- paste0("+", attr.e[,"label"])
  rownames(attr.ne) <- paste0("-",rownames(attr.ne))
  attr.ne[,"label"] <- paste0("-", attr.ne[,"label"])
  attr.gt <- attr[node.fdr.gt,]
  attr <- rbind(attr.e, attr.ne, attr.gt)
  font.e <- rep(2, nrow(attr.e))
  font.ne <- rep(4, nrow(attr.ne))
  font.gt <- rep(3, nrow(attr.gt))
  font.gt[attr.gt[,"type"]=="trait"] <- 2
  attr <- cbind(attr, font=c(font.e, font.ne, font.gt))
  node.fdr <- rownames(attr)

  plot(result.crp$cscore, xlim=lim.xy[,"x"], ylim=lim.xy[,"y"], col="gray", pch=19,
       cex=min(attr[,"size"]), xlab="1st axis",ylab="2nd axis")

  node.draw <- NULL
  for(cedge.i in 1:nrow(result.fdr)){
    cedge <- result.fdr[cedge.i,]
    cgt <- as.character(cedge[,"trait_gene"])
    cenv <- as.character(cedge[,"environment"])
    if(cedge[,"z"]<0){cenv <- paste0("-",cenv)}
    cxy <- result.crp$cscore[c(cenv,cgt),]
    lines(x=cxy[,1], y=cxy[,2], col="gray")
    node.draw <- c(node.draw, cgt, cenv)
  }
  node.draw <- unique(node.draw)

  points(0,0, pch=3, cex=5)
  points(result.crp$cscore[node.draw,], col="gray", #col=as.character(attr[node.draw,"col"]),
         pch=19)#, cex=attr[node.draw,"size"])

  text(result.crp$cscore[node.draw,], labels=as.character(attr[node.draw,"label"]),
       col=as.character(attr[node.draw,"col"]), font=as.integer(attr[node.draw,"font"]))

  if(is.null(pop.label)){pop.label <- rownames(result.crp$rscore)}
  if(is.null(pop.color)){pop.color <- "red"}
  text(result.crp$rscore, labels=pop.label, col=pop.color)
}

plot.GET.correspET <- function(result.crp, result.fdr, data.get.attr, pop.label=NULL, pop.color=NULL,
                               lim.xy=NULL, cex.pop=1, orig.list=NULL){
  if(is.null(lim.xy)){
    lim.xy <- rbind(result.crp$rscore,result.crp$cscore)
    lim.xy <- rbind(apply(lim.xy, 2, min),apply(lim.xy, 2, max))
    lim.xy <- lim.xy*1.1
    dimnames(lim.xy) <- list(c("min","max"),c("x","y"))
  }
  attr.e <- data.get.attr[data.get.attr[,"type"]=="environment",]
  attr.ne <- attr.e
  attr.e[,"label"] <- paste0("+", attr.e[,"label"])
  rownames(attr.ne) <- paste0("-",rownames(attr.ne))
  attr.ne[,"label"] <- paste0("-", attr.ne[,"label"])
  attr.t <- data.get.attr[data.get.attr[,"type"]=="trait",]
  attr <- rbind(attr.e, attr.ne, attr.t)
  font.e <- rep(1, nrow(attr.e))
  font.ne <- rep(3, nrow(attr.ne))
  font.t <- rep(1, nrow(attr.t))
  attr <- cbind(attr, font=c(font.e, font.ne, font.t))

  result.fdr.use <- NULL
  for(corig in orig.list){
    csign <- substr(corig, 1,1)
    cenv <- substr(corig, 2,nchar(corig));if(cenv=="ALT"){cenv<-"alt"}
    cfdr <- result.fdr[result.fdr[,"environment"]==cenv,]
    cfdr.use <- cfdr[,"z"]>=0
    if(csign=="-"){cfdr.use <- !cfdr.use}
    cfdr <- cfdr[cfdr.use,,drop=F]
    result.fdr.use <- rbind(result.fdr.use, cfdr)
  }
  attr.g.use <- data.get.attr[result.fdr.use[,"trait_gene"],]
  font.g.use <- rep(2, nrow(attr.g.use))
  attr <- rbind(attr, cbind(attr.g.use, font=font.g.use))

  plot(result.crp$cscore, xlim=lim.xy[,"x"], ylim=lim.xy[,"y"], col="gray", pch=19,
       cex=min(data.get.attr[,"size"]), xlab="1st axis",ylab="2nd axis")

  node.draw <- NULL
  if(!is.null(orig.list)){
    for(cedge.i in 1:nrow(result.fdr.use)){
      cedge <- result.fdr.use[cedge.i,]
      cgt <- as.character(cedge[,"trait_gene"])
      cenv <- as.character(cedge[,"environment"])
      if(cedge[,"z"]<0){cenv <- paste0("-",cenv)}
      cxy <- result.crp$cscore[c(cenv,cgt),]
      lines(x=cxy[,1], y=cxy[,2], col="gray")
      node.draw <- c(node.draw, cgt, cenv)
    }
  }
  node.draw <- unique(c(node.draw, rownames(attr)))

  points(0,0, pch=3, cex=5)

  text(result.crp$cscore[node.draw,],
       labels=as.character(attr[node.draw,"label"]), cex=as.numeric(attr[node.draw,"size"]),
       col=as.character(attr[node.draw,"col"]), font=as.integer(attr[node.draw,"font"]))

  if(is.null(pop.label)){pop.label <- rownames(result.crp$rscore)}
  if(is.null(pop.color)){pop.color <- "red"}
  text(result.crp$rscore, labels=pop.label, col=pop.color, cex=cex.pop)
}


cov.pop <- function(popdata){
  n.pop <- popdata$num_pop
  Cmaf.lp <- t(minor.allele.freq(popdata))
  Mmaf.l <- rowMeans(Cmaf.lp,na.rm=T)
  CmafS.lp <- Cmaf.lp/sqrt(Mmaf.l*(1-Mmaf.l))

  cov.mafMoment <- array(NA, c(n.pop,n.pop))
  for(cp1 in 1:(n.pop-1)){
  for(cp2 in (cp1+1):n.pop){
    cmafs1 <- CmafS.lp[,cp1]
    cmafs2 <- CmafS.lp[,cp2]
    cmafs12 <- cmafs1*cmafs2
    cnl <- sum(!is.na(cmafs12))
    cov.mafMoment[cp1,cp2] <- cov.mafMoment[cp2,cp1] <- sum(cmafs12, na.rm=T) / cnl
  }}
  for(cp in 1:n.pop){
    cmafs <- CmafS.lp[,cp]
    cmaf <- Cmaf.lp[,cp]
    cnl <- sum(!is.na(cmafs))
    cn <- popdata$pop_sizes[cp] * 2
    t1 <- 1/cnl * sum(cmafs^2, na.rm=T)
    t2 <- 1/cn * 1/cnl * sum( (cmaf*(1-cmaf))/(Mmaf.l*(1-Mmaf.l)) ,na.rm=T)
    #message("t1=",t1, " t2=",t2)
    cov.mafMoment[cp,cp] <- t1 - t2
  }

  return(cov.mafMoment)
}


cov.pop.maf <- function(popdata){
  n.pop <- popdata$num_pop
  n.loci <- popdata$num_loci
  n.ind.lp <- matrix(unlist(popdata$ind_count), byrow=T, ncol=n.pop)
  n.ind.p <- popdata$pop_size
  N <- sum(n.ind.p) * 2
  Cmaf.lp <- t(minor.allele.freq(popdata))
  Cmac.l <- colSums(minor.allele.count(popdata)) 
  #Mmaf.l <- rowMeans(Cmaf.lp,na.rm=T)
  Mmaf.l <- Cmac.l / N

  MmafS.l <- 1/(Mmaf.l*(1-Mmaf.l))
  cov.pop <- array(NA, c(n.pop,n.pop), dimnames=list(popdata$pop_names, popdata$pop_names))
  var.maf <- array(NA, c(n.pop,n.loci), dimnames=list(popdata$pop_names, popdata$loci_names))
  for(cp1 in 1:(n.pop-1)){
  for(cp2 in (cp1+1):n.pop){
    cov12 <- Cmaf.lp[,cp1] * Cmaf.lp[,cp2]
    cov.pop[cp1,cp2] <- cov.pop[cp2,cp1] <- mean(MmafS.l * cov12, na.rm=T)
  }}
  for(cp in 1:n.pop){
    cmaf <- Cmaf.lp[,cp]
    cn <- n.ind.lp[,cp] * 2
    ppn <- cmaf * (1-cmaf) / cn
    v <- cmaf^2
    cov.pop[cp,cp] <- mean(MmafS.l * (v - ppn) , na.rm=T)
    var.maf[cp,] <- ppn
  }

  cov.maf <- array(NA, c(n.pop,n.pop,n.loci), dimnames=list(popdata$pop_names, popdata$pop_names,popdata$loci_names))
  for(cl in 1:n.loci){
    ccov <- cov.pop * Mmaf.l[cl] * (1-Mmaf.l[cl])
    diag(ccov) <- diag(ccov) + var.maf[,cl]
    cov.maf[,,cl] <- ccov
  }

  return(list(cov.pop=cov.pop, cov.maf=cov.maf))
}

cov.pop.mafW <- function(popdata, af.cut=NULL, af.depth=NULL){
  n.pop <- popdata$num_pop
  n.loci <- popdata$num_loci
  n.ind.lp <- matrix(unlist(popdata$ind_count), byrow=T, ncol=n.pop)
  n.ind.p <- popdata$pop_size
  Cmaf.lp <- t(minor.allele.freq(popdata))
  Cmac.l <- colSums(minor.allele.count(popdata)) 
  Mmaf.l <- rowMeans(Cmaf.lp,na.rm=T)

  n <- sum(n.ind.p) * 2
  pa.l <- NULL
  w.l <- NULL
  if(is.null(af.cut)|is.null(af.depth)){
    pa.l <- Cmac.l / n 
    w.l <- rep(1, n.loci)
  }else{
    d <- af.depth * 2
    C <- af.depth * 2 * af.cut
    pASC.i <- array(NA, n-1)
    n.i <- array(NA, n-1)
    for(i in 1:(n-1)){
      cnum.i <- 0
      for(j in as.integer(C):as.integer(n-C)){cnum.i <- cnum.i + choose(i,j) * choose(n-i,d-j)}
      pASC.i[i] <- cnum.i / choose(n,d)
      n.i[i] <- sum(Cmac.l==i) ##### �����炪�{��
      #num.pk.i[i] <- n.i[i] / pASC.i[i]
    }
    num.pk.i <- n.i / pASC.i

    pk <- num.pk.i / sum(num.pk.i, na.rm=T)
    w.l <- array(NA, n.loci)
    pa.l <- array(NA, n.loci)
    for(cl in 1:n.loci){
      cmac <- as.integer(Cmac.l[cl])
      w.l <- 1 / pASC.i[cmac]
      pa.l[cl] <- sum( pk * ((1:(n-1)/n)^(cmac+1)) * ((1-1:(n-1)/n)^(n-cmac)) ,na.rm=T) / 
                  sum( pk * ((1:(n-1)/n)^cmac) * ((1-1:(n-1)/n)^(n-cmac)) ,na.rm=T)
    }
  }

  #MmafS.l <- 1/(Mmaf.l*(1-Mmaf.l))
  AmafS.l <- w.l / (pa.l*(1-pa.l))
  cov.pop <- array(NA, c(n.pop,n.pop), dimnames=list(popdata$pop_names, popdata$pop_names))
  var.maf <- array(NA, c(n.pop,n.loci), dimnames=list(popdata$pop_names, popdata$loci_names))
  for(cp1 in 1:(n.pop-1)){
  for(cp2 in (cp1+1):n.pop){
    cov12 <- (Cmaf.lp[,cp1] - pa.l) * (Cmaf.lp[,cp2] - pa.l)
    #cov12 <- Cmaf.lp[,cp1] * Cmaf.lp[,cp2]
    cov.pop[cp1,cp2] <- cov.pop[cp2,cp1] <- sum(AmafS.l * cov12, na.rm=T) / sum(w.l,na.rm=T)
  }}
  for(cp in 1:n.pop){
    cmaf <- Cmaf.lp[,cp]
    cn <- n.ind.lp[,cp] * 2
    ppn <- cmaf * (1-cmaf) / cn
    v <- (cmaf - pa.l)^2
    #v <- cmaf^2
    cov.pop[cp,cp] <- sum(AmafS.l * (v - ppn), na.rm=T) / sum(w.l,na.rm=T)
    var.maf[cp,] <- ppn
  }

  cov.maf <- array(NA, c(n.pop,n.pop,n.loci),dimnames=list(popdata$pop_names, popdata$pop_names,popdata$loci_names))
  for(cl in 1:n.loci){
    ccov <- cov.pop * pa.l[cl] * (1-pa.l[cl])
    diag(ccov) <- diag(ccov) + var.maf[,cl]
    cov.maf[,,cl] <- ccov
  }

  return(list(cov.pop=cov.pop, cov.maf=cov.maf))
}

cov.pop.mafW2 <- function(popdata, locus=F){
  n.pop <- popdata$num_pop
  n.loci <- popdata$num_loci
  n.ind.lp <- matrix(unlist(popdata$ind_count), byrow=T, ncol=n.pop)
  n.ind.p <- popdata$pop_size
  Cmaf.lp <- t(minor.allele.freq(popdata))
  Cmac.l <- colSums(minor.allele.count(popdata)) 
  Mmaf.l <- rowMeans(Cmaf.lp,na.rm=T)

  n <- sum(n.ind.p) * 2
  n.p <- n.ind.p * 2

  pa.l <- Cmac.l / n
  w.l <- rep(1, n.loci)

  AmafS.l <- w.l / (pa.l*(1-pa.l))
  cov.pop <- array(NA, c(n.pop,n.pop), dimnames=list(popdata$pop_names, popdata$pop_names))
  var.maf <- array(NA, c(n.pop,n.loci), dimnames=list(popdata$pop_names, popdata$loci_names))
  for(cp1 in 1:(n.pop-1)){
  for(cp2 in (cp1+1):n.pop){
    #cov12 <- (Cmaf.lp[,cp1] - pa.l) * (Cmaf.lp[,cp2] - pa.l)
    cov12 <- Cmaf.lp[,cp1] * Cmaf.lp[,cp2]
    cov.pop[cp1,cp2] <- cov.pop[cp2,cp1] <- sum(AmafS.l * cov12, na.rm=T) / sum(w.l,na.rm=T)
  }}
  for(cp in 1:n.pop){
    cmaf <- Cmaf.lp[,cp]
    cn <- n.ind.lp[,cp] * 2
    #v <- (cmaf - pa.l)^2
    v <- cmaf^2
    cov.pop[cp,cp] <- 
       sum( w.l * (v/(pa.l*(1-pa.l)) - 1/cn) / (1-1/cn) ,na.rm=T) / sum(w.l,na.rm=T)
    var.maf[cp,] <- cmaf * (1-cmaf) / cn
  }
  ppn <- pa.l * (1-pa.l) / n

  cov.maf <- array(NA, c(n.pop,n.pop,n.loci),dimnames=list(popdata$pop_names, popdata$pop_names,popdata$loci_names))
  for(cl in 1:n.loci){
    ccov <- cov.pop * pa.l[cl] * (1-pa.l[cl])
    #diag(ccov) <- diag(ccov) + var.maf[,cl]
    diag(ccov) <- diag(ccov) + ppn[cl]
    cov.maf[,,cl] <- ccov
  }

  output <- list(cov.pop=cov.pop, cov.maf=cov.maf)
  class(output) <- "cov.loci"
  return(output)
}


cov.pop.mafW3 <- function(popdata, outlier.threshold=NULL){
  n.pop <- popdata$num_pop
  n.loci <- popdata$num_loci
  n.ind.pl <- matrix(unlist(popdata$ind_count), byrow=F, nrow=n.pop)
  n.ind.p <- popdata$pop_size
  Cmaf.pl <- minor.allele.freq(popdata)
  Cmac.l <- colSums(minor.allele.count(popdata)) 
  Mmaf.l <- colMeans(Cmaf.pl,na.rm=T)

  #Cmaf.pl <- apply(Cmaf.pl, 2, function(x){(x-min(x))/(max(x)-min(x))})

  if(is.null(outlier.threshold)){outlier.threshold <- 1/(popdata$num_loci*10)}

  gFST <- suppressMessages(locus_specificFST(popdata))

  #k <- n.pop-1
  #med.chi <- k - 2/3 + 4/(27*k) - 8/(729*k^2)
  #phai.median <- median(gFST) / med.chi
  #theta_hat_median <- gFST / phai.median
  #p <- pchisq(theta_hat_median, df=k, lower.tail=F)
  #q <- p.adjust(p, "BH")
  #order.p <- order(p)
  #loci.non.outlier <- p > outlier.threshold
  #loci.non.outlier <- q > outlier.threshold
  #loci.non.outlier <- order.p > outlier.threshold*n.loci

  gammafit.gfst <- suppressWarnings(fitdistr(x=gFST, densfun="gamma"))
  gs <- gammafit.gfst$estimate["shape"]
  gr <- gammafit.gfst$estimate["rate"]
  q.threshold <- qgamma(outlier.threshold, shape=gs, rate=gr, lower.tail=F)
  loci.non.outlier <- gFST < q.threshold

  n.loci.no <- sum(loci.non.outlier)
  message(n.loci.no, "/", n.loci)

  n <- sum(n.ind.p) * 2
  n.p <- n.ind.p * 2

  #pa.l <- Cmac.l[loci.non.outlier] / n
  pa.l <- Mmaf.l[loci.non.outlier]
  AmafS.l <- 1 / (pa.l*(1-pa.l))
  cov.pop <- array(NA, c(n.pop,n.pop), dimnames=list(popdata$pop_names, popdata$pop_names))
  for(cp1 in 1:(n.pop-1)){
  for(cp2 in (cp1+1):n.pop){
    cmaf1 <- Cmaf.pl[cp1,loci.non.outlier]
    cmaf2 <- Cmaf.pl[cp2,loci.non.outlier]
    cov12 <- cmaf1 * cmaf2
    cov.pop[cp1,cp2] <- cov.pop[cp2,cp1] <- mean(AmafS.l * cov12, na.rm=T)
  }}
  for(cp in 1:n.pop){
    cmaf <- Cmaf.pl[cp,loci.non.outlier]
    cn <- n.ind.pl[cp,loci.non.outlier] * 2
    v <- cmaf^2
    cov.pop[cp,cp] <- mean( (v/(pa.l*(1-pa.l)) - 1/cn) / (1-1/cn) ,na.rm=T)
  }

  #pa.l <- Cmac.l / n
  pa.l <- Mmaf.l
  var.maf <- array(NA, c(n.pop,n.loci), dimnames=list(popdata$pop_names, popdata$loci_names))
  for(cp in 1:n.pop){
    cn <- n.ind.pl[cp,] * 2
    #cmaf <- Cmaf.pl[cp,]
    #var.maf[cp,] <- cmaf * (1-cmaf) / cn
    var.maf[cp,] <- pa.l * (1-pa.l) / cn
    #var.maf[cp,] <- ( pa.l - pa.l^2 - pa.l*(1-pa.l)*cov.pop[cp,cp] ) / cn
  }
  cov.maf <- array(NA, c(n.pop,n.pop,n.loci),dimnames=list(popdata$pop_names, popdata$pop_names,popdata$loci_names))
  for(cl in 1:n.loci){
    ccov <- cov.pop * pa.l[cl] * (1-pa.l[cl])
    diag(ccov) <- diag(ccov) + var.maf[,cl]
    cov.maf[,,cl] <- ccov
  }

  output <- list(cov.pop=cov.pop, cov.maf=cov.maf)
  class(output) <- "cov.loci"
  return(output)
}
```
