## Supplementary Script for "Exploratory analysis of multiple traits co-adaptations in the population history": OmegaFunction20210927.R

```
# Originally created by Yamin Deng, Tao He, and Yuehua Cui et al. (2020) Frontiers in Genetics 11:437
# Modified by R. Nakamichi

######Gene-based analysis for single trait analysis#############

##########function to generate kernel matrix######
#Note:for demonstration reasons, only certain commonly used kernels are defined in the function,
#such as Gaussion/Quadratic/Linear/IBS. 
#The users can define the candidate kernel set of their interests.

gen.ker <- function(covx,kernel.index)
#covx:X; 
#kernel.index=c("Gau","Lin","Quad","IBS"): index of kernel function;
{ 
  n <- nrow(covx)
  p <- ncol(covx)

  if (kernel.index=="Gau"){
    ker <- gausskernel(X=covx,sigma=p) # Gaussian kernel
  }
  if (kernel.index=="Lin"){
    f <- function(x,y) x%*%y # linear kernel
    ker <- outer(1:n, 1:n, Vectorize( function(i,j) f(covx[i,], covx[j,]) ) )
  }
  if (kernel.index=="Quad"){
    f <- function(x,y) (x%*%y+1)^2 # Quadratic kernel
    ker <- outer(1:n, 1:n, Vectorize( function(i,j) f(covx[i,], covx[j,]) ) )
  }
  if (kernel.index=="IBS"){
    f <- function(x,y) 1-sum(abs(x-y))/(2*p) #IBS kernel
    ker <- outer(1:n, 1:n, Vectorize( function(i,j) f(covx[i,], covx[j,]) ) )
  } 
  ker <- as.matrix(forceSymmetric(ker))
  ker0 <- ker
  diag(ker0) <- rep(0,n)
  J <- matrix(1,n,n)
  ker.cen <- ker-J%*%ker0/(n-1)-ker0%*%J/(n-1)+J%*%ker0%*%J/n/(n-1)
  v1.cen <- tr(ker.cen)/n
  return(list(ker.cen=ker.cen,v1.cen=v1.cen))
}

Chisq.Ustat<-function(y,covx,z,ker)
{
  n=length(y)
  p=ncol(covx)
  re.norm=re.lin=re.u=0
  vec1=rep(1,n)
  h=diag(n)-z%*%solve(t(z)%*%z)%*%t(z)
  ker1=ker
  diag(ker1)=rep(0,n)
  ############################################################################
  
  A=h%*%ker1%*%h
  hkhk=A%*%ker1
  hkh.hkh=A*A
  hk=h%*%ker1
  sigma.hat=t(y)%*%h%*%y/(n-ncol(z))
  sd=as.numeric(sqrt(sigma.hat))
  z0=h%*%y/sd
  m4=mean(z0^4)
  
  e1=tr(hk)
  e2=tr(hkhk)
  mu1=2*e1
  mu2=e1*(1+2/(n-1))
  delta=m4-3

  var0=e1^2*(-2/(n-1))+e2*(2-12/(n-1)) #high order terms---simulation 1
  var=var0+delta*(-e1^2/n+6*e2/n+tr(hkh.hkh)) #high order terms---simulation 1
  #print(var)
  test=t(h%*%y)%*%ker1%*%(h%*%y)/(sigma.hat)
  
  v1=sum(diag(ker))/n
  theta=-sum(ker1)/n/(n-1)
  theta=0  #if using centralized kernel
  
  a=var/2/(n-1)^2/v1
  g=v1/a
  
  if (test>qnorm(0.95,theta*(n-1),sqrt(var))) 
    re.norm=1 
  if ((test/(n-1)+v1-theta)>a*qchisq(0.95,g)) 
    re.u=1
  sd.chi=test/(n-1)+v1-theta
  p.u=pchisq(sd.chi/a,g,lower.tail = F)
  test.sd=(test-theta*(n-1))/sqrt(var)

  #return(list(re.norm=re.norm, re.u=re.u,test.sd=test.sd,A.ma=h%*%ker1%*%h/sqrt(var),p.u=p.u))  
  return(p.u)
}


#######################Fisher combination with dependent traits####################
FisherCombPval <- function(P.values, Pheno,
  kendall.only=FALSE,miss.pval=FALSE,
  verbose=FALSE){
  
  if(!is.data.frame(Pheno)) Pheno <- as.data.frame(Pheno)
  if(is.data.frame(P.values)) P.values <- as.data.frame(P.values)
  if(ncol(Pheno) != ncol(P.values))
  stop("The number of phenotypes does not match the number of p-values")
  log.P.values <- log(P.values)
  G <- nrow(P.values)
  res <- rep(NA, G)
  
  PAR <- cal.gamma.par(Pheno,kendall.only,verbose)
  SHAPE <- PAR$SHAPE
  SCALE <- PAR$SCALE
  
  T <- -2 * apply(log.P.values,1,sum)
  res <- pgamma(T, shape=SHAPE, scale=SCALE, lower.tail=FALSE)
  
  if(miss.pval && is.na(res)){
    idx <- which(is.na(res))
    for (i in idx){
    ##cat("i = ",i,"\n")
      x <- log.P.values[i,]
      y <- x[!is.na(x)]
      if(length(y) == 1){
        res[i] <- exp(y)
      }else{
        T <- -2*sum(y)
        PAR <- cal.gamma.par(Pheno[,!is.na(x)],kendall.only,verbose)
        SHAPE <- PAR$SHAPE
        SCALE <- PAR$SCALE
        res[i] <- pgamma(T, shape=SHAPE, scale=SCALE, lower.tail=FALSE)
      }
    }
  }
  return(res)
}

### utilities
adj.cor <- function(r,n){
  fac <- (1-r^2) / (2*(n-3))
  res <- r * (1 + fac)
  return(res)
}

cal.cor.pearson <- function(x,y){
  r <- cor(x,y) #,use="pairwise.complete.obs")
  return(r)
}

### internal functions
cal.cor.kendall <- function(x,y){
  tau <- cor(x,y, method="kendall")
  # use="pairwise.complete.obs")
  # tau <- cor.fk(x)
  r <- sin(pi*tau/2)
  return(r)
}


cal.tetrachoric <- function(x,y){
  obj <- table(x,y)
  if(nrow(obj) != 2 | ncol(obj) != 2){
    res <- NA
  }else{
    if(obj[1,1] ==0 && obj[2,2]==0){
      res <- -1
    }else if(obj[1,2]==0 & obj[2,1]==0){
      res <- 1
    }else{
      idx <- obj == 0
      if(any(idx)){
        obj[idx] <- 0.5
      }
      obj <- obj/sum(obj)
      p11 <- obj[1,1]
      p12 <- obj[1,2]
      p21 <- obj[2,1]
      p22 <- obj[2,2]

      pr <- p11 + p12
      pc <- p11 + p21
      h <- qnorm(1-pr)
      k <- qnorm(1-pc)

      make.score.eq <- function(h,k,p11){
        function(r){
          mvtnorm::pmvnorm(lower=c(h,k),
          cor=matrix(c(1,r,r,1),2,2)) - p11
        }
      }

      score.eq <- make.score.eq(h,k,p11)

      res <- uniroot(score.eq,c(-1,1))$root
    }
  }
  return(res)
}

cal.r.BR <- function(x.bin, y.cont){
  x.bin <- as.numeric(factor(x.bin)) - 1
  if(length(unique(x.bin)) != 2){
    warning("Neither x nor y is binary variable. Use Kendall tau to calculate correlation")
    res <- cal.cor.kendall(x.bin, y,cont)
  }else{
    x.bar <- mean(x.bin)
    y.bar <- mean(y.cont)
    n <- length(x.bin)
    fac <- n * x.bar * y.bar
    nom <- sum(x.bin * y.cont) - fac
    den <- sum(sort(x.bin) * sort(y.cont)) - fac
    res <- nom/den
  }
  return(res)
}

cal.r.Lord <- function(x.bin, y.cont){
  res <- cal.r.BR(x.bin, y.cont)
  if(res < 0) res <- - cal.r.BR(x.bin, -y.cont)
  return(res)
}
  
biserial <- function(x,y){
  # x: binary 0 and 1;
  # y: continuous
  res <- cal.r.Lord(x.bin=x, y.cont=y)
  return(res)
}
  
polyserial <- function(x,y,ML=FALSE){
  # x: ordinal
  # y: continuous
  x.tab <- table(x)
  N <- sum(x.tab)
  x.cut <- qnorm(cumsum(x.tab)/N)
  x.cut <- x.cut[-length(x.cut)]
  res <- sqrt((N - 1)/N)*sd(x)*cor(x, y)/sum(dnorm(x.cut))
  if(res> 1) res <- 1
  if(res< -1) res <- -1
  return(res)
}

polychoric <- function(x,y,ML=FALSE){
  # x: oridinal
  # y: oridinal
  obj <- table(x,y)
  if(nrow(obj) < 2 | ncol(obj) < 2){
    res <- NA
  }else{
    if(FALSE){## Add 0.5 to 0 cell
      idx <- obj == 0
      if(any(idx)){
        obj[idx] <- 0.5
      }
    }
    R <- nrow(obj)
    C <- ncol(obj)
    tab <- obj/sum(obj)
    row.cut <- qnorm(cumsum(rowSums(obj))/sum(obj))[-R]
    col.cut <- qnorm(cumsum(colSums(obj))/sum(obj))[-C]

    make.like.fn <- function(obj,row.cut, col.cut){
      R <- nrow(obj)
      C <- ncol(obj)
      lo <- - Inf
      up <- Inf
      row.cut1 <- c(lo,row.cut,up)
      col.cut1 <- c(lo,col.cut,up)
      function(rho){
        COR <- matrix(c(1,rho,rho,1),2,2)
        PHI <- matrix(0,R,C)
        for (i in 1:R){
          a1 <- row.cut1[i]
          a2 <- row.cut1[i+1]
          for (j in 1:C){
            b1 <- col.cut1[j]
            b2 <- col.cut1[j+1]
            PHI[i,j] <- mvtnorm::pmvnorm(lower=c(a1,b1), upper=c(a2,b2), cor=COR)
          }
        }
        res <- - sum(obj* log(PHI))
        return(res)
      }
    }

    like.fn <- make.like.fn(obj,row.cut,col.cut)
    res <- optimize(like.fn,c(-1, 1))$minimum
  }
  return(res)
}


detect.type <- function(x){
  res <- "NULL"
  if(is.null(x)){
    res <- "NULL"
  }else{
    val <- unique(x)[!is.na(unique(x))]
    len <- length(val)
    if(len == 2){
      res <- "dichotomous"
    }else if (len > 2 & len <= 5){
      res <- "ordinal"
    }else{
      res <- "continuous"
    }
  }
  return(res)
}

mixed.cor <- function(x, y,kendall.only=FALSE,verbose=FALSE){
  idx <- complete.cases(x, y)
  x <- x[idx]
  y <- y[idx]

  num.x <- length(unique(x))
  num.y <- length(unique(y))
  if(kendall.only){
    x.type <- "continuous"
    y.type <- "continuous"
    x <- as.numeric(x)
    y <- as.numeric(y)
  }else{
    x.type <- detect.type(x)
    y.type <- detect.type(y)
  }
  if(x.type == "dichotomous" & y.type=="continuous"){ # biserial
    if(verbose) message("biserial:")
    x <- as.numeric(factor(x)) - 1
    res <- biserial(x,y)
  }else if(y.type == "dichotomous" & x.type == "continuous"){ # biserial
    if(verbose) message("biserial:")
    y <- as.numeric(factor(y)) - 1
    res <- biserial(y,x)
  }else if(x.type == "dichotomous" & y.type == "dichotomous"){ # tetrachoric
    if(verbose) message("tetrachoric:")
    x <- as.numeric(factor(x)) - 1
    y <- as.numeric(factor(y)) - 1
    res <- cal.tetrachoric(x,y)
  }else if(x.type == "ordinal" & y.type == "continuous"){ # polyserial
    if(verbose) message("polyserial:")
    x <- as.numeric(factor(x))
    res <- polyserial(x,y)
  }else if(y.type == "ordinal" & x.type == "continuous"){ # polyserial
    if(verbose) message("polyserial:")
    y <- as.numeric(factor(y))
    res <- polyserial(y,x)
  }else if((x.type == "ordinal" & y.type == "ordinal") |
    (x.type == "dichotomous" & y.type == "ordinal") |
    (x.type == "ordinal" & y.type == "dichotomous")){ # polychoric
    if(verbose) message("polychoric:")
    x <- as.numeric(factor(x))
    y <- as.numeric(factor(y))
    res <- polychoric(x,y)
  }else{ # if(x.type == "continuous" & y.type == "continuous"){ # kendall
    if(verbose) message("kendall correlation\n")
    res <- cal.cor.kendall(x,y)
  }
  return(res)
}

cal.cor.v <- function(obj,kendall.only=FALSE,verbose=FALSE){
  m <- ncol(obj)
  res <- rep(NA, m*(m-1)/2)
  k <- 1
  for (i in 1:(m-1)){
  for (j in (i+1):m){
    res[k] <- mixed.cor(obj[,i],obj[,j],kendall.only,
    verbose=verbose)
    k <- k + 1
  }}
  return(res)
}

cal.gamma.par <- function(Pheno,kendall.only=FALSE,verbose=FALSE){
  a1 <- 3.9081
  a2 <- 0.0313
  a3 <- 0.1022
  a4 <- -0.1378
  a5 <- 0.0941

  n <- nrow(Pheno)
  K <- ncol(Pheno)

  rho.v <- cal.cor.v(Pheno,kendall.only,verbose)
  rho.adj <- adj.cor(rho.v, n)

  vT <- 4*K + 2* sum(
    a1 * (rho.adj^2) +
    a2 * (rho.adj^4) +
    a3 * (rho.adj^6) +
    a4 * (rho.adj^8) +
    a5 * (rho.adj^10) -
    a1/n*(1-rho.adj^2)^2
  )
  ET <- 2*K
  v <- 2*ET^2/vT
  r <- vT/(2*ET)
  return(list(SHAPE=v/2,SCALE=2*r))
}
```
