## Supplementary Script for "Exploratory analysis of multiple traits co-adaptations in the population history": PolyGraphFunction20211001.R

```
# PolyGraph functions
# Originally implemented in Racimo, Berg and Pickrell (2018) Genetics 208:1565-1584
# Modified by R. Nakamichi


# make supergraph for PolyGraph from Treemix result
treemix2polygraph <- function(tm.tree){
  tm.vert <- tm.tree$d[,1:10]
  tm.edge <- tm.tree$e

  leaves <- as.character(tm.vert[!is.na(tm.vert[,2]),1])
  leaves.name <- tm.vert[!is.na(tm.vert[,2]),2]
  names(leaves.name) <- leaves
  inner_nodes <- tm.vert[is.na(tm.vert[,2]),1]
  inner_nodes.name <- paste0("x", inner_nodes)
  names(inner_nodes.name) <- inner_nodes

  root_node.name <- as.character(inner_nodes.name[as.character(tm.vert[tm.vert[,3]=="ROOT","V1"])])

  tm.edge12 <- tm.edge[,1:2]
  for(cli in names(leaves.name)){
    cln <- as.character(leaves.name[cli])
    tm.edge12[tm.edge12==cli] <- cln
  }
  for(cli in names(inner_nodes.name)){
    cln <- as.character(inner_nodes.name[cli])
    tm.edge12[tm.edge12==cli] <- cln
  }
  tm.edge[,1:2] <- tm.edge12
  tm.edge[tm.edge[,"V3"]<0.001, "V3"] <- 0.001
  leaves <- as.character(leaves.name)
  inner_nodes <- as.character(inner_nodes.name)

  edges <- NULL
  admixtures <- NULL
  admvalues <- NULL
  for(ce in 1:nrow(tm.edge)){
    edges <- c(edges, edge(as.character(tm.edge[ce,2]), as.character(tm.edge[ce,1])))
  }
  for(ce in 1:nrow(tm.edge)){
  if(tm.edge[ce,5]=="MIG"){
    cmig.o <- tm.edge[ce,"V2"]
    cmig <- tm.edge[tm.edge[,"V2"]==cmig.o,]
    coffs <- as.character(cmig.o)
    cpars <- as.character(cmig[,"V1"])
    edges <- c(edges, admixture_edge(coffs, cpars[1], cpars[2]))
    admixtures <- c(admixtures, admix_props(coffs, cpars[1], cpars[2], paste0("gamma",ce)))
    cadvalues <- as.numeric(cmig[,"V4"])
    admvalues <- rbind(admvalues, c(paste0("gamma",ce), cadvalues[1]))
  }}
  edges <- parent_edges(edges)

  if(sum(tm.edge$V5=="MIG")>0){  
    admixtures <- admixture_proportions(admixtures)
    if(nrow(admvalues) > 0){
      colnames(admvalues) <- c("ratename","rate")
      admvalues <- as.data.frame(admvalues)
      admvalues[,"rate"] <- as.numeric(as.character(admvalues[,"rate"]))
    }
  }

  edgevalues <- tm.edge[,c("V2","V1","V3")]
  colnames(edgevalues) <- c("child","parent","value")
  edgevalues[,"child"] <- as.character(edgevalues[,"child"])
  edgevalues[,"parent"] <- as.character(edgevalues[,"parent"])
  edgevalues[,"value"] <- as.numeric(as.character(edgevalues[,"value"]))
  edgevalues[edgevalues[,"value"]<0.001, "value"] <- 0.001

  graph <- agraph(leaves, inner_nodes, edges, admixtures)
  supergraph <- OrderBranches(list(graph, edgevalues, admvalues))

  return(list(
    supergraph=supergraph,
    leaves=leaves,
    edge=tm.edge,
    root_node.name=root_node.name
  ))
}

# Plot PCA of PolyGraph alpha on Treemix topology
plot.pca.alpha <- function(pcadata, pcares, axisA=1, axisB=2, title=NULL, PropVar=F, pop.label.only=F, ...){
  pca.score.label <- rownames(pcadata)
  i.alpha.pca <- grep("alpha_", pca.score.label)
  pca.score.label[i.alpha.pca] <- gsub("alpha_", "", pca.score.label[i.alpha.pca])
  if(pop.label.only){
    pca.score.label[i.alpha.pca] <- gsub("x[0-9]*", "", pca.score.label[i.alpha.pca])
    pca.score.label[i.alpha.pca] <- gsub("_", "", pca.score.label[i.alpha.pca])
  }
  pca.score.color <- rep("darkgreen", length(pca.score.label))
  pca.score.color[i.alpha.pca] <- "blue"
  is.inner.node <- pca.score.label==""

  pca.loading.label <- colnames(pcadata)
  pca.loading.color <- "red"

  pca.loading.scaled <- pcares$rotation * rep(pcares$sdev,each=ncol(pcadata)) *
                        sqrt(nrow(pcadata)/ncol(pcadata))
  pca.score.scaled <- pcares$x / rep(pcares$sdev, each=nrow(pcadata)) / sqrt(nrow(pcadata))

  propv <- summary(pcares)$importance[2, c(axisA, axisB)]
  propv <- sprintf("%.1f",propv*100)
  x.lab <- paste0("PC",axisA)
  if(PropVar){x.lab <- paste0(x.lab, " (", propv[1], "%)")}
  y.lab <- paste0("PC",axisB)
  if(PropVar){y.lab <- paste0(y.lab, " (", propv[2], "%)")}

  x.pca <- c(pca.loading.scaled[,axisA]*1.2, pca.score.scaled[,axisA])
  y.pca <- c(pca.loading.scaled[,axisB]*1.2, pca.score.scaled[,axisB])

  plot(x=x.pca, y=y.pca, type="n", xlab=x.lab, ylab=y.lab, main=title, ...)
  points(pca.score.scaled[is.inner.node,c(axisA,axisB)], col="gray60", pch=19)
  text(pca.score.scaled[,c(axisA,axisB)], pca.score.label, col=pca.score.color)
  text(pca.loading.scaled[,c(axisA,axisB)], pca.loading.label, col=pca.loading.color)
  arrows(x0=rep(0, ncol(pcadata)), y0=rep(0, ncol(pcadata)),
         x1=pca.loading.scaled[,axisA]*0.9, y1=pca.loading.scaled[,axisB]*0.9,
         col=pca.loading.color,length=0.1)
}

plot.pca.pop <- function(pcadata, pcares, col.loading=NULL, col.score=NULL, axisA=1, axisB=2, title=NULL, PropVar=F){
  pca.score.label <- rownames(pcadata)
  pca.score.color <- if(is.null(col.loading)){"blue"}else{col.loading}

  pca.loading.label <- colnames(pcadata)
  pca.loading.color <- if(is.null(col.score)){"red"}else{col.score}

  pca.loading.scaled <- pcares$rotation * rep(pcares$sdev,each=ncol(pcadata)) *
                        sqrt(nrow(pcadata)/ncol(pcadata))
  pca.score.scaled <- pcares$x / rep(pcares$sdev, each=nrow(pcadata)) / sqrt(nrow(pcadata))

  propv <- summary(pcares)$importance[2, c(axisA, axisB)]
  propv <- sprintf("%.1f",propv*100)
  x.lab <- paste0("PC",axisA)
  if(PropVar){x.lab <- paste0(x.lab, " (", propv[1], "%)")}
  y.lab <- paste0("PC",axisB)
  if(PropVar){y.lab <- paste0(y.lab, " (", propv[2], "%)")}

  plot(x=c(pca.loading.scaled[,axisA]*1.2, pca.score.scaled[,axisA]),
       y=c(pca.loading.scaled[,axisB]*1.2, pca.score.scaled[,axisB]),
       type="n", xlab=x.lab, ylab=y.lab, main=title)
  text(pca.score.scaled[,c(axisA,axisB)], pca.score.label, col=pca.score.color)
  text(pca.loading.scaled[,c(axisA,axisB)], pca.loading.label, col=pca.loading.color)
  arrows(x0=rep(0, ncol(pcadata)), y0=rep(0, ncol(pcadata)),
         x1=pca.loading.scaled[,axisA]*0.9, y1=pca.loading.scaled[,axisB]*0.9,
         col=pca.loading.color,length=0.1)
}


# Function for breakin graph into component pieces (from admixture graph package)
break_graph <- function(graph) {
  nodes <- graph$nodes
  edges <- list()
  admixtures <- list()
  parents <- graph$parents
  probs <- graph$probs
  for (i in seq(1, NROW(parents))) {
    match <- which(parents[i, ] == TRUE)
    if (length(match) == 0) {
      root <- nodes[i]
    } else if (length(match) == 1) {
      edges[[length(edges) + 1]] <- c(nodes[i], nodes[match[1]])
    } else if (length(match) == 2) {
      if (nchar(probs[i, match[1]]) > nchar(probs[i, match[2]])) {
        admixtures[[length(admixtures) + 1]] <- c(nodes[i], nodes[match[2]], nodes[match[1]], probs[i, match[2]])
      } else {
        admixtures[[length(admixtures) + 1]] <- c(nodes[i], nodes[match[1]], nodes[match[2]], probs[i, match[1]])
      }
    }
  }
  return(list(leaves = graph$leaves, inner_nodes = graph$inner_nodes, edges = edges, admixtures = admixtures, root = root))
}


# Function for choosing a tree given a particular combination of admixture choices
choosetree <- function(supergraph,chosenedges){
  
  if( length(chosenedges) > 2) {chosenedgestab <- t(matrix(chosenedges,ncol=2))
  } else{chosenedgestab <-  t(matrix(chosenedges))}
  
  test <- break_graph(supergraph[[1]])
  newedgevalues <- as.matrix(supergraph[[2]])
  newedgestab <- t(matrix(unlist(test$edges),nrow=2))
  newinner_nodes <- test$inner_nodes
  newleaves <- test$leaves
  
  # For each chosen path in list of chosen paths
  for(idx in seq(1,dim(chosenedgestab)[1])){
    
    firsthalf <- chosenedgestab[idx,]
    firsthalf.edge <- newedgevalues[which(newedgevalues[,1] == firsthalf[1] & newedgevalues[,2] == firsthalf[2]),3]
    secondhalf <- newedgestab[which(newedgestab[,2]==firsthalf[1]),]
    secondhalf.edge <- newedgevalues[which(newedgevalues[,1] == secondhalf[1] & newedgevalues[,2] == secondhalf[2]),3]
    taped <- c(secondhalf[1],firsthalf[2])
    taped.edge <- as.numeric(firsthalf.edge) + as.numeric(secondhalf.edge)
    
    newedgestab <- newedgestab[which(!(newedgestab[,1] == secondhalf[1] & newedgestab[,2] == secondhalf[2])),]
    newedgestab <- rbind(newedgestab,taped)
    rownames(newedgestab) <- c()
    
    nodetoextract <- secondhalf[2]
    newinner_nodes <- newinner_nodes[which(newinner_nodes != nodetoextract)]
    
    newedgevalues <- newedgevalues[which(!(newedgevalues[,1] == firsthalf[1] & newedgevalues[,2] == firsthalf[2])),]
    newedgevalues <- newedgevalues[which(!(newedgevalues[,1] == secondhalf[1] & newedgevalues[,2] == secondhalf[2])),]
    newedgevalues <- rbind(newedgevalues,c(taped,taped.edge))
    
    loneredge <- newedgevalues[which(newedgevalues[,1] == firsthalf[1] & newedgevalues[,2] != firsthalf[2]),]
    lonernode <- loneredge[2]
    
    firstloner <- newedgestab[which(newedgestab[,1] == lonernode),]
    firstloner.edge <- newedgevalues[which(newedgevalues[,1] == firstloner[1] & newedgevalues[,2] == firstloner[2]),3]
    secondloner <- newedgestab[which(newedgestab[,2] == lonernode),]
    secondloner.edge <- newedgevalues[which(newedgevalues[,1] == secondloner[1] & newedgevalues[,2] == secondloner[2]),3]
    tapedloner <- c(secondloner[1],firstloner[2])
    tapedloner.edge <- as.numeric(firstloner.edge) + as.numeric(secondloner.edge)
    
    newedgestab <- newedgestab[which(!(newedgestab[,1] == firstloner[1] & newedgestab[,2] == firstloner[2])),]
    newedgestab <- newedgestab[which(!(newedgestab[,1] == secondloner[1] & newedgestab[,2] == secondloner[2])),]
    newedgestab <- rbind(newedgestab,tapedloner)
    rownames(newedgestab) <- c()
    
    newinner_nodes <- newinner_nodes[which(newinner_nodes != lonernode)]
    
    newedgevalues <- newedgevalues[which(!(newedgevalues[,1] == firstloner[1] & newedgevalues[,2] == firstloner[2])),]
    newedgevalues <- newedgevalues[which(!(newedgevalues[,1] == secondloner[1] & newedgevalues[,2] == secondloner[2])),]
    newedgevalues <- rbind(newedgevalues,c(tapedloner,tapedloner.edge))
    
    # Remove dangling branch
    newedgevalues <- newedgevalues[which(!(newedgevalues[,1] == secondhalf[2] & newedgevalues[,2] == firstloner[1])),]
    
  }
  newedgevalues <- as.data.frame(newedgevalues)
  edgevec <- as.vector(matrix(t(newedgestab),nrow=1))
  edgevec <- t(matrix(edgevec,nrow=2))
  newedges <- cbind(edgevec,NA)

  newgraph <- admixturegraph::agraph(newleaves, newinner_nodes, newedges, NULL)
  
  supertree <- list(newgraph,newedgevalues)
  return(supertree)
  
}

# Function for extracting embedded trees and embedded probabilities of each tree from a graph
# Returns SUPERTREE object: 1) graph, 2) edge lengths, 3) product of chosen admixture rates, 4) adm path chosen, 5) intermediate paths for all edges
extract_the_trees <- function(supergraph){
  
  brokengraph <- break_graph(supergraph[[1]])
  graphedges <- supergraph[[2]]
  
  if(length(brokengraph$admixtures) > 0){
    
    alladm <- t(matrix((unlist(brokengraph$admixtures)),nrow=4))
    colnames(alladm) <- c("comb","left","right","ratename")
    alladm <- merge(supergraph[[3]],alladm)
    
    admA <- alladm[,c(2,3,4)]
    admB <- alladm[,c(2,3,5)]; admB[,1] <- 1 - admB[,1]
    admA <- apply(admA,1,function(x){paste(x,collapse="_")})
    admB <- apply(admB,1,function(x){paste(x,collapse="_")})
    admAB <- cbind(admA,admB)
    allchoices <- expand.grid(split(admAB, seq(nrow(admAB))))
    
    finaledges <- t(apply(allchoices,1,function(line){
      linetab <- t(matrix(unlist(sapply(line, function(x){
        strsplit(x,"_")
      })),nrow=3))
      finalprob <- prod(as.numeric(linetab[,1]))
      finaledges <-as.vector(t(linetab[,c(2,3)]))
      return(finaledges)
    }))
    
    finalprob <- as.vector(t(apply(allchoices,1,function(line){
      linetab <- t(matrix(unlist(sapply(line, function(x){
        strsplit(x,"_")
      })),nrow=3))
      finalprob <- prod(as.numeric(linetab[,1]))
      return(finalprob)
    })))
    
    
    treelist <- list()
    for(i in seq(1,dim(finaledges)[1])){
      treelist[[length(treelist)+1]] <- choosetree(supergraph,finaledges[i,])
      treelist[[length(treelist)]][[3]] <- finalprob[i]
      treelist[[length(treelist)]][[4]] <- finaledges[i,]
      
      intermediatelist <- apply(treelist[[length(treelist)]][[2]],1,function(x){
        list(GetConnection(x[1],x[2],finaledges[i,],graphedges))
        })
      namesinter <- apply(treelist[[length(treelist)]][[2]],1,function(x){
        paste(x[1],x[2],sep="_")
      })
      names(intermediatelist) <- namesinter
      treelist[[length(treelist)]][[5]] <- intermediatelist
    }
  }
  else{
    treelist <- list()
    treelist[[1]] <- list()
    treelist[[1]][[1]] <- supergraph[[1]]
    treelist[[1]][[2]] <- supergraph[[2]]
    treelist[[1]][[3]] <- 1
    treelist[[1]][[4]] <- NULL
    
    # Collect intermediate paths
    intermediatelist <- apply(treelist[[1]][[2]],1,function(x){
      list(GetConnection(x[1],x[2],finaledges[i,],graphedges))
    })
    namesinter <- apply(treelist[[1]][[2]],1,function(x){
      paste(x[1],x[2],sep="_")
    })
    names(intermediatelist) <- namesinter
    treelist[[1]][[5]] <- intermediatelist
    
  }
  return(treelist)
}


ZeroOne <- function(vecnum){
  return(pmax(0.001,pmin(0.999,vecnum)))
}


# Function for obtaining the log-density of a path in a tree (selection based on size of effect)
#log_prob_path_sel <- function(child,parent,drift,alpha,effect){
#  if(parent <= 0){parent <- 0.001
#  } else if(parent >= 1){parent <- 0.999}
#  variance <- drift*parent*(1-parent)
#  sel <- parent*(1-parent)*alpha*effect
#  density <- log(dtnorm(child,parent+sel,sqrt(variance),0,1))
#  return(max(-100000,density))
#}

# Function for obtaining the log-density of a path in a tree (selection based on sign of effect)
#log_prob_path_selsign <- function(child,parent,drift,alpha,effect){
#  if(parent <= 0){parent <- 0.001
#  } else if(parent >= 1){parent <- 0.999}
#  variance <- drift*parent*(1-parent)
#  sel <- parent*(1-parent)*alpha*sign(effect)
#  density <- log(dtnorm(child,parent+sel,sqrt(variance),0,1))
#  return(max(-100000,density))
#}

# Function for obtaining the density of a path in a tree (selection based on sign of effect)
#prob_path_selsign <- function(child,parent,drift,alpha,effect){
#  parent <- pmax(0.001,pmin(0.999,parent))
#  variance <- drift*parent*(1-parent)
#  sel <- parent*(1-parent)*alpha*sign(effect)
#  density <- dtnorm(child,parent+sel,sqrt(variance),0,1)
#  return(max(0,density))
#}


# Function for obtaining the density of a path in a tree (selection based on sign of effect)
prob_path_selsign <- function(child,parent,drift,alpha,effect){
  parent <- ZeroOne(parent)
  variance <- drift*parent*(1-parent)
  sel <- parent*(1-parent)*alpha*sign(effect)
  density <- dnorm(child,parent+sel,sqrt(variance))
  return(max(0,density))
}


#log_prob_path <- function(child,parent,drift){
#  if(parent <= 0){parent <- 0.001
#  } else if(parent >= 1){parent <- 0.999}
#  variance <- drift*parent*(1-parent)
#  density <- log(dtnorm(child,parent,sqrt(variance),0,1))
#  return(max(-100000,density))
#}


# Function for obtaining the total log-density of a tree
#log_prob_tree <- function(supertree,freqs,graphedges,alphas,effects,runmode){
#  
#  edgevalues <- supertree[[2]]
#  finaledges <- supertree[[4]]
#  intermediatelist <- supertree[[5]]
#  
#  freqs <- cbind(freqs,effects)
#  finalvec <- apply(freqs,1,function(freqline){
#    probsnp <- apply(as.matrix(edgevalues),1,function(x){
#      parent <- x[2]
#      child <- x[1]
#      effect <- freqline[length(freqline)]
#     
#      pair <- paste(child,parent,sep="_")
#      intermediates <- intermediatelist[[which(names(intermediatelist) == pair)]][[1]]
#      probtosum <- apply(intermediates,1,function(y){
#        Xchild <- y[1]
#        Xparent <- y[2]
#        Xdrift <- as.numeric(y[3])
#        Xalpha <- as.numeric(alphas[which(alphas[,1] == Xchild & alphas[,2] == Xparent),3])
#        Xchild_freq <- freqline[which(names(freqline) == Xchild)]
#        Xparent_freq <- freqline[which(names(freqline) == Xparent)]
#        Xprob <- log_prob_path_selsign(Xchild_freq,Xparent_freq,Xdrift,Xalpha,effect)
#        return(Xprob)
#      })
#      prob <- sum(probtosum)
#      return(prob)
#    })
#    sumprobsnp <- sum(probsnp)
#    return(sumprobsnp)
#  })
#  return(finalvec)
#}


# Function for obtaining the total log-density of an admixture graph
#pgraph_old <- function(treelist,graphedges,freqs,alphas,effects,runmode){
#
#  # Obtain probability of each tree in a vector - Using Rmpfr precision library
#  all_prob_trees <- rep(0,dim(freqs)[1])
#  for( i in seq(1,length(treelist))){
#    admixprob <- treelist[[i]][[3]]
#    
#    logtreeprob <- log_prob_tree(treelist[[i]],freqs,graphedges,alphas,effects,runmode)
#    logtreeprob <- mpfr(logtreeprob, precBits = 256)
#    prob_tree <- exp(logtreeprob)
#    all_prob_trees <- all_prob_trees + admixprob*prob_tree
#  }
#  return(as.numeric(log(all_prob_trees)))
#}


DeconstructGraph <- function(supergraph){
  
  graphedges <- supergraph[[2]]
  nameadmixpars <- as.character(supergraph[[3]][,1])

  allbranches <- sapply( seq(1,dim(supergraph[[1]]$parents)[1]), function(x){
    child <- rownames(supergraph[[1]]$parents)[x]
    idxchild <- x
    parents <- which(supergraph[[1]]$parents[x,] == TRUE)
    if( length(parents) == 0 ){ return(c(NaN,NaN,NaN,NaN,NaN,NaN,NaN,FALSE))
    } else if( length(parents) == 1){
      parent <- names(parents)
      branch <- graphedges[which(graphedges[,1] == child & graphedges[,2] == parent),3]
      return(c(child,parent,branch,1,NaN,NaN,NaN,FALSE))
    } else{
      parentsrates <- supergraph[[1]]$probs[x,][ which(names(supergraph[[1]]$probs[x, ]) %in% names(parents) ) ]
      recrate <- parentsrates[which(parentsrates %in% nameadmixpars)]
      unrecrate <- parentsrates[which(!(parentsrates %in% nameadmixpars))]
      parentA <- names(recrate)
      parentB <- names(unrecrate)
      rateA <- supergraph[[3]][which(nameadmixpars == recrate),2]
      rateB <- 1 - rateA
      branchA <- graphedges[which(graphedges[,1] == child & graphedges[,2] == parentA),3]
      branchB <- graphedges[which(graphedges[,1] == child & graphedges[,2] == parentB),3]
      return(c(child,parentA,branchA,rateA,parentB,branchB,rateB,TRUE))
    }
  })

  allbranches <- t(matrix(unlist(allbranches),nrow=8))
  allbranches <- allbranches[which(allbranches[,1] != "NaN"),]
  colnames(allbranches) <- c("child","parent","branch","admixrate","parentB","branchB","admixrateB","admevent")
  
  return(allbranches)
  
}

# Log-likelihood of graph
pgraph <- function(deconsgraph,freqs,effects,alphas){
  
  freqs <- cbind(freqs,effects)
  finalvec <- apply(freqs,1,function(freqline){
    probsnp <- apply(deconsgraph,1,function(x){
      
      effect <- freqline[length(freqline)]
      
      child <- x[1]
      childfreq <- freqline[child]
      
      if(x[8] == "FALSE"){
        
        parent <- x[2]
        branch <- as.numeric(x[3])
        admixrate <- as.numeric(x[4])
        alpha <- as.numeric(alphas[which(alphas[,1] == child & alphas[,2] == parent),3])
        childfreq <- freqline[x[1]]
        parentfreq <- freqline[parent]
        return(log(prob_path_selsign(childfreq,parentfreq,branch,alpha,effect)))
        
      } else{

        parentA <- x[2]
        parentAfreq <- freqline[parentA] 
        branchA <- as.numeric(x[3])
        admixrateA <- as.numeric(x[4])
        alphaA <- as.numeric(alphas[which(alphas[,1] == child & alphas[,2] == parentA),3])
        
        parentB <- x[5]
        parentBfreq <- freqline[parentB]
        branchB <- as.numeric(x[6])
        admixrateB <- as.numeric(x[7])
        alphaB <- as.numeric(alphas[which(alphas[,1] == child & alphas[,2] == parentB),3])
        return( log( admixrateA*prob_path_selsign(childfreq,parentAfreq,branchA,alphaA,effect) + 
                       admixrateB*prob_path_selsign(childfreq,parentBfreq,branchB,alphaB,effect) ) )

      }
    })
  sumprobsnp <- sum(probsnp)
  return(sumprobsnp)
  })
    
return(finalvec)
}


# Binomial sampling log likelihood
pbinom <- function(input_leaf_der,input_leaf_tot,innerfreqs){
  
  leaf_innerfreqs <- innerfreqs[,colnames(input_leaf_der)]
  totalpops <- dim(input_leaf_der)[2]
  logbinomprobs <- sapply(seq(1,totalpops),function(x){
    sampprob <- ZeroOne(leaf_innerfreqs[,x])
    logprob <- dbinom(input_leaf_der[,x],input_leaf_tot[,x], sampprob,log=TRUE)
    return(logprob)
    })
  finallogprob <- apply(logbinomprobs,1,sum)
  return(finallogprob)
  
}


# Function for proposing frequencies (truncated normal)
#propose_innerfreqs <- function(freqs,proposize,numSNPs){
#  freqs <- as.numeric(freqs)
#  newinnerfreqs_vec <-  rtnorm(length(freqs),freqs,proposize,lower=0,upper=1)
#  newinnerfreqs_mat <- matrix(newinnerfreqs_vec,nrow=numSNPs)
#  return(newinnerfreqs_mat)
#}

# Another function for proposing frequencies (truncated normal)
#propose_innerfreqsB <- function(freqs,proposize,numSNPs){
#  freqs <- as.numeric(freqs)
#  newinnerfreqs_vec <-  rtnorm(length(freqs),freqs,proposize,lower=0,upper=1)
#  newinnerfreqs_mat <- matrix(newinnerfreqs_vec,nrow=numSNPs)
#  return(newinnerfreqs_mat)
#}

# Another function for proposing frequencies (Normal)
propose_innerfreqsB <- function(freqs,proposize,numSNPs){
  freqs <- as.numeric(freqs)
  freqs <- ZeroOne(freqs)
  newinnerfreqs_vec <-  rnorm(length(freqs),freqs,proposize*freqs*(1-freqs))
  newinnerfreqs_mat <- matrix(newinnerfreqs_vec,nrow=numSNPs)
  return(newinnerfreqs_mat)
}


# Function for proposing frequencies
#update_innerfreqs <- function(freqs,proposize,numSNPs){
#  newinnerfreqs_vec <- sapply(freqs,function(x){return(x+sample(c(-1,1),1)/10000)})
#  newinnerfreqs_mat <- matrix(newinnerfreqs_vec,nrow=numSNPs)
#  return(newinnerfreqs_mat)
#}

# Function for proposing alphas
#propose_alphas <- function(alphas,proposize){
#  numalphas <- dim(alphas)[1]
#  newalphas_vec <- as.numeric(alphas[,3]) + rnorm(numalphas,0,proposize)
#  newalphas <- cbind(alphas[,1],alphas[,2],as.character(newalphas_vec))
#  colnames(newalphas) <- colnames(alphas)
#  return(newalphas)
#}

propose_alphasB <- function(alphas,proposize,teststat=c(),qbfactor){
  numalphas <- dim(alphas)[1]
  if(length(teststat) > 0){
    if(qbfactor != 0){cutoff <- max(teststat) / qbfactor
    } else{ cutoff <- qchisq( 1 - (0.05/length(teststat)), df=1)}
    teststat[which(teststat < cutoff)] <- 0
    weightsalphas = as.vector(teststat/sum(teststat))
    tochangeidx <- sample(seq(1,numalphas),1,prob = weightsalphas)
  } else { tochangeidx <- sample(seq(1,numalphas),1) }
  tochangealpha <- alphas[tochangeidx,3]
  newalpha <- as.character(as.numeric(tochangealpha) + rnorm(1,0,proposize))
  newalphas <- alphas
  newalphas[tochangeidx,3] <- newalpha
  colnames(newalphas) <- colnames(alphas)
  return(newalphas)
}

#propose_alphas_dir <- function(alphas,proposize){
#  numalphas <- dim(alphas)[1]
#  
#  oldvec <- as.numeric(alphas[,3])
#
#  shuffledvec <- sample(standardvec)
#  newmass <- rnorm(1,totalmass,proposize)
#  newalphas_vec <- shuffledvec*newmass
#
#  newalphas <- cbind(alphas[,1],alphas[,2],as.character(newalphas_vec))
#  colnames(newalphas) <- colnames(alphas)
#  return(newalphas)
#
#}


# Inner freqs prior (Uniform) - returns a vector with one element per SNP
log_innerfreqs_prior <- function(innerfreqs,lower,upper){
  innerfreqs <- matrix(ZeroOne(innerfreqs),nrow=dim(innerfreqs)[1])
  densities <- dunif(innerfreqs,lower,upper,log=TRUE)
  #densities[which(densities < -1)] <- -10000
  finaldens <- apply(densities,1,function(x){sum(x)})
  return(finaldens)
}

# Alphas prior (Normal)
log_alphas_prior <- function(alphas,mean,stdev){
  densities <- dnorm(alphas,mean,stdev,log=TRUE)
  return(sum(densities))
}

# Spike and slab function
spikeslab <- function(x,mean,sdslab,sdspike,prop){
  spike <- dnorm(x,mean,sdspike)
  slab <- dnorm(x,mean,sdslab)
  return(prop*spike+(1-prop)*slab)
}


# Spike-and-slab prior
log_alphas_prior_spikeslab <- function(alphas,mean,sd1,ssfactor,prop){
  densities <- log(spikeslab(alphas,mean,sd1,sd1/ssfactor,prop))
  return(sum(densities))
}


# Transition ratio for frequencies
translogdiff_innerfreqs <- function(old_innerfreqs,new_innerfreqs,innerfreqs_proposize,numSNPs){
  old_innerfreqs <- as.numeric(old_innerfreqs)
  new_innerfreqs <- as.numeric(new_innerfreqs)
  old_innerfreqs <- ZeroOne(old_innerfreqs)
  new_innerfreqs <- ZeroOne(new_innerfreqs)
  numerator <- dnorm(old_innerfreqs,new_innerfreqs,innerfreqs_proposize*(new_innerfreqs)*(1-new_innerfreqs),log=TRUE)
  denominator <- dnorm(new_innerfreqs,old_innerfreqs,innerfreqs_proposize*(old_innerfreqs)*(1-old_innerfreqs),log=TRUE)
  diff <- numerator - denominator
  diff_mat <- matrix(diff,nrow=numSNPs)
  return(diff_mat)
}

RemoveFixed <- function(nodes){
    nodes[which(nodes==1)] <- 0.999
    nodes[which(nodes==0)] <- 0.001
    return(nodes)
}


# Get child-ancestor connection possibly involving more than one intermediate parent 
GetConnection <- function(child,parent,finaledges,graphedges){
  
  retrieve <- which(graphedges[,1] == child & graphedges[,2] == parent)
  if(length(retrieve) == 1){
    branches <- cbind(child,parent,as.character(graphedges[retrieve,3]))
  } else{
      tempparent <- NaN
      tempchild <- child
      allnodes <- c(child)
      while(parent != tempparent){
        if( tempchild %in% finaledges[which(seq(1,length(finaledges)) %% 2 == 1)] ){
          tempparent <- finaledges[which(finaledges == tempchild)+1]
          allnodes <- c(allnodes,tempparent)
          tempchild <- tempparent
        } else{
          tempparent <- as.character(graphedges[which(graphedges[,1] == tempchild),2])
          allnodes <- c(allnodes,tempparent)
          tempchild <- tempparent
        }
      }
      branches <- c()
      for( i in seq(1,length(allnodes)-1)){
        retrieve <- which(graphedges[,1] == allnodes[i] & graphedges[,2] == allnodes[i+1])
        branches <- rbind(branches, c(allnodes[i],allnodes[i+1],as.character(graphedges[retrieve,3])))
      }
  }
  return(matrix(branches,ncol=3))
}


min.f1f2 <- function(x, mu1, mu2, sd1, sd2) {
  f1 <- dnorm(x, mean=mu1, sd=sd1)
  f2 <- dnorm(x, mean=mu2, sd=sd2)
  return(pmin(f1, f2))
}

integ.min.f1f2 <- function(m1,m2,v1,v2){
  s1 <- sqrt(v1)
  s2 <- sqrt(v2)
  return(integrate(min.f1f2, -Inf, Inf, mu1=m1, mu2=m2, sd1=s1, sd2=s2)$value)
}

shift <- function(x, n = 1) {
  if (n == 0) x else c(tail(x, -n), head(x, n))
}


AncestralAverage <- function(leaf_vector,supergraph){
  
  recorded <- leaf_vector
  notrecorded <- supergraph[[1]]$inner_nodes
  
  while( length(notrecorded) != 0 ){
    
    testinner <- notrecorded[1]
    rowidx <- which(rownames(supergraph[[1]]$children) == testinner)
    colidx <- which(supergraph[[1]]$children[rowidx,] == TRUE)
    children <- names(colidx)
    
    if( 0 %in% ( children %in% names(recorded)) ){
      notrecorded <- shift(notrecorded)
    } else{
      assigned <- mean(recorded[children])
      recorded <- c(recorded,assigned)
      names(recorded)[length(names(recorded))] <- testinner
      notrecorded <- notrecorded[-1]
    }
    
  }
  
  innerfinal <- recorded[supergraph[[1]]$inner_nodes]
  return(innerfinal)
  
}


CollectBranches <- function(supergraph,targetbranch,deconsgraph,leaves){
  
  graphedges <- supergraph[[2]]
  
  # Start from child
  firstparent <- targetbranch[1]
  
  # Depth-first search of all downstream paths
  tovisit <- c(firstparent)
  branchpaths <- c()
  discovered <- c()
  while(length(tovisit) != 0){
    check = tovisit[length(tovisit)]
    tovisit = tovisit[-length(tovisit)]
    
    
    if (!(check %in% discovered)){
      
      discovered <- c(discovered,check)
      daughters <-  as.character(graphedges[graphedges$parent == check,1])
      for(daughter in daughters){
        
        daughtervec <- deconsgraph[which(deconsgraph[,1] == daughter),]
        
        
        if(daughtervec[2] == check){ prob <- as.numeric(daughtervec[4])
        } else if(daughtervec[5] == check){ prob <- as.numeric(daughtervec[7])}
        
        path <- c(daughter,check,prob)
        branchpaths <- rbind(branchpaths,path)
        
      }
      if(length(daughters) > 0){
        tovisit <- c(tovisit,daughters)
      }
    }
  }
  

  
  prevpaths <- matrix(NaN)
  # Connect all possible paths from leaves to the starting parent
  #for(rep in seq(1,10)){
  while(TRUE){  
    for(pathidx in which(branchpaths[,1] %in% leaves)){
      path <-  branchpaths[pathidx,]
      for(conidx in which(branchpaths[,1] == path[2])){
        con <- branchpaths[conidx,]
        branchpaths <- rbind(branchpaths, c( path[1], con[2], as.numeric(path[3]) * as.numeric(con[3]) )  )
      }
    }
    
    branchpaths <- unique(branchpaths)
    currpaths <- branchpaths
    #print(branchpaths)
    #print(dim(currpaths))
    #print(dim(prevpaths))
    if( prod(dim(currpaths) == dim(prevpaths))){break}
    prevpaths <- currpaths
    
  }
  

  # Only keep paths that go from a leaf to the starting parent
  branchpaths <- unique(branchpaths[which(branchpaths[,1] %in% leaves & branchpaths[,2] == firstparent),])
  
  leafcontrib <- c()
  
  # Get first adm probability
  candidate <- which(deconsgraph[,1] == targetbranch[1] & deconsgraph[,2] == targetbranch[2])
  if(length(candidate) == 1){ 
    firstprod <- as.numeric(deconsgraph[candidate,4])
  } else{
    candidate <- which(deconsgraph[,1] == targetbranch[1] & deconsgraph[,5] == targetbranch[2])
    firstprod <- as.numeric(deconsgraph[candidate,7])
  }
  if(is.vector(branchpaths)){ branchpaths <- t(matrix(branchpaths))}
  
  
  # Compute vectors of leaf-contributions
  if(firstparent %in% leaves){
    leafcontrib <- as.numeric(leaves == firstparent)
  } else {
    for(leaf in leaves){
      if( !(leaf %in% branchpaths[,1]) ){
        leafcontrib <- c(leafcontrib,0)
      } else{
        
        tempcontrib <- c()
        i <- 1
        for( currparent in branchpaths[which(branchpaths[,1] == leaf),2]){
          currchild <- leaf
          prod <- firstprod * as.numeric(branchpaths[which(branchpaths[,1] == currchild & branchpaths[,2] == currparent),3][i])
          tempcontrib <- c(tempcontrib,prod)
          i <- i + 1
        }
        
        leafcontrib <- c(leafcontrib,sum(tempcontrib))
      }
    }
  }
  
  return(leafcontrib)
  
}


OrderBranches <- function(supergraph){

  oldedgevalues <- supergraph[[2]]
  root <- as.character(unique(supergraph[[2]]$parent[!(supergraph[[2]]$parent %in% supergraph[[2]]$child)]))
  newedgevalues <- c()
  parents <- c(root)
  
  while( length(parents) < length(supergraph[[1]]$nodes) ){

    children_to_test <- supergraph[[1]]$nodes[which( !(supergraph[[1]]$nodes %in% parents) )]
    
    for( child in children_to_test ){
      child <- as.character(child)
      
      allparents <- names(which(supergraph[[1]]$parents[child,]))
      
      if ( prod(names(which(supergraph[[1]]$parents[child,])) %in% parents) == 1){
        toadd <- oldedgevalues[oldedgevalues$child == child,]
        newedgevalues <- rbind(newedgevalues,toadd)
        parents <- c(parents,child)
      }
      parents <- unique(parents)
    }
  }
  
  newedgevalues <- data.frame(newedgevalues)
  colnames(newedgevalues) <- c("child","parent","value")
  
  #print(newedgevalues)
  
  supergraph <- list(supergraph[[1]],newedgevalues,supergraph[[3]])
  
  return(supergraph)
}


norm_vec <- function(x) sqrt(sum(x^2))


ChiSquared <- function(supergraph,leaves_freqs,effects,neut_leaves_freqs,total=FALSE,randomize=FALSE){
  
  graphedges <- supergraph[[2]]
  deconsgraph <- DeconstructGraph(supergraph)
  leaves <- supergraph[[1]]$leaves
  
  checkseg <- which( apply(leaves_freqs,1,sum)/length(leaves) < 0.99  & apply(leaves_freqs,1,sum)/length(leaves) > 0.01 )
  leaves_freqs <- leaves_freqs[checkseg,]
  effects <- effects[checkseg]
  
  # Randomize effects if necessary
  if(randomize == TRUE){effects <- effects * sample(c(-1,1),length(effects),replace=TRUE)}
  
  # Compute empirical covariance matrix
  checksegneut <- which( apply(neut_leaves_freqs,1,sum)/length(leaves) < 0.99  & apply(neut_leaves_freqs,1,sum)/length(leaves) > 0.01 )
  neut_leaves_freqs <- neut_leaves_freqs[checksegneut,]
  
  neut_leaves_freqs_means <- apply(neut_leaves_freqs, 1, mean)
  mean_hetero <- neut_leaves_freqs_means*(1-neut_leaves_freqs_means)
  numSNPs <- length(neut_leaves_freqs_means)
  
  #Fmat <- sapply(seq(1,dim(neut_leaves_freqs)[2]),function(x){
  #  sapply(seq(1,dim(neut_leaves_freqs)[2]),function(y){
  #    cov(neut_leaves_freqs[,x],neut_leaves_freqs[,y])
  #  })
  #})
  
  Fmat <- sapply(seq(1,dim(neut_leaves_freqs)[2]),function(x){
    sapply(seq(1,dim(neut_leaves_freqs)[2]),function(y){
      cov(neut_leaves_freqs[,x]/sqrt(mean_hetero), neut_leaves_freqs[,y]/sqrt(mean_hetero))
    })
  })
  
  colnames(Fmat) <- colnames(neut_leaves_freqs)
  rownames(Fmat) <- colnames(neut_leaves_freqs)
 
  
  # Compute contributions of each branch to each leaf
  if(total == FALSE){
  contribmat <- c()
  contribmat_names <- c()
  for(branchidx in seq(1,dim(graphedges)[1])){
    targetbranch <- c(as.character(graphedges[branchidx,1]),as.character(graphedges[branchidx,2]))
    name <- paste(targetbranch,collapse="_")
    branchvec <- CollectBranches(supergraph,targetbranch,deconsgraph,leaves)
    contribmat <- rbind(contribmat,branchvec)
    contribmat_names <- rbind(contribmat_names,name)
  }
  rownames(contribmat) <- contribmat_names
  colnames(contribmat) <- leaves
  
  # Standardize vectors to be of unit length
  contribmat <- t(apply(contribmat,1, function(x) {
    x <- x - mean(x)
    return(x / norm_vec(x))
  }))
  }
  
  # Compute mean genetic values
  meangen <- apply(leaves_freqs * effects, 2, function(x){sum(x)})

  # Scale by average genetic value
  meangen <- (meangen - mean(meangen))

  # Compute the estimated ancestral genetic variance over all populations
  meanfreqs <- apply(leaves_freqs,1,mean)
  
  varmean <- sum(sapply(seq(1,length(meanfreqs)),function(i){
  score = meanfreqs[i]*(1-meanfreqs[i])*effects[i]^2
  return(score)
  }))
  
  if(total == FALSE){
  contribmat <- contribmat[,names(meangen)]
  }
  
  if(total == FALSE){
  # Compute Q_B test statistic
  Qteststat <- apply(contribmat,1, function(x){
    #print(varmean)
    lambda <- varmean * (t(x) %*% Fmat %*% x)
    numerator <- (meangen %*% x)^2
    final <- numerator / (lambda)
    return(final)
  } )
  
  # Compute q_B test statistic
  qteststat <- apply(contribmat,1, function(x){
    #print(varmean)
    lambda <- varmean * (t(x) %*% Fmat %*% x)
    numerator <- (meangen %*% x)
    final <- numerator / sqrt(lambda)
    return(final)
  } )
  
  branchorder <- apply(graphedges,1,function(x){paste(as.character(x[1]),as.character(x[2]),sep="_")})
  Qteststat <- Qteststat[branchorder]
  Pval <- 1 - pchisq(Qteststat,1)
  qteststat <- qteststat[branchorder]
  allstats <- cbind(Qteststat,qteststat,Pval)
  
  }
  
  if(total == TRUE){
    
    # Compute Q_X statistic
    numerator <- t(meangen) %*% solve(Fmat) %*% meangen
    denominator <- varmean
    Qteststat <- numerator / denominator
    
    Pval <- 1 - pchisq(Qteststat,qr(Fmat)$rank)
    allstats <- c(Qteststat,NaN,Pval)
  }

  return(allstats)
}


ObtainFreqs <- function(countdat){
  freqs <- apply(countdat,c(1,2),function(x){splitted <- strsplit(x,",")[[1]]; return( as.numeric(splitted[2]) / (as.numeric(splitted[2])+as.numeric(splitted[1])) )})
  return(as.matrix(freqs))
}

# Trim out white space
trim <- function (x) gsub("^\\s+|\\s+$", "", x)

LoadCounts <- function(filename,pops){
  table <- as.matrix(read.table(filename,header=TRUE,sep="\t",strip.white=TRUE))
  tokeep <- which(colnames(table) %in% pops)
  tokeep <- c(1,2,3,4,tokeep)
  table <- table[,tokeep, drop=F]
  table[,1] <- trim(table[,1])
  table[,2] <- trim(table[,2])
  return(table)
}


# Correct positions
CorrectPos <- function(inputvals,coords,listvertices,root,minbranch=0.075){
  # Use only one parent branch when dealing with admixed pops
  inputvals <- inputvals[!duplicated(inputvals[,1]),]
  
  todo <- as.matrix(inputvals[inputvals[,2] == root,])
  done <- rbind( c(root,coords[listvertices == root,]) )
  
  while(length(todo) > 0){
    
    if( !(is.vector(todo)) ){
      elem <- todo[1,1]
      parent <- todo[1,2]
    } else {
      elem <- todo[1]
      parent <- todo[2]
    }
    
    elemx <- coords[which(listvertices == elem),1]
    elemy <- coords[which(listvertices == elem),2]
    parenty <- as.numeric(done[which(done[,1] == parent),3])
    minbranch <- as.numeric(minbranch)
    addy <- max(minbranch,inputvals[which(as.character(inputvals[,1]) == elem & as.character(inputvals[,2]) == parent),3])
    addy <- as.numeric(addy)
    newy <- parenty - addy
    
    done <- rbind(done, c(elem,elemx,newy))
    
    if( !(is.vector(todo)) ){
      todo <- todo[-1,]
    } else {
      todo <- c()
    }
    
    if(sum(as.character(inputvals[,2]) == elem) > 0){
      toadd <- inputvals[as.character(inputvals[,2]) == elem,]
      toadd <- sapply(toadd,as.character)
      
      if(is.vector(toadd)){
        todo <- rbind(todo,toadd)
      } else{ 
        for(j in seq(1,dim(toadd)[1])){
          todo <- rbind(todo,toadd[j,])
        }
      }
    }
  }
  
  done <- done[!duplicated(done[,1]),]
  done <- done[match(listvertices,done[,1]),]
  
  final <-  t(apply(done[,c(2,3)],1,as.numeric))
  return(final)
}


# Image-scale function created by Pretty R at inside-R.org
image.scale <- function(z, zlim, col = heat.colors(12),
                        breaks, horiz=TRUE, ylim=NULL, xlim=NULL, ...){
  if(!missing(breaks)){
    if(length(breaks) != (length(col)+1)){stop("must have one more break than colour")}
  }
  if(missing(breaks) & !missing(zlim)){
    breaks <- seq(zlim[1], zlim[2], length.out=(length(col)+1)) 
  }
  if(missing(breaks) & missing(zlim)){
    zlim <- range(z, na.rm=TRUE)
    zlim[2] <- zlim[2]+c(zlim[2]-zlim[1])*(1E-3)#adds a bit to the range in both directions
    zlim[1] <- zlim[1]-c(zlim[2]-zlim[1])*(1E-3)
    breaks <- seq(zlim[1], zlim[2], length.out=(length(col)+1))
  }
  poly <- vector(mode="list", length(col))
  for(i in seq(poly)){
    poly[[i]] <- c(breaks[i], breaks[i+1], breaks[i+1], breaks[i])
  }
  xaxt <- ifelse(horiz, "s", "n")
  yaxt <- ifelse(horiz, "n", "s")
  if(horiz){YLIM<-c(0,1); XLIM<-range(breaks)}
  if(!horiz){YLIM<-range(breaks); XLIM<-c(0,1)}
  if(missing(xlim)) xlim=XLIM
  if(missing(ylim)) ylim=YLIM
  plot(1,1,t="n",ylim=ylim, xlim=xlim, xaxt=xaxt, yaxt=yaxt, xaxs="i", yaxs="i", ...)  
  for(i in seq(poly)){
    if(horiz){
      polygon(poly[[i]], c(0,0,1,1), col=col[i], border=NA)
    }
    if(!horiz){
      polygon(c(0,0,1,1), poly[[i]], col=col[i], border=NA)
    }
  }
}


# Make a graph network plot
MakeGraphPlotInterior <- function(inputvals, selcoefs, root, inner_nodes=NULL, title,
                                  minsel=NULL, maxsel=NULL, xlabname=NULL, minbranch=0.075){
  
  if(is.null(minsel)){minsel <- floor(min(as.numeric(selcoefs))*100)/100}
  if(is.null(maxsel)){maxsel <- ceiling(max(as.numeric(selcoefs))*100)/100}
  maxsel <- max(abs(minsel), abs(maxsel))
  minsel <- -maxsel  


  g <- graph_from_data_frame(cbind(as.character(inputvals[,2]),as.character(inputvals[,1])))
  
  colfunc <- colorRampPalette(c("blue","grey","red"),space="rgb")
  colrange <- seq(minsel*100,maxsel*100,1)
  colpal <- colfunc(length(colrange))
  colpal <- cbind(colrange,colpal)
  
  breaks <- seq(minsel, maxsel,length.out=200)
  
  E(g)$drifts <- as.character(inputvals[,3])
  E(g)$selcoefs <- selcoefs
  
  E(g)$color <- sapply(E(g)$selcoefs, function(x){
    ccol <- colpal[which(colpal[,1] == round(as.numeric(x)*100)),2]
    if(length(ccol>0)){ccol}else{if(x<0){colpal[1,2]}else{colpal[nrow(colpal),2]}}
  })
  #E(g)$width <- round(2*log(as.numeric(E(g)$drifts)*500))
  E(g)$width <- rep(5,length(E(g)$drifts))
  
  coords <- layout.reingold.tilford(g,root=root)
  
  listvertices <- lapply(list.vertex.attributes(g),function(x) get.vertex.attribute(g,x))[[1]]
  
  coords <- CorrectPos(inputvals,coords,listvertices,root,minbranch)
  
  coords[,2] <- jitter(coords[,2],1)
  
  layout(matrix(c(1,2), nrow=2, ncol=1), widths=c(1), heights=c(6,1))
  #layout.show(2)

  cvg <- names(V(g))
  inner.cvg <- is.element(cvg, inner_nodes)
  vertex.label.color <- ifelse(inner.cvg, "black", "black")
  vertex.label.size <- ifelse(inner.cvg, 0.7, 1.2)
  vertex.frame.color <- ifelse(inner.cvg, NA, NA)
  vertex.color <- ifelse(inner.cvg, "white", "yellow")
  vertex.size <- ifelse(inner.cvg, 5, 9)

  par(mar=c(1,1,1,1))
  plot(g, layout=coords, edge.color=E(g)$color,
       edge.width=E(g)$width, edge.arrow.mode=2, edge.arrow.size=E(g)$width/10, edge.arrow.width=E(g)$width/2, 
       main=title, vertex.color=vertex.color, vertex.frame.color=vertex.frame.color, vertex.size=vertex.size,
       vertex.label.color=vertex.label.color, vertex.label.cex=vertex.label.size)
  
  par(mar=c(3,2,1,2), mgp=c(1.7,0.8,0))
  image.scale(col=colfunc(length(breaks)-1), breaks=breaks, horiz=TRUE, xlab=xlabname,ylab="")
  box()
}


MakeGraphPlotQfile <- function(tablename, edgevalues, root, inner_nodes=NULL, phenotype,
                               minsel=NULL, maxsel=NULL, minbranch=0.075){
  table <- read.table(tablename,header=TRUE)
  table <- table[which(table$branc != "Total"),]
  
  alphacols <- 3
  
  selcoefs <- table[,alphacols]
  namesselmat <- t(matrix(unlist(strsplit(as.character(table[,1]),"_")),nrow=2))
  selcoefs <- as.data.frame(cbind(namesselmat,selcoefs))
  rownames(selcoefs) <- c()
  colnames(selcoefs) <- c("child","parent","sel")  

  merged <- merge(edgevalues,selcoefs,by=c("child","parent"))
  xlabname <- "q_b statistic"

  MakeGraphPlotInterior(inputvals=merged[c(1,2,3)], selcoefs=as.numeric(as.character(merged[,4])),
                        root=root, inner_nodes=inner_nodes, title=phenotype, minsel=minsel, maxsel=maxsel,
                        xlabname=xlabname, minbranch=minbranch)  
}


MakeGraphPlot <- function(tablename=NULL, alpha.mean=NULL, edgevalues, root, inner_nodes=NULL, phenotype,
                          minsel=NULL, maxsel=NULL, minbranch=0.075){
  if(!is.null(alpha.mean)){
    selcoefs <- alpha.mean
  }else{
    table <- read.table(tablename,header=TRUE)
    trace <- table[seq(1,dim(table)[1]-1,1),]
    alphacols <- grep("alpha_",colnames(table))
    selcoefs <- apply(trace[,alphacols],2,function(x){mean(x)})
    names(selcoefs) <- colnames(trace[,alphacols])
  }

  namesselmat <- t(matrix(unlist(strsplit(names(selcoefs),"_")),nrow=3))[,c(2,3)]
  selcoefs <- as.data.frame(cbind(namesselmat,selcoefs))
  rownames(selcoefs) <- c()
  colnames(selcoefs) <- c("child","parent","sel")
  
  merged <- merge(edgevalues,selcoefs,by=c("child","parent"))
  xlabname <- "selection parameter (alpha)"
  MakeGraphPlotInterior(inputvals=merged[c(1,2,3)], selcoefs=as.numeric(as.character(merged[,4])),
                        root=root, inner_nodes=inner_nodes, title=phenotype, minsel=minsel, maxsel=maxsel,
                        xlabname=xlabname, minbranch=minbranch)
  
}


AlphaBoxPlot <- function(table, titleplot, addselline=FALSE, selline){
  data_loc <- grep("step|nodevar|nodevalue|alpha|logprob|logprior|propss|acceptance",colnames(table),invert=T)
  nodes <- length(unique(sapply(colnames(table)[data_loc],function(x){gsub('[[:digit:]]+', '', x)})))
  alphacols <- which(sapply(colnames(table),function(x){grep("alpha_",x)}) == 1)
  data <- c()
  for(i in alphacols){data <- cbind(data,jitter(table[,i],0.1))}
  colnames(data) <- colnames(table)[alphacols]
  colnames(data) <- sapply(colnames(data),function(x){gsub("alpha_","",x)})
  data <- melt(as.data.frame(data))
  colnames(data) <- c("branch","alpha")
  finalplot <- ggplot(data, aes(branch, alpha)) +
                 geom_boxplot(width=0.4,fill="light blue") +
                 geom_hline(yintercept=0,colour="black") +
                 ggtitle(titleplot) +
                 theme(plot.title = element_text(hjust=0.5)) +
                 theme(axis.text.x = element_text(angle=90, hjust=1, vjust=0.5))
  if(addselline == TRUE){finalplot <- finalplot + geom_hline(yintercept=selline,colour="red")}
  return(finalplot)
}


SimulateNormWithFix <- function(parentvalue,variance){
  
  if(parentvalue == 1){ drawnvalue <- 1
  } else if(parentvalue == 0){ drawnvalue <- 0
  } else{ drawnvalue <- rnorm(1,parentvalue,sqrt(variance))
  }
  if(drawnvalue > 1){ drawnvalue <- 1
  } else if(drawnvalue < 0){ drawnvalue <- 0}
  
  return(drawnvalue)
  
}


SimulateNorm <- function(supergraph,numSNPs,meaneffect,heritability,selmode,selbranch,selalpha,startmode="beta",sampsize=NaN,effectvec=NaN,filename="simul_output.txt"){
  
  rootname <- names(which(apply(supergraph[[1]]$parents,1,sum)==0))
  nodeorder <- supergraph[[1]]$nodes
  edgevalues <- supergraph[[2]]
  leaves <- supergraph[[1]]$leaves
  
  broken_graph <- break_graph(supergraph[[1]])
  
  #print(broken_graph)
  
  
  if(length(broken_graph$admixtures) > 0){
    admnames <- t(matrix(unlist(broken_graph$admixtures),nrow=4))
    admvalues <- supergraph[[3]]
    admlogic <- TRUE
  } else{
    admnames <- NaN
    admvalues <- NaN
    admlogic <- FALSE
  }
  
  allSNPs <- t(sapply(seq(1,numSNPs), function(x){
    
    
    # Frequency at root
    if(startmode == "allmid"){
      rootvalue <- 0.5
    } else if(startmode == "range"){
      rootvalue <- runif(1,0.1,0.9)
    } else if(startmode == "beta")  {
      rootvalue <- rbeta(1,2,2)
    } else if(startmode == "unif") {
      rootvalue <- runif(1,0,1)
    }
    
    # Draw effect sizes - Loh et al. (2015)
    alphaeffect <- -0.25
    if(is.na(sum(effectvec))){
      sdeffect <- sqrt( (heritability / numSNPs) * (rootvalue*(1-rootvalue))^(1 + alphaeffect) )
      draweffect <- rnorm(1,meaneffect,sdeffect)
    } else{
      draweffect <- sample(effectvec,1)
    }
    names(draweffect) <- "effect"
    
    
    nodevalues <- t(matrix(c(rootname,rootvalue)))
    for(i in seq(1,dim(edgevalues)[1])){
      #print(i)
      
      parentname <- as.character(edgevalues[i,2])
      childname <- as.character(edgevalues[i,1])
      drift <- as.numeric(edgevalues[i,3])
      parentvalue <- as.numeric(nodevalues[which(nodevalues[,1] == parentname),2])
      variance <- parentvalue*(1-parentvalue)*drift
      #print(parentvalue)
      #print(variance)
      
      #print(nodevalues)
      #print(c(parentname,childname,drift,parentvalue,variance))
      
      # Add selection
      if(selmode == "byeffectsize"){
        if( parentname == selbranch[2] & childname == selbranch[1] ){
          parentvalue <- parentvalue + parentvalue*(1-parentvalue)*selalpha*draweffect
        }
      } else if(selmode == "byeffectsign"){
        if( parentname == selbranch[2] & childname == selbranch[1] ){
          parentvalue <- parentvalue + parentvalue*(1-parentvalue)*selalpha*sign(draweffect)
        }
      }
      
      # Check for admixture
      add = TRUE
      if(admlogic == TRUE){
        if(childname %in% admnames[,1]){
          if(childname %in% nodevalues[,1]){
            probunif <- runif(1,0,1)
            admrateline <- admnames[which(admnames[,1] == childname),]
            admratename <- admrateline[4]
            admratevalue <- admvalues[which(as.character(admvalues[,1]) == admratename),2]
            if(parentvalue == admrateline[2]){ admrate <- admratevalue
            } else{ admrate <- 1 - admratevalue }
            if(probunif < admrate){
              #print(c(x,i))
              
              #print(c(parentvalue,variance))
              
              drawnvalue <- SimulateNormWithFix(parentvalue,variance)
              
              nodevalues[which(nodevalues[,1] == childname),] <- c(childname,drawnvalue)
              add = FALSE
            } else{ add = FALSE }
          }
        }
      }
      
      if(add == TRUE){
        #print(c(parentvalue,variance))
        drawnvalue <- SimulateNormWithFix(parentvalue,variance)
        nodevalues <- rbind(nodevalues,c(childname,drawnvalue))
      }
    }
    
    nodenames <- nodevalues[,1]
    nodevalues <- as.numeric(nodevalues[,2])
    names(nodevalues) <- nodenames
    nodevalues <- nodevalues[nodeorder]
    return(c(nodevalues,draweffect))
  }))
  
  alleffects <- allSNPs[,c("effect")]
  leafSNPs <- allSNPs[,leaves]
  
  # Binomial sampling
  if( !is.na(sampsize) ){
    leafSNPs <- apply(leafSNPs, 1:2, function(x){ 
      probsamp <- x
      dercounts <- rbinom(1,sampsize,probsamp)
      return( paste(sampsize - dercounts, dercounts,sep=",") )
    })
  }
  
  output <- cbind("X","X","X",alleffects,leafSNPs)
  colnames(output)[1] <- "CHROM"
  colnames(output)[2] <- "POS"
  colnames(output)[3] <- "SNPID"
  
  write.table(output,file=filename,col.names=TRUE,row.names=FALSE,quote=FALSE,sep="\t",append=FALSE)
  
  return(paste("Simulation finished and printed to",filename,sep=" "))
}


SimulateWFWithFix <- function(parentvalue,Ne,branchnumgen,WFmode,selcoef,tsel,draweffect){
  
  if(parentvalue == 1){ drawnvalue <- 1
  } else if(parentvalue == 0){ drawnvalue <- 0
  } else{ 
    
    drawnvalue <- parentvalue
    if(WFmode == "neutral"){
      for( gen in seq(1,branchnumgen)){
        drawnvalue <- rbinom(1,2*Ne,drawnvalue)/(2*Ne)
      }
    } else if(WFmode == "byeffectsize"){
      for( gen in seq(1,tsel)){
        selterm <- (selcoef*drawnvalue^2 + selcoef*drawnvalue + drawnvalue)/(1 + 2*selcoef*drawnvalue)
        selprob <- drawnvalue + selterm*draweffect
        drawnvalue <- rbinom(1,2*Ne,selprob)/(2*Ne)
      }
      for( gen in seq(tsel+1,branchnumgen)){
        drawnvalue <- rbinom(1,2*Ne,drawnvalue)/(2*Ne)
      }
    } else if(WFmode == "byeffectsign"){
      for( gen in seq(1,tsel)){
        selprob <- (selcoef*sign(draweffect)*drawnvalue^2 + selcoef*sign(draweffect)*drawnvalue + drawnvalue)/(1 + 2*selcoef*sign(draweffect)*drawnvalue)
        drawnvalue <- rbinom(1,2*Ne,selprob)/(2*Ne)
      }
      for( gen in seq(tsel+1,branchnumgen)){
        drawnvalue <- rbinom(1,2*Ne,drawnvalue)/(2*Ne)
      }
    }
    
  }
  
  if(drawnvalue > 1){ drawnvalue <- 1
  } else if(drawnvalue < 0){ drawnvalue <- 0}
  
  return(drawnvalue)
  
}


SimulateWF <- function(supergraph,numSNPs,meaneffect,heritability,selmode,selbranch,startmode="beta",sampsize=NaN,effectvec=NaN,filename="simul_output.txt",Ne=10000,selcoef=0.001,tsel=100){
  
  rootname <- names(which(apply(supergraph[[1]]$parents,1,sum)==0))
  nodeorder <- supergraph[[1]]$nodes
  edgevalues <- supergraph[[2]]
  leaves <- supergraph[[1]]$leaves
  
  broken_graph <- break_graph(supergraph[[1]])
  
  #print(broken_graph)
  
  
  if(length(broken_graph$admixtures) > 0){
    admnames <- t(matrix(unlist(broken_graph$admixtures),nrow=4))
    admvalues <- supergraph[[3]]
    admlogic <- TRUE
  } else{
    admnames <- NaN
    admvalues <- NaN
    admlogic <- FALSE
  }
  
  allSNPs <- t(sapply(seq(1,numSNPs), function(x){
    
    
    # Frequency at root
    if(startmode == "allmid"){
      rootvalue <- 0.5
    } else if(startmode == "range"){
      rootvalue <- runif(1,0.1,0.9)
    } else if(startmode == "beta")  {
      rootvalue <- rbeta(1,2,2)
    } else if(startmode == "unif") {
      rootvalue <- runif(1,0,1)
    }
    
    # Draw effect sizes - Loh et al. (2015)
    alphaeffect <- -0.25
    if(is.na(sum(effectvec))){
      sdeffect <- sqrt( (heritability / numSNPs) * (rootvalue*(1-rootvalue))^(1 + alphaeffect) )
      draweffect <- rnorm(1,meaneffect,sdeffect)
    } else{
      draweffect <- sample(effectvec,1)
    }
    names(draweffect) <- "effect"
    
    
    nodevalues <- t(matrix(c(rootname,rootvalue)))
    for(i in seq(1,dim(edgevalues)[1])){
      #print(i)
      
      parentname <- as.character(edgevalues[i,2])
      childname <- as.character(edgevalues[i,1])
      drift <- as.numeric(edgevalues[i,3])
      parentvalue <- as.numeric(nodevalues[which(nodevalues[,1] == parentname),2])
      branchnumgen <- round(drift * 2 * Ne)
      
      #print(nodevalues)
      #print(c(parentname,childname,drift,parentvalue,variance))
      
      # Add selection
      if( parentname == selbranch[2] & childname == selbranch[1] ){
          WFmode <- selmode
      } else{ 
          WFmode <- "neutral"
      }
      
      # Check for admixture
      add = TRUE
      if(admlogic == TRUE){
        if(childname %in% admnames[,1]){
          if(childname %in% nodevalues[,1]){
            probunif <- runif(1,0,1)
            admrateline <- admnames[which(admnames[,1] == childname),]
            admratename <- admrateline[4]
            admratevalue <- admvalues[which(as.character(admvalues[,1]) == admratename),2]
            if(parentvalue == admrateline[2]){ admrate <- admratevalue
            } else{ admrate <- 1 - admratevalue }
            if(probunif < admrate){
              #print(c(x,i))
              
              #print(c(parentvalue,variance))
              drawnvalue <- SimulateWFWithFix(parentvalue,Ne,branchnumgen,WFmode,selcoef,tsel,draweffect)
              
              nodevalues[which(nodevalues[,1] == childname),] <- c(childname,drawnvalue)
              add = FALSE
            } else{ add = FALSE }
          }
        }
      }
      
      if(add == TRUE){
        #print(c(parentvalue,variance))
        drawnvalue <- SimulateWFWithFix(parentvalue,Ne,branchnumgen,WFmode,selcoef,tsel,draweffect)
        nodevalues <- rbind(nodevalues,c(childname,drawnvalue))
      }
    }
    
    nodenames <- nodevalues[,1]
    nodevalues <- as.numeric(nodevalues[,2])
    names(nodevalues) <- nodenames
    nodevalues <- nodevalues[nodeorder]
    return(c(nodevalues,draweffect))
  }))
  
  alleffects <- allSNPs[,c("effect")]
  leafSNPs <- allSNPs[,leaves]
  
  # Binomial sampling
  if( !is.na(sampsize) ){
    leafSNPs <- apply(leafSNPs, 1:2, function(x){ 
      probsamp <- x
      dercounts <- rbinom(1,sampsize,probsamp)
      return( paste(sampsize - dercounts, dercounts,sep=",") )
    })
  }
  
  output <- cbind("X","X","X",alleffects,leafSNPs)
  colnames(output)[1] <- "CHROM"
  colnames(output)[2] <- "POS"
  colnames(output)[3] <- "SNPID"
  
  write.table(output,file=filename,col.names=TRUE,row.names=FALSE,quote=FALSE,sep="\t",append=FALSE)
  
  return(paste("Simulation finished and printed to",filename,sep=" "))
}


log_beta_prior <- function(freqs,par1,par2){
    return(dbeta(freqs,par1,par2,log=TRUE))
}


ChiSquaredReduced <- function(graphedges,contribmat,Fmat,leaves_freqs,effects,total=FALSE,randomize=FALSE){

  checkseg <- which( apply(leaves_freqs,1,sum)/dim(leaves_freqs)[2] < 0.99  & apply(leaves_freqs,1,sum)/dim(leaves_freqs)[2] > 0.01 )
  leaves_freqs <- leaves_freqs[checkseg,]
  effects <- effects[checkseg]

  # Randomize effects if necessary
  if(randomize == TRUE){effects <- effects * sample(c(-1,1),length(effects),replace=TRUE)}

  # Compute mean genetic values
  meangen <- apply(leaves_freqs * effects, 2, function(x){sum(x)})

  # Scale by average genetic value
  meangen <- (meangen - mean(meangen))

  # Compute the estimated ancestral genetic variance over all populations
  meanfreqs <- apply(leaves_freqs,1,mean)

  varmean <- sum(sapply(seq(1,length(meanfreqs)),function(i){
  score = meanfreqs[i]*(1-meanfreqs[i])*effects[i]^2
  return(score)
  }))

  if(total == FALSE){
  contribmat <- contribmat[,names(meangen)]
  }

  if(total == FALSE){
  # Compute Q_B test statistic
  Qteststat <- apply(contribmat,1, function(x){
    #print(varmean)
    lambda <- varmean * (t(x) %*% Fmat %*% x)
    numerator <- (meangen %*% x)^2
    final <- numerator / (lambda)
    return(final)
  } )

  # Compute q_B test statistic
  qteststat <- apply(contribmat,1, function(x){
    #print(varmean)
    lambda <- varmean * (t(x) %*% Fmat %*% x)
    numerator <- (meangen %*% x)
    final <- numerator / sqrt(lambda)
    return(final)
  } )

  branchorder <- apply(graphedges,1,function(x){paste(as.character(x[1]),as.character(x[2]),sep="_")})
  Qteststat <- Qteststat[branchorder]
  Pval <- 1 - pchisq(Qteststat,1)
  qteststat <- qteststat[branchorder]
  allstats <- cbind(Qteststat,qteststat,Pval)

  }

  if(total == TRUE){

    # Compute Q_X statistic
    numerator <- t(meangen) %*% solve(Fmat) %*% meangen
    denominator <- varmean
    Qteststat <- numerator / denominator

    Pval <- 1 - pchisq(Qteststat,qr(Fmat)$rank)
    allstats <- c(Qteststat,NaN,Pval)
  }

  return(allstats)
}

##########################
PolyGraph <- function(tracefile, leaves_counts, neut_leaves_counts, effects, runmode, branchcandidate, numsteps, numsample,
                      innerfreqs_proposize, alpha_prior_stdev, alpha_proposize, ssfactor, qbfactor, rootprior="unif", pg.tree){
  supergraph <- pg.tree$supergraph

  #print(branchcandidate) 
  print("Running MCMC...")
  
  # Run mode
  #runmode <- "neutral"; branchcandidate <- NaN
  #runmode <- "alphas"; branchcandidate <- NaN
  #runmode <- "onealpha"; branchcandidate <- c("v","q")

  # Set up prior parameters
  innerfreqs_prior_lower <- 0
  innerfreqs_prior_upper <- 1
  alpha_prior_mean <- 0

  # Input
  input_graph <- supergraph
  edgevalues <- supergraph[[2]]
  input_leaf_counts <- leaves_counts
  input_leaf_freqs <- ObtainFreqs(input_leaf_counts)
  input_leaf_der <- as.matrix(as.data.frame(apply(input_leaf_counts,c(1,2),function(x){as.numeric(strsplit(x,",")[[1]][2])})))
  input_leaf_tot <- as.matrix(as.data.frame(apply(input_leaf_counts,c(1,2),function(x){
                      splitted <- strsplit(x,",")[[1]]; return(as.numeric(splitted[1])+as.numeric(splitted[2]))
                    })))
  numSNPs <- dim(input_leaf_freqs)[1]

  # Neutral SNPs for F matrix
  neut_leaf_freqs <- ObtainFreqs(neut_leaves_counts)

  # Break input graph
  broken_graph <- break_graph(input_graph[[1]])
  num_inner_nodes <- length(broken_graph$inner_nodes)
  num_nonmig_edges <- length(broken_graph$edges)
  num_mig_edges <- length(broken_graph$admixtures)*2
  num_total_edges <- num_nonmig_edges + num_mig_edges

  # Starting values - inner freqs
  start_innerfreqs <- jitter(cbind( input_leaf_freqs ,t(sapply(seq(1,numSNPs), function(x) {AncestralAverage(input_leaf_freqs[x,],input_graph)}))),0.5)
  start_innerfreqs <- RemoveFixed(start_innerfreqs)
  if(dim(start_innerfreqs)[1] == 1){start_innerfreqs <- t(start_innerfreqs)}

  # Starting values - alphas
  start_alphas <- cbind(as.character(edgevalues[,1]),as.character(edgevalues[,2]),rep(0,dim(input_graph[[2]])[1]))
  colnames(start_alphas) <- c("child","parent","value")
  num_alphas <- dim(start_alphas)[1]

  # Spike-and-slab proportion (assuming only sparse selected branches)
  propss_dev <- 0.1
  minpropss <- 0.6
  maxpropss <- 0.8
  propss <- (minpropss + maxpropss)/2
  old_propss <- propss
  
  # Deconstruct graph
  graphedges <- supergraph[[2]]
  deconsgraph <- DeconstructGraph(input_graph)

  # Compute chi-squared statistics
  teststats <- ChiSquared(supergraph,input_leaf_freqs,effects,neut_leaf_freqs,total=FALSE)
  Qstat <- teststats[,1]

  # Initialize MCMC
  old_innerfreqs <- start_innerfreqs
  old_alphas <- start_alphas

  # Set-up initial probabilities
  log_prob_old_binom <- pbinom(input_leaf_der,input_leaf_tot,old_innerfreqs)
  log_prob_old_graph <- pgraph(deconsgraph,old_innerfreqs,effects,old_alphas)

  # Root frequency prior
  if(rootprior == "unif"){ log_prob_old_innerfreqs <- 0
  } else if(rootprior == "beta"){ log_prob_old_innerfreqs <- log_beta_prior(old_innerfreqs[,which(colnames(old_innerfreqs) == "r")],2,2)}
    
  # Alpha prior
  log_prob_old_alphas <- log_alphas_prior_spikeslab(as.numeric(old_alphas[,3]),alpha_prior_mean,alpha_prior_stdev,ssfactor,old_propss)

  numaccepted_freqs <- 0
  totalmoves_freqs <- 0
  numaccepted_alphas <- 0
  totalmoves_alphas <- 0
  acceptance_rate_freqs <- NaN
  acceptance_rate_alphas <- NaN
  genvaluesvec <- c()

  # Make header for trace file
  header <- "step"
  header <- c(header,as.vector(sapply(c(broken_graph$leaves,broken_graph$inner_nodes), function(x) {return( paste(x,seq(1,numSNPs,1),sep="") )})))
  header <- c(header,sapply(colnames(old_innerfreqs),function(x){paste("nodevalue",x[1],sep="_")}))
  header <- c(header,sapply(colnames(old_innerfreqs),function(x){paste("nodevar",x[1],sep="_")}))
  header <- c(header,apply(old_alphas[,c(1,2)],1,function(x){paste("alpha",x[1],x[2],sep="_")}))
  header <- c(header,"logprob","logprior")
  header <- c(header,"propss")
  header <- c(header,"acceptance_rate_freqs")
  header <- c(header,"acceptance_rate_alphas")
  header <- paste(as.vector(header),collapse="\t")
  write(header,file=tracefile,append=FALSE)

  # RUN MCMC
  for( i in seq(1,numsteps)){

    # Propose new frequencies
    new_innerfreqs <- propose_innerfreqsB(old_innerfreqs,innerfreqs_proposize,numSNPs)
    colnames(new_innerfreqs) <- colnames(old_innerfreqs)
    
    freqtranslogdiff <- translogdiff_innerfreqs(old_innerfreqs,new_innerfreqs,innerfreqs_proposize,numSNPs)

    # Evaluate likelihood of new frequencies
    log_prob_new_binom <- pbinom(input_leaf_der,input_leaf_tot,new_innerfreqs)
    log_prob_new_graph <- pgraph(deconsgraph,new_innerfreqs,effects,old_alphas)
  
    # Root frequency prior
    if(rootprior == "unif"){ log_prob_new_innerfreqs <- 0
    } else if(rootprior == "beta"){ log_prob_new_innerfreqs <- log_beta_prior(new_innerfreqs[,which(colnames(new_innerfreqs) == "r")],2,2)}
    
    log_prob_trans_freq <- apply(freqtranslogdiff, 1, function(y){return(sum(y))})
    log_prob_new <- log_prob_new_binom + log_prob_new_graph + log_prob_new_innerfreqs
    log_prob_old <- log_prob_old_binom + log_prob_old_graph + log_prob_old_innerfreqs
    
    logpostdiff <-  log_prob_new - log_prob_old + log_prob_trans_freq
    logunifsample <- log(runif(numSNPs,0,1))

    replacevec <- (logpostdiff > logunifsample)

    # Update frequencies
    old_innerfreqs[which(replacevec),] <- new_innerfreqs[which(replacevec),]
    log_prob_old_binom[which(replacevec)] <- log_prob_new_binom[which(replacevec)]
    log_prob_old_graph[which(replacevec)] <- log_prob_new_graph[which(replacevec)]
    log_prob_old_innerfreqs <- log_prob_new_innerfreqs
    numaccepted_freqs <- numaccepted_freqs + sum(replacevec)
    totalmoves_freqs <- totalmoves_freqs + numSNPs

    acceptance_rate_freqs <- numaccepted_freqs / totalmoves_freqs
  
  
    if(runmode != "neutral"){
    
      # Propose new alphas
      if(runmode == "alphas"){
        new_alphas <- propose_alphasB(old_alphas,alpha_proposize,Qstat,qbfactor)
      } else if(runmode == "onealpha"){
        alphaidx <- which(old_alphas[,1] == branchcandidate[1] & old_alphas[,2] == branchcandidate[2])
        alphatochange <- old_alphas[alphaidx,]
        alphatochange <- matrix(alphatochange,ncol=3)
        new_candidate_alpha <- propose_alphas(alphatochange,alpha_proposize)
        new_alphas <- old_alphas
        new_alphas[alphaidx,] <- new_candidate_alpha
      }

      # Evaluate alpha move
      log_prob_new_graph <- pgraph(deconsgraph,old_innerfreqs,effects,new_alphas)
      log_prob_new_alphas <- log_alphas_prior_spikeslab(as.numeric(new_alphas[,3]),alpha_prior_mean,alpha_prior_stdev,ssfactor,old_propss)
      log_prob_new <- sum(log_prob_old_binom) + sum(log_prob_new_graph) + log_prob_new_alphas
      log_prob_old <- sum(log_prob_old_binom) + sum(log_prob_old_graph) + log_prob_old_alphas
      logpostdiff <- log_prob_new - log_prob_old
      unifsample <- runif(1,0,1)
      logunifsample <- log(runif(1,0,1))
      replace <- (logpostdiff > logunifsample)
      if(replace == TRUE){
        old_alphas <- new_alphas
        log_prob_old_graph <- log_prob_new_graph
        log_prob_old_alphas <- log_prob_new_alphas
        log_prob_old <- log_prob_new
      }
      numaccepted_alphas <- numaccepted_alphas + sum(replace)
      totalmoves_alphas <- totalmoves_alphas + 1
      acceptance_rate_alphas <- numaccepted_alphas / totalmoves_alphas
    }
  
    # Propose propss
    new_propss <- rtnorm(1,old_propss,propss_dev,lower=minpropss,upper=maxpropss)
  
    # Calculate propss transition log difference
    propsstranslogdiff <- log(dtnorm(old_propss,new_propss,propss_dev,lower=minpropss,upper=maxpropss)) - log(dtnorm(new_propss,old_propss,propss_dev,lower=minpropss,upper=maxpropss))
  
    #Evaluate propss move (hyperprior for propss is Unif[0,1], so no need to calculate hyperprior difference)
    log_prob_alphas_old_propss <- log_alphas_prior_spikeslab(as.numeric(old_alphas[,3]),alpha_prior_mean,alpha_prior_stdev,ssfactor,old_propss)
    log_prob_alphas_new_propss <- log_alphas_prior_spikeslab(as.numeric(old_alphas[,3]),alpha_prior_mean,alpha_prior_stdev,ssfactor,new_propss)
    logpropssdiff <- log_prob_alphas_new_propss  - log_prob_alphas_old_propss + propsstranslogdiff
    logunifsample <- log(runif(1,0,1))
    replace <- (logpropssdiff > logunifsample)
    if(replace == TRUE){
      old_propss <- new_propss
    }
  
    # Record current prior
    log_old_prior <- sum(log_prob_old_innerfreqs) + sum(log_prob_old_alphas)

    # Print out values
    genvalues <- apply(old_innerfreqs,2,function(x){sum(2*x*effects)})
    genvar <- apply(old_innerfreqs,2, function(x){sum(2*x*(1-x)*effects^2)})
    finalvec <- c(i,as.vector(old_innerfreqs),genvalues,genvar,as.numeric(old_alphas[,3]),sum(log_prob_old_graph)+sum(log_prob_old_binom),log_old_prior,old_propss,acceptance_rate_freqs,acceptance_rate_alphas)
    if( (i %% numsample) == 0){
      write(paste(finalvec,collapse="\t"),file=tracefile,append=TRUE)
      #print(finalvec)
    }
    #print(c(i,genvalues,genvar,as.numeric(old_alphas[,3]),sum(log_prob_old_graph)+sum(log_prob_old_binom),log_old_prior,old_propss,acceptance_rate_freqs,acceptance_rate_alphas))
  } # end of MCMC
}


#################################
calc_qstat <- function(gwasfile, neutfile, qfile, pg.tree){
  leaves <- pg.tree$leaves
  supergraph <- pg.tree$supergraph

  # Load GWAS data  
  print("Loading data...")
  filename <- gwasfile
  data <- LoadCounts(filename, leaves)
  leaves_counts <- as.data.frame(data[,seq(5,dim(data)[2])])
  leaves_freqs <- ObtainFreqs(leaves_counts)
  effects <- as.numeric(data[,4])
  
  # Load Neutral data
  neutfilename <- neutfile
  neutdata <- LoadCounts(neutfilename, leaves)
  neut_leaves_counts <- as.data.frame(neutdata[,seq(5,dim(neutdata)[2])])
  neut_leaves_freqs <- ObtainFreqs(neut_leaves_counts)

  # Deconstruct graph
  graphedges <- supergraph[[2]]
  deconsgraph <- DeconstructGraph(supergraph)
  leaves <- supergraph[[1]]$leaves

  # Compute empirical covariance matrix
  checksegneut <- which( apply(neut_leaves_freqs,1,sum)/dim(neut_leaves_freqs)[2] < 0.99  & apply(neut_leaves_freqs,1,sum)/dim(neut_leaves_freqs)[2] > 0.01 )
  neut_leaves_freqs <- neut_leaves_freqs[checksegneut,]

  # Compute neutral hetero
  neut_leaves_freqs_means <- apply(neut_leaves_freqs, 1, mean)
  mean_hetero <- neut_leaves_freqs_means*(1-neut_leaves_freqs_means)
  numSNPs <- length(neut_leaves_freqs_means)

  # Compute F matrix
  print("Computing F matrix...")
  Fmat <- sapply(seq(1,dim(neut_leaves_freqs)[2]),function(x){
       sapply(seq(1,dim(neut_leaves_freqs)[2]),function(y){
	cov(neut_leaves_freqs[,x]/sqrt(mean_hetero), neut_leaves_freqs[,y]/sqrt(mean_hetero))
	})
       })
  colnames(Fmat) <- colnames(neut_leaves_freqs)
  rownames(Fmat) <- colnames(neut_leaves_freqs)

  # Compute contributions of each branch to each leaf
  contribmat <- c()
  contribmat_names <- c()
  for(branchidx in seq(1,dim(graphedges)[1])){
    targetbranch <- c(as.character(graphedges[branchidx,1]),as.character(graphedges[branchidx,2]))
    name <- paste(targetbranch,collapse="_")
    branchvec <- CollectBranches(supergraph,targetbranch,deconsgraph,leaves)
    contribmat <- rbind(contribmat,branchvec)
    contribmat_names <- rbind(contribmat_names,name)
  }
  rownames(contribmat) <- contribmat_names
  colnames(contribmat) <- leaves

  # Standardize vectors to be of unit length
  contribmat <- t(apply(contribmat,1, function(x) {
    x <- x - mean(x)
    return(x / norm_vec(x))
  }))

  # Calculate chi-squared statistics
  print("Computing Q_b statistics...")
  stats <- ChiSquaredReduced(graphedges,contribmat,Fmat,leaves_freqs,effects,total=FALSE,randomize=FALSE)
  qtab <- cbind(rownames(stats),round(stats[,1],3),round(stats[,2],3),stats[,3])
  colnames(qtab) <- c("branch","Q_B","q_B","Pval")
  totalstat <- ChiSquaredReduced(graphedges,contribmat,Fmat,leaves_freqs,effects,total=TRUE,randomize=FALSE)
  qtab <- rbind(qtab, cbind("Total",round(totalstat[1],3),round(totalstat[2],3),totalstat[3]))

  # For later use
  pvaltotal <- totalstat[3]
    
  # Calculate sign-randomized chi-squared statistics
  pseudorep <- 1000
  print(paste("Computing sign-randomized P-values, using ",pseudorep," pseudo-replicates...",sep=""))
  stats <- ChiSquaredReduced(graphedges,contribmat,Fmat,leaves_freqs,effects,total=FALSE,randomize=TRUE)
  totalstat <- ChiSquaredReduced(graphedges,contribmat,Fmat,leaves_freqs,effects,total=TRUE,randomize=TRUE)
  allstats <- c( round(stats[,1],3), round(totalstat[1],3) )
  names(allstats)[length(names(allstats))] <- "Total"
  randqtab <- allstats
  for(i in seq(2,pseudorep)){
    stats <- ChiSquaredReduced(graphedges,contribmat,Fmat,leaves_freqs,effects,total=FALSE,randomize=TRUE)
    totalstat <- ChiSquaredReduced(graphedges,contribmat,Fmat,leaves_freqs,effects,total=TRUE,randomize=TRUE)
    allstats <- c( round(stats[,1],3), round(totalstat[1],3) )
    randqtab <- cbind(randqtab, allstats)
  }
  
  # Calculate sign-randomized p-values
  randpvals <- sapply( seq(1, length(as.numeric(qtab[,2]))), function(i) { sum(randqtab[i,] > as.numeric(qtab[,2])[i]) / dim(randqtab)[2] } )
  qtab <- cbind(qtab,randpvals)
  colnames(qtab)[length(colnames(qtab))] <- "SignRandPval"
  
  # Write chi-squared statistics table
  write.table(qtab,file=qfile,quote=FALSE,col.names=TRUE,row.names=FALSE)
}
```
