## Supplementary Script for "Exploratory analysis of multiple traits co-adaptations in the population history": poplar_pop25_analysis20211002.R

```
# Copyright (C) Reiichiro Nakamichi
# License: http://www.gnu.org/licenses/gpl.html GPL version 2 or higher


# for main analysis
library(FinePop2)
source("FinePop2Ext20211002.R")
library(gtools)
library(MASS)
library(sf)
library(shape)
library(foreach)
library(doParallel)

# for Kernel GWAS
library(ttutils)
library(methods)
library(psych)
library(EQL)
library(mvtnorm)
library(Matrix)
library(KRLS)
library(kernlab)
source("Omegafunction20210927.R")

# for TREEMIX
library(RColorBrewer)
source("TreemixFunction20211001.R")

# for PolyGraph
library(msm)
library(reshape2)
library(pscl)
library(ggplot2)
source("admixturegraphFunction20210927.R")
source("PolygraphFunction20211001.R")


#################################
### data load (individual)
# genotype
gt.ind <- read.csv("data_poplar_genotype_30k_ind445.csv", row.names=1)

# environment
geo.ind <- read.csv("data_poplar_geo_ind445.csv", row.names=1)
soil.ind <- read.csv("data_poplar_soil_ind445.csv", row.names=1)

# trait
trait.ind <- read.csv("data_poplar_trait_ind445.csv", row.names=1)

# sampled area of individuals
area.ind <- read.csv("data_poplar_sample_area_ind445.csv", row.names=1)

# gene-snp list, annotation 
info.snp.gene <- read.csv("data_poplar_snps_genes_30k.csv")
rownames(info.snp.gene) <- info.snp.gene[,"SNP"]
info.gene <- read.csv("data_poplar_annot_genes_3k.csv")
rownames(info.gene) <- info.gene[,"GENE"]
info.gene.entrez <- read.csv("data_poplar_annot_genes_3k_used.csv")
rownames(info.gene.entrez) <- info.gene.entrez[,"GENE"]


# North America map
poplarmap <- read_sf("data_poplar_map/poplarmap.shp")


### map (individual)
x11()
par(mar=c(2.5,2.5,1,1))
plot(poplarmap$geometry, axes=T, lwd=0.1, border="gray80", col="gray80", xlim=c(-140,-115), ylim=c(44,60))
text(-geo.ind[,"LON"], geo.ind[,"LAT"], labels=area.ind[,"label"], col=area.ind[,"areaID"])
box()


### data csreeming ###
# ind neighbours
n.neighbour <- 3
ind.dist <- as.matrix(dist(scale(geo.ind[,c("LAT","LON","ALT")])))
ind.neighbour <- list()
for(ci in rownames(geo.ind)){
  cdist <- ind.dist[,ci]
  cdist <- cdist[names(cdist)!=ci]
  cdist.select <- (cdist==0)|(order(cdist) <= n.neighbour)
  ind.neighbour[[ci]] <- names(cdist[cdist.select])
}

# trait year average
trait.set <- colnames(trait.ind)
trait.set <- substr(trait.set, 1, nchar(trait.set)-2)
trait.set[1] <- "ADd"
trait.set[2] <- "ABd"
trait.set[5] <- "Ramets"
trait.set <- gsub("_", "", trait.set)

traitY.ind <- array(NA, c(nrow(trait.ind),length(unique(trait.set))))
dimnames(traitY.ind) <- list(rownames(trait.ind), unique(trait.set))
for(ct in unique(trait.set)){
  ctrait <- trait.ind[,grep(ct, colnames(trait.ind)), drop=F]
  traitY.ind[,ct] <- rowMeans(ctrait, na.rm=T)
}
traitY.ind[is.nan(traitY.ind)] <- NA
rm(trait.set)

# trait, NA cutoff & imputation
traitYI.ind <- traitY.ind
for(ci in rownames(traitYI.ind)){
  ci.n <- ind.neighbour[[ci]]
  ci.nmean <- colMeans(traitY.ind[ci.n,,drop=F], na.rm=T)
  na.ct <- is.na(traitYI.ind[ci,])
  traitYI.ind[ci,na.ct] <- ci.nmean[na.ct]
}
cn.na.trait <- apply(traitYI.ind, 2, function(x){sum(is.na(x))})
table(cn.na.trait)
traitYI.ind <- traitYI.ind[,which(cn.na.trait==0)]
rm(cn.na.trait)

# create pop data
area.pop <- unique(area.ind)
rownames(area.pop) <- NULL
area.pop <- area.pop[order(area.pop[,"areaID"]),]
rownames(area.pop) <- NULL

make_data_pop <- function(data.ind, pop.list, pop.assign){
  data.pop <- array(NA, c(length(pop.list),ncol(data.ind)), dimnames=list(pop.list,colnames(data.ind)))
  for(cpop in pop.list){
    cind <- which(pop.assign==cpop)
    cdata <- colMeans(data.ind[cind,,drop=F], na.rm=T)
    data.pop[cpop,] <- cdata
  }
  return(data.pop)
}

traitYI.pop <- make_data_pop(traitYI.ind, area.pop[,"label"], area.ind[,"label"])
geo.pop <- make_data_pop(geo.ind, area.pop[,"label"], area.ind[,"label"])
soil.pop <- make_data_pop(soil.ind, area.pop[,"label"], area.ind[,"label"])
maf.pop <- make_data_pop(gt.ind/2, area.pop[,"label"], area.ind[,"label"])

# remove size 1 pop
pop.size1 <- sort(as.integer(names(which(table(area.ind[,"areaID"])==1))))
pop.size1 <- area.pop[match(pop.size1, area.pop[,"areaID"]),"label"]

rm.pop.size1pop <- match(pop.size1, area.pop[,"label"])
traitYI.pop <- traitYI.pop[-rm.pop.size1pop,]
geo.pop <- geo.pop[-rm.pop.size1pop,]
soil.pop <- soil.pop[-rm.pop.size1pop,]
maf.pop <- maf.pop[-rm.pop.size1pop,]
area.pop <- area.pop[-rm.pop.size1pop,]; rownames(area.pop) <- NULL

rm.ind.size1pop <- match(pop.size1, area.ind[,"label"])
gt.ind <- gt.ind[-rm.ind.size1pop,]
geo.ind <- geo.ind[-rm.ind.size1pop,]
soil.ind <- soil.ind[-rm.ind.size1pop,]
trait.ind <- trait.ind[-rm.ind.size1pop,]
traitYI.ind <- traitYI.ind[-rm.ind.size1pop,]
area.ind <- area.ind[-rm.ind.size1pop,]

# remove pop NA loci
rm.loci.popNA <- colSums(is.na(maf.pop))>0
maf.pop <- maf.pop[,!rm.loci.popNA]
gt.ind <- gt.ind[,!rm.loci.popNA]
name.rm.loci.popNA <- names(which(rm.loci.popNA))
info.snp.gene <- info.snp.gene[setdiff(rownames(info.snp.gene), name.rm.loci.popNA),]
rm(rm.loci.popNA, name.rm.loci.popNA)


### map (population)
x11();par(mar=c(2.5,2.5,1,1))
plot(poplarmap$geometry, axes=T, lwd=0.1, border="gray80", col="gray80", xlim=c(-140,-115), ylim=c(44,60))
text(-geo.pop[,"LON"], geo.pop[,"LAT"], labels=area.pop[,"label"])
box()


### correpondense analysis ###
draw_crp <- function(geo, soil, trait, maf, sample.label, xlim=NULL, ylim=NULL){
  maf <- maf[,!(colSums(is.na(maf))>0)]
  crp.data <- cbind(geo, soil, trait, maf)
  crp.data <- apply(crp.data, 2, function(x){(x-min(x))/(max(x)-min(x))})
  res.crp <- corresp(crp.data, nf=2)

  i.geo <- match(colnames(geo),colnames(crp.data))
  i.soil <- match(colnames(soil),colnames(crp.data))
  i.trait <- match(colnames(trait),colnames(crp.data))

  plot(res.crp$cscore, type="n", xlim=xlim, ylim=ylim, xlab="1st axis", ylab="2nd axis")
  points(res.crp$cscore, pch=19, col="gray", cex=0.1)
  text(res.crp$rscore, labels=sample.label, col="blue")
  points(res.crp$cscore[i.geo,], col="green", pch=19)
  text(res.crp$cscore[i.geo,], labels=colnames(geo))
  points(res.crp$cscore[i.soil,], col="orange", pch=19)
  text(res.crp$cscore[i.soil,], labels=colnames(soil))
  points(res.crp$cscore[i.trait,], col="cyan", pch=19)
  text(res.crp$cscore[i.trait,], labels=colnames(trait), col="red")
  segments(x0=c(-0.2, 0), y0=c(0, -0.2), x1=c(0.2, 0), y1=c(0, 0.2))
  return(res.crp)
}

# individual
x11() 
res.crp.ind <- draw_crp(geo.ind, soil.ind, traitYI.ind, gt.ind/2, area.ind[,"label"])

# population
x11(); par(mar=c(3,3,0.2,0.2), mgp=c(2,0.7,0))
res.crp.pop <- draw_crp(geo.pop, soil.pop, traitYI.pop, maf.pop, area.pop[,"label"], xlim=c(-4.5,2.6), ylim=c(-3,4.1))


### GENEPOP format / FST analysis ###
make_genepopfile <- function(gt, pop.list, pop.assign, filestem){
  genepop.file <- paste0(filestem, "_genepop.txt")
  popname.file <- paste0(filestem, "_popname.txt")
  gt.gp <- array("", dim(gt), dimnames=dimnames(gt))
  gt.gp[gt==0] <- "0101"
  gt.gp[gt==1] <- "0102"
  gt.gp[gt==2] <- "0202"
  gt.gp[is.na(gt)] <- "0000"
  outtext <- c("#workLLE", colnames(gt.gp))
  for(cp in pop.list){
    cgt <- gt.gp[pop.assign==cp,,drop=F]
    ciname <- rownames(cgt)
    cgt <- apply(cgt,1,paste0,collapse="\t")
    cblock <- c("POP",paste0(ciname, ",\t", cgt))
    outtext <- c(outtext, cblock)
  }
  cat(outtext, file=genepop.file, sep="\n")
  cat(pop.list, sep="\n", file=popname.file)
}

### pop 25, all 30k SNPs
make_genepopfile(gt.ind, area.pop[,"label"], area.ind[,"label"], "pop25_snp30k")
popdata.pop25.snp30k <- read.GENEPOP("pop25_snp30k_genepop.txt", "pop25_snp30k_popname.txt")

# pop-specific FST
res.psFST.pop25.snp30k <-  pop_specificFST(popdata.pop25.snp30k)
psFST.pop25.snp30k <- res.psFST.pop25.snp30k$fst[,"FST"]
names(psFST.pop25.snp30k) <- popdata.pop25.snp30k$pop_names

plot.psfst <- function(res.psfst, order.psfst=NULL, title=NULL, ...){
  if(is.null(order.psfst)){order.psfst <- 1:nrow(res.psfst$fst)}
  y.psfst <- res.psfst$fst[order.psfst,"FST"]
  names(y.psfst) <- rownames(res.psfst$fst)[as.integer(order.psfst)]
  se.psfst <- res.psfst$fst[order.psfst,"SE"]
  max.y <- max(y.psfst+se.psfst*2)
  min.y <- min(0, min(y.psfst-se.psfst*2))
  x <- barplot(y.psfst, las=3, ylim=c(min.y, max.y), main=title, ...)
  segments(x0=x, x1=x, y0=y.psfst-se.psfst*1.96, y1=y.psfst+se.psfst*1.96)
}
x11(); par(mar=c(4, 3, 0.2, 0.2), mgp=c(2, 0.5, 0))
plot.psfst(res.psFST.pop25.snp30k, order(psFST.pop25.snp30k), xlab="Population", ylab="Population Specific FST")


# locus-specific FST (30k SNPs)
lsFST.pop25.snp30k <- locus_specificFST(popdata.pop25.snp30k)
names(lsFST.pop25.snp30k) <- popdata.pop25.snp30k$loci_names

### pop 25, gene-tag SNP 3k (select representative SNP by lsFST)
list.gene <- unique(info.snp.gene[,"GENE"])
list.gene_snp <- list()
for(cg in list.gene){list.gene_snp[[cg]] <- info.snp.gene[info.snp.gene[,"GENE"]==cg,"SNP"]}

select.gene_snp <- array("", c(length(list.gene_snp), 1), dimnames=list(names(list.gene_snp), "lsFST"))
for(cgi in 1:length(list.gene_snp)){
  cg <- names(list.gene_snp)[cgi]
  csnps <- list.gene_snp[[cg]]
  cfst <- lsFST.pop25.snp30k[csnps]
  select.gene_snp[cg,"lsFST"] <- names(cfst[order(cfst, decreasing=T)[1]])
}

make_genepopfile(gt.ind[,select.gene_snp[,"lsFST"]], area.pop[,"label"], area.ind[,"label"], "pop25_snp3k")
popdata.pop25.snp3k <- read.GENEPOP("pop25_snp3k_genepop.txt", "pop25_snp3k_popname.txt")

# locus-specific FST (3k SNPs)
lsFST.pop25.snp3k <- locus_specificFST(popdata.pop25.snp3k)
names(lsFST.pop25.snp3k) <- popdata.pop25.snp3k$loci_names

### pop 25, highly differentiated gene-tag 0.5k SNPs ( p.gamma(lsFST) < 0.1 )
pv.gamma.cutoff <- 0.1
gammafit.gfst <- fitdistr(x=lsFST.pop25.snp3k, densfun="gamma")
gs <- gammafit.gfst$estimate["shape"]
gr <- gammafit.gfst$estimate["rate"]
pv.lsfst <- pgamma(lsFST.pop25.snp3k, shape=gs, rate=gr, lower.tail=F)
snp.high.gfst <- pv.lsfst < pv.gamma.cutoff
sum(snp.high.gfst)

x11()
h <- hist(lsFST.pop25.snp3k, nclass=100, probability=T, xlab="ls-gFST", main="Histogram of ls-gFST (3k genes)")
curve(dgamma(x,shape=gs, rate=gr), from=0, to=1, add=T, col="red")
qv <- qgamma(pv.gamma.cutoff, shape=gs, rate=gr, lower.tail=F)
#qv.b <- qgamma(1/popdata.pop25.snp3k$num_loci, shape=gs, rate=gr, lower.tail=F)
abline(v=qv,col="blue")
#abline(v=qv.b,col="darkgreen")

make_genepopfile(gt.ind[,names(which(snp.high.gfst))], area.pop[,"label"], area.ind[,"label"], "pop25_snp3kD")
popdata.pop25.snp3kD <- read.GENEPOP("pop25_snp3kD_genepop.txt", "pop25_snp3kD_popname.txt")


### Correspondence Analysis ###
data.get <- GET.data(popdata.pop25.snp3kD, cbind(geo.pop, soil.pop), traitYI.pop)
data.attr.get <- GET.data.attribute(data.get)

# correspondence analsysis
result.crp <- GET.corresp(data.get)

# assiciation, env vs trait/gene, GLS
cov.pop <- cov.pop.mafW3(popdata.pop25.snp30k)
result.gls <- GET.GLS(data.get=data.get, omega.list=cov.pop$cov.maf)
result.fdr <- GET.FDR(result.gls, fdr=0.1)

result.fdr.info <- result.fdr[,c("environment","trait_gene")]
result.fdr.info <- cbind(result.fdr.info, gene=info.snp.gene[result.fdr.info[,"trait_gene"],"GENE"])
result.fdr.info <- result.fdr.info[!is.na(result.fdr.info[,"gene"]),c("environment","gene")]
result.fdr.info <- cbind(ENVIRONMENT=result.fdr.info[,"environment"], info.gene.entrez[result.fdr.info[,"gene"],])
write.csv(result.fdr.info, file="assoc_env_genettrait.csv", row.names=F)

# plot
col.psfst <- psFST.pop25.snp30k
col.psfst <- (col.psfst-min(col.psfst))/(max(col.psfst)-min(col.psfst))
col.psfst <- rgb(1-col.psfst,0,col.psfst)

lim.xy0 <- rbind(result.crp$rscore,result.crp$cscore)
lim.xy0 <- rbind(apply(lim.xy0, 2, min),apply(lim.xy0, 2, max))
dimnames(lim.xy0) <- list(c("min","max"),c("x","y"))

data.attr.get[,"size"] <- 1.3
data.attr.get[data.attr.get[,"type"]=="gene","size"] <- 1

lim.xy <- lim.xy0
lim.xy[,"x"] <- c(-3, 2.4)
lim.xy[,"y"] <- c(-4.1, 2.9)
x11();par(mar=c(2.5, 2.5, 0.2, 0.2), mgp=c(1.5, 0.5, 0))
plot.GET.correspET(result.crp, result.fdr, data.attr.get, pop.color=col.pop.agb,
                   lim.xy=lim.xy, cex.pop=1.3)


### GWAS for PolyGraph ###
# kernel GWAS  (!!! several hours required !!!)
# Preclaculated result of kernel GWAS is in "gwas_kernel_env" folder.

gene3k <- unique(info.snp.gene[,"GENE"])
n.gene3k <- length(gene3k)
Ytrait <- apply(traitYI.ind, 2, function(x){(x-min(x))/(max(x)-min(x))})           # dependent variable
Zenv <- apply(cbind(geo.ind,soil.ind), 2, function(x){(x-min(x))/(max(x)-min(x))})�@#�@population structure�F env

kerGauM <- list()
kgwas.H.env <- array(NA, c(n.gene3k, ncol(Ytrait)), dimnames=list(gene3k, colnames(Ytrait)))
cstep <- ""
for(cgi in 1:n.gene3k){
  message(rep("\b",nchar(cstep)), appendLF=F)
  cstep <- paste0(cgi, "/", n.gene3k)
  message(cstep, appendLF=F); flush.console()
  cg <- gene3k[cgi]
  csnps <- info.snp.gene[info.snp.gene[,"GENE"]==cg,"SNP"]
  cmaf <- gt.ind[,csnps, drop=F] / 2
  cmaf[is.na(cmaf)] <- 0.0 # imputation
  cker <- gen.ker(cmaf, "Gau")
  ckerM <- cker$ker.cen / cker$v1.cen
  kerGauM[[cg]] <- ckerM
  Xcov <- cov(cmaf)
  kgwas.H.env[cg, ] <- apply(Ytrait, 2, function(x){Chisq.Ustat(x, Xcov, Zenv, ckerM)})
}
message(" done.")

kgwas.CCT.env <- tan(pi*(0.5-kgwas.H.env))
kgwas.P.env <- 1 - pcauchy(kgwas.CCT.env)
kgwas.Q.env <- apply(kgwas.P.env, 2, p.adjust, method="BH")

FDR <- 1e-2
kgwas.sig.env <- list()
for(ct in colnames(Ytrait)){
  cq <- kgwas.Q.env[,ct]
  kgwas.sig.env[[ct]] <- names(which(cq < FDR))
}

# GWAS effect/direction snp3k
gwas.effect.snp3k <- array(NA, c(n.gene3k, ncol(Ytrait)), dimnames=list(gene3k, colnames(Ytrait)))
for(ct in colnames(gwas.effect.snp3k)){
  cy <- traitYI.ind[,ct]
for(cg in rownames(gwas.effect.snp3k)){
  csnp <- select.gene_snp[cg,"lsFST"]
  cx <- gt.ind[,csnp]
  clm <- summary(lm(cy~cx))
  gwas.effect.snp3k[cg, ct] <- clm$coefficients["cx","t value"]
}}

# GWAS output for PolyGraph
popdata <- popdata.pop25.snp3k
gwas.sig <- kgwas.sig.env
gwasfile.dir <- "gwas_kernel_env"
for(cti in 1:length(gwas.sig)){
  ct <- names(gwas.sig)[cti]
  ct.gene <- gwas.sig[[ct]]
  cgwasfile <- paste0(gwasfile.dir, "/gwas_", ct, ".txt")
  outtxt <- paste(c("CHROM", "POS", "SNPID", "EFFECT", popdata$pop_names), collapse="\t")
  for(cg in ct.gene){
    csnp <- select.gene_snp[cg,"lsFST"]
    cgi <- which(popdata$loci_names==csnp)
    cg.count <- popdata$allele_count[[cgi]]
    cg.count <- apply(cg.count, 2, function(x){paste(x,collapse=",")})
    ceff <- gwas.effect.snp3k[cg, ct]
    outtxt <- c(outtxt, paste(c(NA,NA,cg,ceff,cg.count),collapse="\t"))
  }
  cat(outtxt, file=cgwasfile, sep="\n")

  cneutfile <- paste0(gwasfile.dir, "/neut_", ct, ".txt")
  outtxt <- paste(c("CHROM", "POS", "SNPID", "EFFECT", popdata$pop_names), collapse="\t") 
  for(cg in setdiff(gene3k, ct.gene)){
    csnp <- select.gene_snp[cg,"lsFST"]
    cgi <- which(popdata$loci_names==csnp)
    cg.count <- popdata$allele_count[[cgi]]
    cg.count <- apply(cg.count, 2, function(x){paste(x,collapse=",")})
    ceff <- gwas.effect.snp3k[cg, ct]
    outtxt <- rbind(outtxt, paste(c(NA,NA,cg,ceff,cg.count),collapse="\t"))
  }
  cat(outtxt, file=cneutfile, sep="\n")
}
rm(popdata, gwas.sig)


# Annotation of signif genes
for(cti in 1:length(gwas.sig)){
  ct <- names(gwas.sig)[cti]
  ct.gene <- gwas.sig[[ct]]
  cinfo <- info.gene[ct.gene, c("GENE","ArabidopsisDesc"),drop=F]
  cinfo <- apply(cinfo, 1, paste0, collapse="\t")
  cinfo <- paste0(cinfo, collapse="\n")
  cat(ct, cinfo, file="signif_gwas_kernel.txt", append=T, sep="\n")
}


### TREEMIX ###
# Preclaculated result of TREEMIX is in "treemix_pop25_snp30k_root_s27" folder.

treemixfile <- "pop25_snp30k_treemix.txt"
treemixdir <- "treemix_pop25_snp30k_root_s27"

make_treemixfile(popdata.pop25.snp30k, treemixfile)

### Gzip "treemixfile", set the zipped file in "treemixdir", and apply TREEMIX. ###
# treemix -i $zipped_treemixfile -root pop1 -o tm_mig0

tm.stem <- paste0(treemixdir, "/tm_mig5")

x11()
tm.tree <- plot_tree(tm.stem, arrow=0.1, arrow.lwd=2, scale=F, mbar=F, disp=0.0005)


### PolyGraph ###
## treemix to polygraph
pg.tree <- treemix2polygraph(tm.tree)

## MCMC
gwas_dir <- "gwas_kernel_env"
polygraph_dir <- "polygraph_pop25_snp3k_root_s27_gwasKernelEnv"

numsteps <- 30000
numsample <- 300
innerfreqs_proposize <- 0.1
alpha_prior_stdev <- 0.1
alpha_proposize <- 0.02
ssfactor <- 25
qbfactor <- 3
pvalcutoff <- 0.05
rootprior <- "unif"
runmode <- "alphas"
branchcandidate <- NaN
pvaltotal <- 0.05

nCPU <- 15

# select traits with significantly associated SNPs
traitYI.use.list.pg <- names(kgwas.sig.env)
for(cti in 1:length(traitYI.use.list.pg)){
  target_trait <- traitYI.use.list.pg[cti]
  gwasfile <- paste0(gwas_dir, "/gwas_", target_trait, ".txt")
  cgwas <- scan(gwasfile, character(), sep="\n")
  if(length(cgwas)<=2){traitYI.use.list.pg[cti] <- NA}
}
traitYI.use.list.pg <- traitYI.use.list.pg[!is.na(traitYI.use.list.pg)]

#�@qstat file
for(cti in 1:length(traitYI.use.list.pg)){
  target_trait <- traitYI.use.list.pg[cti]
  gwasfile <- paste0(gwas_dir, "/gwas_", target_trait, ".txt")
  neutfile <- paste0(gwas_dir, "/neut_", target_trait, ".txt")
  qfile <- paste0(polygraph_dir, "/qfile_", target_trait, ".txt")
  calc_qstat(gwasfile, neutfile, qfile, pg.tree)
}


# PolyGraph MCMC (!!! several hours required !!!)
# Preclaculated result of PolyGraph is in "polygraph_pop25_snp3k_root_s27_gwasKernelEnv" folder.

library(foreach)
library(doParallel)

cl <- makeCluster(nCPU)
registerDoParallel(cl)
test.par <- foreach(cti=1:length(traitYI.use.list.pg), .packages="msm") %dopar% {
  target_trait <- traitYI.use.list.pg[cti]
  gwasfile <- paste0(gwas_dir, "/gwas_", target_trait, ".txt")
  neutfile <- paste0(gwas_dir, "/neut_", target_trait, ".txt")
  mcmcfile <- paste0(polygraph_dir, "/MCMC_trace_", target_trait, ".txt")

  leaves <- pg.tree$supergraph[[1]]$leaves
  data <- LoadCounts(gwasfile, leaves)
  leaves_counts <- as.data.frame(data[,seq(5,dim(data)[2])])
  effects <- as.numeric(data[,4])
  data <- LoadCounts(neutfile, leaves)
  neut_leaves_counts <- as.data.frame(data[,seq(5,dim(data)[2])])

  test.polygraph <- PolyGraph(mcmcfile,
    leaves_counts, neut_leaves_counts, effects, runmode, branchcandidate, numsteps, numsample,
    innerfreqs_proposize, alpha_prior_stdev, alpha_proposize, ssfactor, qbfactor, rootprior, pg.tree=pg.tree)
}
stopCluster(cl)

# polygraph alpha post distribution
alpha.list <- list()
for(cti in 1:length(traitYI.use.list.pg)){
  message(cti, " ", appendLF=F);flush.console()
  target_trait <- traitYI.use.list.pg[cti]
  mcmcfile <- paste0(polygraph_dir, "/MCMC_trace_", target_trait, ".txt")
  mcmc.trace <- read.table(mcmcfile,header=TRUE)
  mcmc.trace <- mcmc.trace[seq(1,dim(mcmc.trace)[1]-1,1),]
  alphacols <- grep("alpha_",colnames(mcmc.trace))
  calpha <- mcmc.trace[,alphacols]
  alpha.list[[target_trait]] <- calpha
  selcoefs <- apply(calpha,2,function(x){mean(x)})
}
alpha.mean <- NULL
for(cti in 1:length(traitYI.use.list.pg)){
  target_trait <- traitYI.use.list.pg[cti]
  calpha <- alpha.list[[target_trait]]
  calpha <- calpha[-(1:10),]  # remove burn-in
  alpha.mean <- cbind(alpha.mean, colMeans(calpha, na.rm=T))
}
colnames(alpha.mean) <- traitYI.use.list.pg

# alpha boxplot
pdffile <- paste0(polygraph_dir, "polygraph_alpha_postdist.pdf")
pdf(pdffile, width=8, height=8)
for(cti in 1:length(traitYI.use.list.pg)){
  target_trait <- traitYI.use.list.pg[cti]
  calpha <- alpha.list[[target_trait]] 
  calpha <- calpha[-(1:10),]  # remove burn-in
  titleboxplot <- paste0("Posterior distributions of alpha: ", target_trait)
  alphacols <- which(sapply(colnames(calpha),function(x){grep("alpha_",x)}) == 1)
  data.alpha <- c()
  for(i in alphacols){data.alpha <- cbind(data.alpha,jitter(calpha[,i],0.1))}
  colnames(data.alpha) <- colnames(calpha)[alphacols]
  colnames(data.alpha) <- sapply(colnames(data.alpha),function(x){gsub("alpha_","",x)})
  data.alpha <- melt(as.data.frame(data.alpha))
  colnames(data.alpha) <- c("branch","alpha")
  plot_alpha <- ggplot(data.alpha, aes(branch, alpha)) +
                 geom_boxplot(width=0.4,fill="light blue") +
                 geom_hline(yintercept=0,colour="black") +
                 ggtitle(titleboxplot) +
                 theme(plot.title = element_text(hjust=0.5)) +
                 theme(axis.text.x = element_text(angle=90, hjust=1, vjust=0.5))
  plot(plot_alpha)
}
dev.off()


### PolyGraph alpha on TREEMIX
pdffile <- paste0("polygraph_alpha_treemix_root_s27.pdf")
pdf(pdffile, width=8, height=8)
for(cti in 1:length(traitYI.use.list.pg)){
  target_trait <- traitYI.use.list.pg[cti]
  plot_tree2(tm.stem, arrow=0.2, arrow.lwd=3, scale=F, mbar=F, lwd=3, title=target_trait, disp=0.0005,
             alpha.color=alpha2col(x=alpha.mean[,target_trait], edge=pg.tree$edge))
}
dev.off()


### Polygraph alpha PCA (TREEMIX all node)
plot.pca.alpha <- function(pcadata, pcares, axisA=1, axisB=2, title=NULL, PropVar=F, pop.label.only=F, ...){
  pca.score.label <- rownames(pcadata)
  i.alpha.pca <- grep("alpha_", pca.score.label)
  pca.score.label[i.alpha.pca] <- gsub("alpha_", "", pca.score.label[i.alpha.pca])
  if(pop.label.only){
    pca.score.label[i.alpha.pca] <- gsub("x[0-9]*", "", pca.score.label[i.alpha.pca])
    pca.score.label[i.alpha.pca] <- gsub("_", "", pca.score.label[i.alpha.pca])
  }
  pca.score.color <- rep("darkgreen", length(pca.score.label))
  pca.score.color[i.alpha.pca] <- "blue"
  is.inner.node <- pca.score.label==""

  pca.loading.label <- colnames(pcadata)
  pca.loading.color <- "red"

  pca.loading.scaled <- pcares$rotation * rep(pcares$sdev,each=ncol(pcadata)) *
                        sqrt(nrow(pcadata)/ncol(pcadata))
  pca.score.scaled <- pcares$x / rep(pcares$sdev, each=nrow(pcadata)) / sqrt(nrow(pcadata))

  propv <- summary(pcares)$importance[2, c(axisA, axisB)]
  propv <- sprintf("%.1f",propv*100)
  x.lab <- paste0("PC",axisA)
  if(PropVar){x.lab <- paste0(x.lab, " (", propv[1], "%)")}
  y.lab <- paste0("PC",axisB)
  if(PropVar){y.lab <- paste0(y.lab, " (", propv[2], "%)")}

  x.pca <- c(pca.loading.scaled[,axisA]*1.2, pca.score.scaled[,axisA])
  y.pca <- c(pca.loading.scaled[,axisB]*1.2, pca.score.scaled[,axisB])

  plot(x=x.pca, y=y.pca, type="n", xlab=x.lab, ylab=y.lab, main=title, ...)
  points(pca.score.scaled[is.inner.node,c(axisA,axisB)], col="gray60", pch=19)
  text(pca.score.scaled[,c(axisA,axisB)], pca.score.label, col=pca.score.color)
  text(pca.loading.scaled[,c(axisA,axisB)], pca.loading.label, col=pca.loading.color)
  arrows(x0=rep(0, ncol(pcadata)), y0=rep(0, ncol(pcadata)),
         x1=pca.loading.scaled[,axisA]*0.9, y1=pca.loading.scaled[,axisB]*0.9,
         col=pca.loading.color,length=0.1)
}

# cor env vs trait
cordata.env <- cbind(geo.ind, soil.ind)
cordata.trait <- traitYI.ind
cor.mat.et <- array(NA, c(ncol(cordata.env),ncol(cordata.trait)),
                   dimnames=list(colnames(cordata.env),colnames(cordata.trait)))
for(ct in colnames(cordata.trait)){
for(ce in colnames(cordata.env)){
  ctv <- cordata.trait[,ct]
  cev <- cordata.env[,ce]
  cor.mat.et[ce,ct] <- cor(ctv,cev)
}}

## pca alpha vs trait
pcadata.alpha <- alpha.mean
pcares.alpha <- prcomp(pcadata.alpha)
summary(pcares.alpha)

x11();plot.pca.alpha(pcadata.alpha, pcares.alpha, title="Polygraph alpha vs trait")

## pca alpha-env vs trait
pcadata.alphaenv <- rbind(alpha.mean, cor.mat.et[,colnames(alpha.mean)])
pcadata.alphaenv <- apply(pcadata.alphaenv.pop, 2, function(x){(x-min(x))/(max(x)-min(x))})
pcares.alphaenv <- prcomp(pcadata.alphaenv)
summary(pcares.alphaenv)

x11();plot.pca.alpha(pcadata.alphaenv, pcares.alphaenv, axisA=1, axisB=2,
                     title="Polygraph alpha-environment vs trait")

x11();par(mar=c(3, 3, 0.5, 0.5), mgp=c(1.5, 0.5, 0))  # !!! used in article !!!
plot.pca.alpha(pcadata.alphaenv, pcares.alphaenv,  axisA=1, axisB=2,
               PropVar=T, pop.label.only=T, xlim=c(-0.42,0.4))

write.csv(pcadata.alphaenv, file="test.csv")


### Polygraph alpha PCA (TREEMIX pop node only)
plot.pca.pop <- function(pcadata, pcares, col.loading=NULL, col.score=NULL, axisA=1, axisB=2, title=NULL, PropVar=F){
  pca.score.label <- rownames(pcadata)
  pca.score.color <- if(is.null(col.loading)){"blue"}else{col.loading}

  pca.loading.label <- colnames(pcadata)
  pca.loading.color <- if(is.null(col.score)){"red"}else{col.score}

  pca.loading.scaled <- pcares$rotation * rep(pcares$sdev,each=ncol(pcadata)) *
                        sqrt(nrow(pcadata)/ncol(pcadata))
  pca.score.scaled <- pcares$x / rep(pcares$sdev, each=nrow(pcadata)) / sqrt(nrow(pcadata))

  propv <- summary(pcares)$importance[2, c(axisA, axisB)]
  propv <- sprintf("%.1f",propv*100)
  x.lab <- paste0("PC",axisA)
  if(PropVar){x.lab <- paste0(x.lab, " (", propv[1], "%)")}
  y.lab <- paste0("PC",axisB)
  if(PropVar){y.lab <- paste0(y.lab, " (", propv[2], "%)")}

  plot(x=c(pca.loading.scaled[,axisA]*1.2, pca.score.scaled[,axisA]),
       y=c(pca.loading.scaled[,axisB]*1.2, pca.score.scaled[,axisB]),
       type="n", xlab=x.lab, ylab=y.lab, main=title)
  text(pca.score.scaled[,c(axisA,axisB)], pca.score.label, col=pca.score.color)
  text(pca.loading.scaled[,c(axisA,axisB)], pca.loading.label, col=pca.loading.color)
  arrows(x0=rep(0, ncol(pcadata)), y0=rep(0, ncol(pcadata)),
         x1=pca.loading.scaled[,axisA]*0.9, y1=pca.loading.scaled[,axisB]*0.9,
         col=pca.loading.color,length=0.1)
}

# alpha, pop node only
pop.names <- unique(area.ind[,"label"])
env.pop <- make_data_pop(cbind(geo.ind,soil.ind), pop.names, area.ind[,"label"])
alpha.pop <- array(NA, dim=c(length(pop.names),ncol(alpha.mean)))
dimnames(alpha.pop) <- list(pop.names, colnames(alpha.mean))
for(cp in pop.names){
  calpha <- alpha.mean[grep(paste0(cp,"_"), rownames(alpha.mean)),,drop=F]
  alpha.pop[cp,] <- colMeans(calpha)
}

# pca alpha-env vs trait 
pcadata.alphaenv.pop <- scale(cbind(alpha.pop,env.pop))
#pcadata.alphaenv.pop <- cbind(alpha.pop,env.pop)
#pcadata.alphaenv.pop <- apply(pcadata.alphaenv.pop, 2, function(x){(x-min(x))/(max(x)-min(x))})
col.alphaenv <- c(rep("red", ncol(alpha.pop)), rep("darkgreen", ncol(env.pop)))
pcares.alphaenv.pop <- prcomp(pcadata.alphaenv.pop)
summary(pcares.alphaenv.pop)

x11();par(mar=c(3, 3, 0.5, 0.5), mgp=c(1.5, 0.5, 0))
plot.pca.pop(pcadata.alphaenv.pop, pcares.alphaenv.pop, col.score=col.alphaenv, axisA=1, axisB=2, PropVar=T)


### Polygraph alpha PCA on treemix ###
# alpha (all node)
pca.score.alpha <- pcares.alpha$x
x11(); par(mfrow=c(1,2))
for(i in 1:2){
  plot_tree2(tm.stem, arrow=0.1, arrow.lwd=3, lwd=3, title=paste0("PCA", i), mbar=F, scale=F, disp=0.001,
             alpha.color=alpha2col(x=pca.score.alpha[,i], edge=pg.tree$edge))
}

# alpha-env (all node)
pca.score.alphaenv <- pcares.alphaenv$x
x11(); par(mfrow=c(1,2)); par(mar=c(3, 1, 3, 1), mgp=c(1.5, 0.5, 0))
for(i in 1:2){
  plot_tree2(tm.stem, arrow=0.1, arrow.lwd=3, lwd=3, title=paste0("PC", i), mbar=F, scale=F, disp=0.001,
             alpha.color=alpha2col(x=pca.score.alphaenv[,i], edge=pg.tree$edge))
}

# alpha (pop node only)
pca.score.alpha.pop <- pcares.alpha$x
is.pop <- rep(F, nrow(pca.score.alpha.pop))
for(cp in pop.names){
  is.pop[grep(paste0(cp,"_"), rownames(pca.score.alpha.pop))] <- T
}
pca.score.alpha.pop[!is.pop,] <- 0
x11(); par(mfrow=c(1,2)); par(mar=c(3, 1, 3, 1), mgp=c(1.5, 0.5, 0))
for(i in 1:2){
  plot_tree2(tm.stem, arrow=0.1, arrow.lwd=3, lwd=3, title=paste0("PCA", i), mbar=F, scale=F, disp=0.001,
             alpha.color=alpha2col(x=pca.score.alpha.pop[,i], edge=pg.tree$edge))
}

# alpha-env (pop node only)
pca.score.alphaenv.pop <- pcares.alphaenv$x
is.pop <- rep(F, nrow(pca.score.alphaenv.pop))
for(cp in pop.names){
  is.pop[grep(paste0(cp,"_"), rownames(pca.score.alphaenv.pop))] <- T
}
pca.score.alphaenv.pop[!is.pop,] <- 0
x11(); par(mfrow=c(1,2))
for(i in 1:2){
  plot_tree2(tm.stem, arrow=0.1, arrow.lwd=3, lwd=3, title=paste0("PCA", i), mbar=F, scale=F, disp=0.001,
             alpha.color=alpha2col(x=pca.score.alphaenv.pop[,i], edge=pg.tree$edge))
}
```
