## Supplementary Script for "Exploratory analysis of multiple traits co-adaptations in the population history": runTreemix.sh

```
#!/bin/bash
treemixdata="./pop25_snp30k_treemix.txt.gz"
#treemix -i $treemixdata -root s27 -o tm_mig0
treemix -i $treemixdata -root s27 -o tm_mig5 -m 5
#treemix -i $treemixdata -g tm_mig5.vertices.gz tm_mig5.edges.gz -root s27 -o tm_mig10 -m 5
#treemix -i $treemixdata -g tm_mig10.vertices.gz tm_mig10.edges.gz -root s27 -o tm_mig20 -m 10
```
