## Supplementary Script for "Exploratory analysis of multiple traits co-adaptations in the population history": TreemixFunction20211001.R

```
# Treemix plotting function
# Originally implemented in Pickrell & Pritchar (2021) PLOS Genetics 8(11):e1002967
# Modified by R. Nakamichi


# make Treemix input file from popdata
make_treemixfile <- function(popdata, treemix.file){
  data.treemix <- array("", c(popdata$num_loci, popdata$num_pop))
  for(cl in 1:popdata$num_loci){
    cloci <- (popdata$allele_count)[[cl]]
    cloci <- paste0(cloci[1,],",",cloci[2,])
    data.treemix[cl,] <- cloci
  }
  data.treemix <- rbind(popdata$pop_names, data.treemix)
  write.table(data.treemix, file=treemix.file, quote=F, row.names=F, col.names=F)
}

set_y_coords = function(d){
	i = which(d[,3]=="ROOT")
	y = d[i,8]/ (d[i,8]+d[i,10])
	d[i,]$y = 1-y
	d[i,]$ymin = 0
	d[i,]$ymax = 1
	c1 = d[i,7]
	c2 = d[i,9]
	ni = which(d[,1]==c1)
	ny = d[ni,8]/ (d[ni,8]+d[ni,10])
        d[ni,]$ymin = 1-y
	d[ni,]$ymax = 1
	d[ni,]$y = 1- ny*(y)

        ni = which(d[,1]==c2)
        ny = d[ni,8]/ (d[ni,8]+d[ni,10])
        d[ni,]$ymin = 0
        d[ni,]$ymax = 1-y
        d[ni,]$y = (1-y)-ny*(1-y)

	for (j in 1:nrow(d)){
		d = set_y_coord(d, j)
	}	
	return(d)
}

set_y_coord = function(d, i){
	index = d[i,1]
	parent = d[i,6]
	if (!is.na(d[i,]$y)){
		return(d)
	}
	tmp = d[d[,1] == parent,]
	if ( is.na(tmp[1,]$y)){
		d = set_y_coord(d, which(d[,1]==parent))
		tmp = d[d[,1]== parent,]
	}
	py = tmp[1,]$y
	pymin = tmp[1,]$ymin
	pymax = tmp[1,]$ymax
	f = d[i,8]/( d[i,8]+d[i,10])
	#print (paste(i, index, py, pymin, pymax, f))
	if (tmp[1,7] == index){
		d[i,]$ymin = py
		d[i,]$ymax = pymax
		d[i,]$y = pymax-f*(pymax-py)
		if (d[i,5]== "TIP"){
			d[i,]$y = (py+pymax)/2
		}
	}
	else{
		d[i,]$ymin = pymin
		d[i,]$ymax = py
		d[i,]$y = py-f*(py-pymin)
                if (d[i,5]== "TIP"){
                        d[i,]$y = (pymin+py)/2
                }	
	
	}
	return(d)
}


set_x_coords = function(d, e){
	i = which(d[,3]=="ROOT")
	index = d[i,1]
        d[i,]$x = 0
        c1 = d[i,7]
        c2 = d[i,9]
        ni = which(d[,1]==c1)
	tmpx =  e[e[,1]==index & e[,2] == c1,3]
	if (length(tmpx) == 0){
		tmp = e[e[,1] == index,]
		tmpc1 = tmp[1,2]
		if ( d[d[,1]==tmpc1,4] != "MIG"){
			tmpc1 = tmp[2,2]
		}
		tmpx = get_dist_to_nmig(d, e, index, tmpc1)
	}
	if(tmpx < 0){
		tmpx = 0
	}
        d[ni,]$x = tmpx

        ni = which(d[,1]==c2)
        tmpx =  e[e[,1]==index & e[,2] == c2,3]
	if (length(tmpx) == 0){
    		tmp = e[e[,1] == index,]
                tmpc2 = tmp[2,2]
                if ( d[d[,1]==tmpc2,4] != "MIG"){
                        tmpc2 = tmp[1,2]
                }
                tmpx = get_dist_to_nmig(d, e, index, tmpc2)
	}
        if(tmpx < 0){
                tmpx = 0
        }
        d[ni,]$x = tmpx
        
        for (j in 1:nrow(d)){
                d = set_x_coord(d, e, j)
        }
        return(d)
	print(d)
}


set_x_coord = function(d, e, i){
	index = d[i,1]
        parent = d[i,6]
        if (!is.na(d[i,]$x)){
                return(d)
        }
        tmp = d[d[,1] == parent,]
        if ( is.na(tmp[1,]$x)){
                d = set_x_coord(d, e, which(d[,1]==parent))
                tmp = d[d[,1]== parent,]
        }
	#print (paste(parent, index))
	tmpx =  e[e[,1]==parent & e[,2] == index,3]
	if (length(tmpx) == 0){
                tmp2 = e[e[,1] == parent,]
                tmpc2 = tmp2[2,2]
		#print
                if ( d[d[,1]==tmpc2,4] != "MIG"){
                        tmpc2 = tmp2[1,2]
                }
		tmpx = get_dist_to_nmig(d, e, parent, tmpc2)
	}
        if(tmpx < 0){
                tmpx = 0
        }
	d[i,]$x = tmp[1,]$x+ tmpx
	return(d)
}

plot_tree_internal = function(d, e, o = NA, cex = 1, disp = 0.005, plus = 0.005, arrow = 0.05, arrow.lwd = 1,
                              ybar = 0.01, scale = T, mbar = T, mse = 0.01, plotmig = T, plotnames = T,
                              xmin = 0, lwd = 1, font = 1, label=NULL){
	plot(d$x, d$y, axes = F, ylab = "", xlab = "Drift parameter", xlim = c(xmin, max(d$x)+plus), pch = "")
	axis(1)
	mw = max(e[e[,5]=="MIG",4])
	mcols = rev(heat.colors(150))
	for(i in 1:nrow(e)){
		col = "black"
		if (e[i,5] == "MIG"){
			w = floor(e[i,4]*200)+50
			if (mw > 0.5){
				w = floor(e[i,4]*100)+50
			}
			col = mcols[w]
			if (is.na(col)){
				col = "blue"
			}
		}
		v1 = d[d[,1] == e[i,1],]
		v2 = d[d[,1] == e[i,2],]
		if (e[i,5] == "MIG"){
			if (plotmig){
			arrows( v1[1,]$x, v1[1,]$y, v2[1,]$x, v2[1,]$y, col = col, length = arrow, lwd = arrow.lwd)
			}
		}
		else{
			lines( c(v1[1,]$x, v2[1,]$x), c(v1[1,]$y, v2[1,]$y), col = col, lwd = lwd)
		}
	}
	tmp = d[d[,5] == "TIP",]
	print(tmp$x)
	print(disp)
	if ( !is.na(o)){
		for(i in 1:nrow(tmp)){
			tcol = o[o[,1] == tmp[i,2],2]
			if(plotnames){
				#print(tmp[i,2])
                        clabel <- tmp[i,2]
                        if(!is.null(label)){clabel <- label[i]}
				text(tmp[i,]$x+disp, tmp[i,]$y, labels=clabel, adj=0, cex=cex, col=tcol, font=font)
			}
		}
	}
	else{
		if (plotnames){
            clabels <- tmp[,2]
            if(!is.null(label)){clabels <- label}
		text(tmp$x+disp, tmp$y, labels = clabels, adj = 0, cex = cex, font = font)
		}
	}
	if (scale){
	print (paste("mse", mse))
        lines(c(0, mse*10), c(ybar, ybar))
	text( 0, ybar - 0.04, lab = "10 s.e.", adj = 0, cex  = 0.8)
	lines( c(0, 0), c( ybar - 0.01, ybar+0.01))
	lines( c(mse*10, mse*10), c(ybar- 0.01, ybar+ 0.01))
	}
        if (mbar){
                mcols = rev( heat.colors(150) )
                mcols = mcols[50:length(mcols)]
                ymi = ybar+0.15
                yma = ybar+0.35
                l = 0.2
                w = l/100
                xma = max(d$x/20)
                rect( rep(0, 100), ymi+(0:99)*w, rep(xma, 100), ymi+(1:100)*w, col = mcols, border = mcols)
                text(xma+disp, ymi, lab = "0", adj = 0, cex = 0.7)
		if ( mw >0.5){ text(xma+disp, yma, lab = "1", adj = 0, cex = 0.7)}
                else{
			text(xma+disp, yma, lab = "0.5", adj = 0, cex =0.7)
		}
		text(0, yma+0.06, lab = "Migration", adj = 0 , cex = 0.6)
		text(0, yma+0.03, lab = "weight", adj = 0 , cex = 0.6)
        }	
}

set_mig_coords = function(d, e){
	for (j in 1:nrow(d)){
		if (d[j,4] == "MIG"){
			p = d[d[,1] == d[j,6],]
			c = d[d[,1] == d[j,7],]
			tmpe = e[e[,1] == d[j,1],]
			y1 = p[1,]$y
			y2 = c[1,]$y
			x1 = p[1,]$x
			x2 = c[1,]$x

			mf = tmpe[1,6]	
			if (is.nan(mf)){
				mf = 0
			}
			#d[j,]$y = (y1+y2)* mf
                        #d[j,]$x = (x1+x2) *mf
                        d[j,]$y = y1+(y2-y1)* mf
			print(paste(mf, x1, x2))
                        d[j,]$x = x1+(x2-x1) *mf
		}	

	}
	return(d)

}

get_f = function(stem){
	d = paste(stem, ".cov.gz", sep = "")
	d2 = paste(stem, ".modelcov.gz", sep = "")
	d = read.table(gzfile(d), as.is = T, comment.char = "", quote = "")
	d2 = read.table(gzfile(d2), as.is = T, comment.char = "", quote = "")
	d = d[order(names(d)), order(names(d))]
	d2 = d2[order(names(d2)), order(names(d2))]
	tmpcf = vector()
        tmpmcf = vector()
        for (j in 1:nrow(d)){
                for (k in (j+1):nrow(d)){
                        tmpcf = append(tmpcf, d[j,k])
                        tmpmcf = append(tmpmcf, d[j,k] - d2[j,k])
                }
        }
        tmpv = var(tmpmcf)/var(tmpcf)
	return(1-tmpv)

}

plot_tree = function(stem, o = NA, cex = 1, disp = 0.003, plus = 0.01, flip = vector(), arrow = 0.05, arrow.lwd=1,
                     scale = T, ybar = 0.1, mbar = T, plotmig = T, plotnames = T, xmin = 0, lwd = 1, font = 1, label=NULL){
	d = paste(stem, ".vertices.gz", sep = "")
	e = paste(stem, ".edges.gz", sep = "")
	se = paste(stem, ".covse.gz", sep = "")
	d = read.table(gzfile(d), as.is = T, comment.char = "", quote = "")
	e = read.table(gzfile(e), as.is  = T, comment.char = "", quote = "")
	if (!is.na(o)){
		o = read.table(o, as.is = T, comment.char = "", quote = "")
	}
	e[,3] = e[,3]*e[,4]
	e[,3] = e[,3]*e[,4]
	
	se = read.table(gzfile(se), as.is = T, comment.char = "", quote = "")
	m1 = apply(se, 1, mean)
	m = mean(m1)
	#m = 0
	for(i in 1:length(flip)){
		d = flip_node(d, flip[i])
	}
	d$x = "NA"
	d$y = "NA"
	d$ymin = "NA"
	d$ymax = "NA"
	d$x = as.numeric(d$x)
	d$y = as.numeric(d$y)
	d$ymin = as.numeric(d$ymin)
	d$ymax = as.numeric(d$ymax)

	d = set_y_coords(d)
	d = set_x_coords(d, e)
	print(d)
	d = set_mig_coords(d, e)
	plot_tree_internal(d, e, o = o, cex = cex, xmin = xmin, disp = disp, plus = plus, arrow = arrow,
                         arrow.lwd=arrow.lwd, ybar = ybar, mbar = mbar, mse = m, scale = scale,
                         plotmig = plotmig, plotnames = plotnames, lwd = lwd, font = font, label=label)
	return(list( d= d, e = e))
}

get_dist_to_nmig = function(d, e, n1, n2){
	toreturn = e[e[,1] == n1 & e[,2] == n2,3]
	#print(toreturn)
	while ( d[d[,1] ==n2,4] == "MIG"){
		tmp = e[e[,1] == n2 & e[,5] == "NOT_MIG",]
		toreturn = toreturn+tmp[1,3]
		n2 = tmp[1,2]
	}
	return(toreturn)
}

flip_node = function(d, n){
	i = which(d[,1] == n)
	t1 = d[i,7]
	t2 = d[i,8]
	d[i,7] = d[i,9]
	d[i,8] = d[i,10]
	d[i,9] = t1
	d[i,10] = t2
	return(d)

}

plot_modelcov = function(stem, pop_order, min = -0.009, max = 0.009, cex = 1, usemax = T){
        c = read.table(gzfile(paste(stem, ".modelcov.gz", sep = "")), as.is = T, head = T)
        o = read.table(pop_order, as.is = T, comment.char = "", quote = "")


        toplot = data.frame(matrix(nrow = nrow(c), ncol = ncol(c)))
        for(i in 1:nrow(o)){
                for( j in 1:nrow(o)){

                        toplot[i, j] = c[which(names(c)==o[i,1]), which(names(c)==o[j,1])]
                }
        }
        if (usemax){
                m1 = max(abs(toplot))
                max = m1*1.1
                min = -(m1*1.1)
        }
        names(toplot) = o[,1]
        plot_resid_internal(toplot, max = max, min = min)
}


plot_cov = function(stem, pop_order, min = -0.009, max = 0.009, cex = 1, usemax = T, wcols = ""){
        c = read.table(gzfile(paste(stem, ".cov.gz", sep = "")), as.is = T, head = T)
        o = read.table(pop_order, as.is = T)


        toplot = data.frame(matrix(nrow = nrow(c), ncol = ncol(c)))
        for(i in 1:nrow(o)){
                for( j in 1:nrow(o)){

                        toplot[i, j] = c[which(names(c)==o[i,1]), which(names(c)==o[j,1])]
                }
        }
        if (usemax){
                m1 = max(abs(toplot))
                max = m1*1.1
                min = 0
        }
        names(toplot) = o[,1]
        plot_cov_internal(toplot, max = max, min = min, wcols = wcols, o = o, cex = cex)
}


plot_resid = function(stem, pop_order, min = -0.009, max = 0.009, cex = 1, usemax = T, wcols = "r"){
	c = read.table(gzfile(paste(stem, ".cov.gz", sep="")), as.is=T, head=T, quote="", comment.char="")
	m = read.table(gzfile(paste(stem, ".modelcov.gz", sep="")), as.is=T, head=T, quote="", comment.char="")
	names(c) = rownames(c)
	names(m) = rownames(m)
	o = read.table(pop_order, as.is=T, comment.char="", quote="")
	se = read.table(gzfile(paste(stem, ".covse.gz", sep="")), as.is=T, head=T, quote="", comment.char="")
	mse = apply(se, 1, mean)
	mse = mean(mse)
	print(mse)	
	c = c[order(names(c)), order(names(c))]
	m = m[order(names(m)), order(names(m))]
	tmp = c -m 
	#tmp = m - c
	#tmp = (m-c)/m
	#print(tmp)
	toplot = data.frame(matrix(nrow = nrow(tmp), ncol = ncol(tmp)))
	for(i in 1:nrow(o)){
	        for( j in 1:nrow(o)){
			#print(paste(o[i,1], o[j,1]))
			if (o[i,1] %in% names(tmp) ==F){
				print(paste("not found", o[i,1]))
			}
			if (o[j,1] %in% names(tmp) ==F){
				print(paste("not found", o[j,1]))
			}
        	        toplot[i, j] = tmp[which(names(tmp)==o[i,1]), which(names(tmp)==o[j,1])]
        	}
	}
	#print(toplot)
	if (usemax){
		m1 = max(abs(toplot), na.rm = T)
		max = m1*1.02
		min = -(m1*1.02)	
	}
	print("here")
	names(toplot) = o[,1]
	toreturn = plot_resid_internal(toplot, max = max, min = min, wcols = wcols, mse = mse, o = o, cex = cex)
	return(toreturn)
}

plot_cov_internal = function(d, o = NA, max = 0.009, min = -0.009, cex =0.5, wcols = "", mse = 5){
        npop = nrow(d)
        width = 1/npop
        height = 1/npop
        colors = brewer.pal(9, "Spectral")
        colors = c("red", "orange","yellow", "white", "green", "blue", "black")
        pal = colorRampPalette(colors)
        ncol = 80
        cols = pal(ncol)
        plot("NA", xlim = c(0, 1), ylim = c(0, 1), axes = F, xlab = "", ylab = "")
        for (i in 1:npop){
                for( j in 1:i){
                        v = d[i,j]
                        col= "white"
                        if (v < 0){
                                if (wcols == "rb"){
                                col = rgb(0, 0, 1, v/min)
                                }
                                else{
                                #col = rgb(0, 0, 1, 0.1+0.9*(v/min))
                                col = cols[ncol/2-floor( (v/min)*(ncol/2))]
                                }
                        }
                        else{
                                if (wcols == "rb"){
                                col = rgb(1, 0, 0, v/max)
                                }
                                else{
                                #col = rgb(1, 0, 0, 0.1+0.9*(v/max))
                                col = cols[ceiling((v/max)*(ncol))]
                                }
                        }
                        xmin = j/npop - 1/npop
                        xmax = j/npop
                        ymin = 1-(i/npop)
                        ymax = 1-(i/npop)+1/npop
			if (v == 0){ col = "white"}
                        rect(xmin, ymin, xmax, ymax, col = col, border = col)
                }
                tcol = "black"
                tmp = o[o[,1] == names(d)[i],]
                if (length(tmp) != 1){
                        tcol = tmp[1,2]
                }
                mtext(names(d)[i], side = 2, at = 1-i/npop+0.5/npop, las = 1, cex = cex, col = tcol)
                mtext(names(d)[i], side = 1, at =  i/npop-0.5/npop, las = 3, cex = cex, col = tcol)
        }
        if ( !is.na(mse)){
                ymi = 0.5
                yma = 0.9
                w = (yma-ymi)/ncol
                xma = 0.80
                lmi = round(min, digits = 1)
                lma = round(max, digits = 1)
                print(cols)
                print(ymi+(0:ncol)*w)
                rect( rep(0.75, ncol), ymi+(0:(ncol-1))*w, rep(xma, ncol), ymi+(1:ncol)*w, col=cols, border=cols)
                text(xma+0.01, ymi, lab = paste(lmi),  adj = 0, cex = 0.8)
                text(xma+0.01, yma, lab = paste(lma, "(Variance)"), adj = 0, cex = 0.8)

        }
        return(d)
        #image(as.matrix(d), col = cols)
}

plot_resid_internal = function(d, o = NA, max = 0.009, min = -0.009, cex =0.5, wcols = "rb", mse = NA){
        npop = nrow(d)
        width = 1/npop
        height = 1/npop
	colors = brewer.pal(9, "Spectral")
	colors = c("red", "orange","yellow", "white", "green", "blue", "black")
	pal = colorRampPalette(colors)
	ncol = 80
	cols = pal(ncol)
        plot("NA", xlim = c(0, 1), ylim = c(0, 1), axes = F, xlab = "", ylab = "")
        for (i in 1:npop){
                for( j in 1:i){
                        v = d[i,j]
			print(paste(i, j, v))
                        col= "white"
                        if (v < 0){
				if (wcols == "rb"){
				col = rgb(0, 0, 1, v/min)
				}
				else{
                                #col = rgb(0, 0, 1, 0.1+0.9*(v/min))
				col = cols[ncol/2-floor( (v/min)*(ncol/2))]
				#col = "white"
				}
                        }
                        else{
				if (wcols == "rb"){
				col = rgb(1, 0, 0, v/max)
				}
				else{
                                #col = rgb(1, 0, 0, 0.1+0.9*(v/max))
				col = cols[ncol/2+ceiling((v/max)*(ncol/2))]
				}
                        }
                        xmin = j/npop - 1/npop
                        xmax = j/npop
                        ymin = 1-(i/npop)
                        ymax = 1-(i/npop)+1/npop
                        rect(xmin, ymin, xmax, ymax, col = col, border = col)
                }
		tcol = "black"
		tmp = o[o[,1] == names(d)[i],]
		if (length(tmp) != 1){
			tcol = tmp[1,2]
		}
                mtext(names(d)[i], side = 2, at = 1-i/npop+0.5/npop, las = 1, cex = cex, col = tcol)
                mtext(names(d)[i], side = 1, at =  i/npop-0.5/npop, las = 3, cex = cex, col = tcol)
        }
	if ( !is.na(mse)){
                ymi = 0.5
                yma = 0.9
                w = (yma-ymi)/ncol
                xma = 0.80
		lmi = round(min/mse, digits = 1)
		lma = round(max/mse, digits = 1)
		print(cols)
		print(ymi+(0:ncol)*w)
                rect( rep(0.75, ncol), ymi+(0:(ncol-1))*w, rep(xma, ncol), ymi+(1:ncol)*w, col=cols, border=cols)
                text(xma+0.01, ymi, lab = paste(lmi, "SE"),  adj = 0, cex = 0.8)
                text(xma+0.01, yma, lab = paste(lma, "SE"), adj = 0, cex = 0.8)

	}
	return(d)
	#image(as.matrix(d), col = cols)
}

plot_tree_internal2 = function(d, e, o = NA, cex = 1, disp = 0.005, plus = 0.005,
                               arrow = 0.05, arrow.lwd = 1, ybar = 0.01, scale = T,
                               mbar = T, mse = 0.01, plotmig = T, plotnames = T, xmin = 0, lwd = 1, font = 1,
                               alpha.color=NULL, title=NULL, label=NULL, cex.main=1, xlab="", ...){
	plot(d$x, d$y, axes=F, ylab="", xlab=xlab, xlim = c(xmin, max(d$x)+plus), pch = "", ...)
	axis(1)
	mw = max(e[e[,5]=="MIG",4])
	mcols = rev(heat.colors(150))
	for(i in 1:nrow(e)){
		w = floor(e[i,4]*200)+50
		if (mw > 0.5){
			w = floor(e[i,4]*100)+50
		}
		#col = mcols[w]
            col <- alpha.color[i]
		v1 = d[d[,1] == e[i,1],]
		v2 = d[d[,1] == e[i,2],]
		if (e[i,5] == "MIG"){
			if (plotmig){
			Arrows( v1[1,]$x, v1[1,]$y, v2[1,]$x, v2[1,]$y, col = col, arr.width=arrow,
                          lwd = arrow.lwd, lty="dotted", arr.type="triangle")
			}
		}
		else{
			lines( c(v1[1,]$x, v2[1,]$x), c(v1[1,]$y, v2[1,]$y), col = col, lwd = lwd)
		}
	}
	tmp = d[d[,5] == "TIP",]
	print(tmp$x)
	print(disp)
	if ( !is.na(o)){
		for(i in 1:nrow(tmp)){
			tcol = o[o[,1] == tmp[i,2],2]
			if(plotnames){
				#print(tmp[i,2])
                        clabel <- tmp[i,2]
                        if(!is.null(label)){clabel <- label[i]}
				text(tmp[i,]$x+disp, tmp[i,]$y, labels=clabel, adj=0, cex=cex, col=tcol, font=font)
			}
		}
	}
	else{
		if (plotnames){
            clabels <- tmp[,2]
            if(!is.null(label)){clabels <- label}
		text(tmp$x+disp, tmp$y, labels = clabels, adj = 0, cex = cex, font = font)
		}
	}
	if (scale){
	print (paste("mse", mse))
        lines(c(0, mse*10), c(ybar, ybar))
	text( 0, ybar - 0.04, lab = "10 s.e.", adj = 0, cex  = 0.8)
	lines( c(0, 0), c( ybar - 0.01, ybar+0.01))
	lines( c(mse*10, mse*10), c(ybar- 0.01, ybar+ 0.01))
	}
        if (mbar){
                mcols = rev( heat.colors(150) )
                mcols = mcols[50:length(mcols)]
                ymi = ybar+0.15
                yma = ybar+0.35
                l = 0.2
                w = l/100
                xma = max(d$x/20)
                rect( rep(0, 100), ymi+(0:99)*w, rep(xma, 100), ymi+(1:100)*w, col=mcols, border=mcols)
                text(xma+disp, ymi, lab = "0", adj = 0, cex = 0.7)
		if ( mw >0.5){ text(xma+disp, yma, lab = "1", adj = 0, cex = 0.7)}
                else{
			text(xma+disp, yma, lab = "0.5", adj = 0, cex =0.7)
		}
		text(0, yma+0.06, lab = "Migration", adj = 0 , cex = 0.6)
		text(0, yma+0.03, lab = "weight", adj = 0 , cex = 0.6)
        }
      mtext(title, cex=cex.main, line=0.1)	
}

plot_tree2 = function(stem, o = NA, cex = 1, disp = 0.003, plus = 0.01, flip = vector(), arrow = 0.05,
                      arrow.lwd=1, scale = T, ybar = 0.1, mbar = T, plotmig = T, plotnames = T,
                      xmin = 0, lwd = 1, font = 1, alpha.color=NULL, title=NULL, label=NULL,
                      cex.main=1, xlab="", ...){
	d = paste(stem, ".vertices.gz", sep = "")
	e = paste(stem, ".edges.gz", sep = "")
	se = paste(stem, ".covse.gz", sep = "")
	d = read.table(gzfile(d), as.is = T, comment.char = "", quote = "")
	e = read.table(gzfile(e), as.is  = T, comment.char = "", quote = "")
	if (!is.na(o)){
		o = read.table(o, as.is = T, comment.char = "", quote = "")
	}
	e[,3] = e[,3]*e[,4]
	e[,3] = e[,3]*e[,4]
	
	se = read.table(gzfile(se), as.is = T, comment.char = "", quote = "")
	m1 = apply(se, 1, mean)
	m = mean(m1)
	#m = 0
	for(i in 1:length(flip)){
		d = flip_node(d, flip[i])
	}
	d$x = "NA"
	d$y = "NA"
	d$ymin = "NA"
	d$ymax = "NA"
	d$x = as.numeric(d$x)
	d$y = as.numeric(d$y)
	d$ymin = as.numeric(d$ymin)
	d$ymax = as.numeric(d$ymax)

	d = set_y_coords(d)
	d = set_x_coords(d, e)
	print(d)
	d = set_mig_coords(d, e)
	plot_tree_internal2(d, e, o=o, cex=cex, xmin=xmin, disp=disp, plus=plus, arrow=arrow,
                         arrow.lwd=arrow.lwd, ybar=ybar, mbar=mbar, mse=m, scale=scale,
                         plotmig=plotmig, plotnames=plotnames, lwd=lwd, font=font,
                         alpha.color=alpha.color, title=title, label=label, cex.main, ...)
	return(list( d= d, e = e))
}

alpha2col <- function(x, lim.x=NULL, edge){
  if(is.null(lim.x)){
    max.x <- max(x)
    min.x <- min(x)
    max.x <- max(abs(min.x), abs(max.x))
    min.x <- -max.x
  }else{
    max.x <- lim.x
    min.x <- -max.x
  }  
  colfunc <- colorRampPalette(c("blue","grey","red"),space="rgb")
  colrange <- seq(min.x*100,max.x*100,1)
  colpal <- colfunc(length(colrange))
  colpal <- cbind(colrange,colpal)
  colpal <- data.frame(colpal)
  colpal[,"colrange"] <- as.numeric(colpal[,"colrange"])
  breaks <- seq(min.x, max.x,length.out=200)
  x2 <- sapply(x, function(x){
    ccol <- colpal[which(round(colpal[,1]) == round(as.numeric(x)*100)),2]
    if(length(ccol>0)){ccol}else{if(x<0){colpal[1,2]}else{colpal[nrow(colpal),2]}}
  })
  order.edge <- paste0("alpha_", edge[,2], "_", edge[,1])
  x2 <- x2[order.edge]

  return(x2)
}
```
