## Supplementary Script for "Exploratory analysis of multiple traits co-adaptations in the population history": README.txt

R script used in the analysis of simulation. R system version 4.0.0 or newer is required.

HOW TO USE
1. Put all files and folders in a same working directory.
2. Start R in the working directory.
3. Open "SimMigrationE1_1way0mig_20211001.R" and execute all lines from the beginning.
