## Supplementary Script for "Exploratory analysis of multiple traits co-adaptations in the population history": SimFunction20211001.R

```
# genetic drift of neutral allele
makeP.neutral <- function(obsP, Ne){
  cnloci <- dim(obsP)[2]
  obsP2 <- array(NA, dim(obsP))
  colnames(obsP2) <- colnames(obsP)
  for(cpop in 1:npop){
    cNe <- Ne[cpop]
    cP <- obsP[cpop,]
    cP2 <- rbinom(cnloci,cNe,prob=cP)/cNe
    obsP2[cpop,] <- cP2
  }
  return(obsP2)
}

# generate "nbirth" individual genotypes fro allele freq "obsP"
makeG <- function(obsP, nbirth){
  cnloci <- dim(obsP)[2]
  cnpop <- dim(obsP)[1]
  obsG <- array(NA, c(npop, nbirth, cnloci))
  for(cpop in 1:cnpop){
    cP <- obsP[cpop,]
    cP <- cbind(1-cP, cP)
    cG <- apply(cP, 1, function(x){
           if(is.na(x[1])){ rep(NA,nbirth*2) }else{ sample(0:1, size=nbirth*2, rep=T,prob=x) }
         })
    cG <- cG[c(T,F),]+cG[c(F,T),]
    obsG[cpop,,] <- cG
  }
  return(obsG)
}

# generate traits from individual genotype "obsG"
makeTm <- function(obsG, testE){
  obsT <- array(NA, dim(obsG))
  cGET <- dim(obsG)[3]
  cnpop <- dim(obsG)[1]
  cE <- testE
  for(ct in 1:cGET){
    cG <- obsG[,,ct]
    cTe <- array(rnorm(n=length(cG),mean=0,sd=1), dim(cG))
    cT <- cG*gammaG + cE*gammaE + cG*cE*gammaGE + cTe
    cT <- (cT-min(cT,na.rm=T))/(max(cT,na.rm=T)-min(cT,na.rm=T))
    obsT[,,ct] <- cT
  }
  return(obsT)
}


# current gen allele freq (obsP) -> genotype (cG) -> trait (cT) -> survival rate (cS) -> next gen allele freq (obsP2)
survivePxm <- function(obsP, pop.exist, testE){
  cnpop <- dim(obsP)[1]
  obsP2 <- array(NA,dim(obsP))
  colnames(obsP2) <- colnames(obsP)
  loci.exist <- which(apply(obsP, 2, function(x){sum(!is.na(x))}) > 0)
  cE <- matrix(rep(testE, 3), byrow=F, nrow=cnpop)
  for(cg in loci.exist){
    cG <- array(NA,c(cnpop,3))
    cG[which(pop.exist),] <- rep(c(0,1,2),each=sum(pop.exist))
    cT <- cG*gammaG + cE*gammaE + cG*cE*gammaGE
    cS <- pnorm(q=0, mean=cT, sd=1, lower.tail=F)
    cP <- obsP[,cg]
    cGP <- cbind((1-cP)^2, 2*cP*(1-cP), cP^2)
    cGPs <- t(apply(cGP*cS, 1, function(x){x/sum(x)}))
    cPs <- sqrt(cGPs[,c(1,3)])
    cPs <- t(apply(cPs, 1, function(x){x/sum(x)}))
    obsP2[,cg] <- cPs[,2]
  }
  return(obsP2)
}

# migration to neighbouring place : Ne x mig.rate individuals
migratePxV <- function(obsP, Ne, mig.rate, pop.exist, vacant){
  cnloci <- dim(obsP)[2]
  cnpop <- dim(obsP)[1]
  cmig <- array(NA, dim(connect))
  cmig[,1] <- !is.na(match(connect[,1],which(pop.exist)))
  cmig[,2] <- !is.na(match(connect[,2],which(pop.exist)))
  cmig <- which(if(vacant){cmig[,1]|cmig[,2]}else{cmig[,1]&cmig[,2]})
  cNe <- Ne * pop.exist
  for(cmigi in cmig){
    cpop1 <- as.integer(connect[cmigi, 1])
    cpop2 <- as.integer(connect[cmigi, 2])
    cP1 <- obsP[cpop1,]
    cP2 <- obsP[cpop2,]
    cP1[is.na(cP1)] <- 0
    cP2[is.na(cP2)] <- 0
    cmr <- as.numeric(mig.rate[cmigi])
    cN1 <- cNe[cpop1]
    cN2 <- cNe[cpop2]
    n12 <- as.integer(cN1 * cmr); n1 <- cN1 - n12
    n21 <- as.integer(cN2 * cmr); n2 <- cN2-n21
    cP1m <- (rbinom(cnloci, n1, cP1) + rbinom(cnloci, n21, cP2)) / (n1 + n21)
    cP2m <- (rbinom(cnloci, n2, cP2) + rbinom(cnloci, n12, cP1)) / (n2 + n12)
    cP1m[is.nan(cP1m)] <- NA
    cP2m[is.nan(cP2m)] <- NA
    obsP[cpop1,] <- cP1m
    obsP[cpop2,] <- cP2m
  }
  return(obsP)
}

# mutaion of newly derived SNPs
mutateP <- function(npop, init.af, nloci.1step, Ne.scale, pop.exist, generation){
  nL1 <- nloci.1step * Ne.scale
  nL.all <- sum(nL1[pop.exist])
  output.newP <- array(NA,c(npop,nL.all))
  newloci.name <- rep("", nL.all)
  cloc <- 1
  for(cpop in which(pop.exist)){
    cnL1 <- nL1[cpop]
    newP <- array(NA,c(npop,cnL1))
    newP[pop.exist,] <- 0
    newP[cpop,] <- init.af
    output.newP[,cloc+(1:cnL1)-1] <- newP
    newloci.name[cloc+(1:cnL1)-1] <- paste0("g", generation, "p", cpop, "s",1:cnL1)
    cloc <- cloc + cnL1
  }
  colnames(output.newP) <- newloci.name
  return(output.newP)
}

# cut off SNPs to use by allelefreq
af.cut.off <- function(obsP, af.threshold){
  meanP <- colMeans(obsP,na.rm=T)
  obsP <- obsP[,meanP>af.threshold,drop=F]
  return(obsP)
}
maf.cut.off <- function(obsP, af.threshold){
  meanP <- colMeans(obsP,na.rm=T)
  for(cl in 1:length(meanP)){if(meanP[cl]>0.5){obsP[,cl] <- 1-obsP[,cl]}}
  meanP <- colMeans(obsP,na.rm=T)
  obsP <- obsP[,meanP>af.threshold,drop=F]
  return(obsP)
}
af.cut.off.pop <- function(obsP, af.threshold, pops=NULL){
  if(is.null(pops)){pops <- 1:nrow(obsP)}
  meanP <- colMeans(obsP[pops,],na.rm=T)
  obsP <- obsP[,meanP>af.threshold,drop=F]
  return(obsP)
}


# Output of simulated population in GENEPOP format
make_genepop <- function(obsP, obsG.gp, gpfile){
  outtxt <- paste0("\"pop=", npop, " snp=", ncol(obsP), " ", csim.label, "\"") 
  #snpnames <- paste0(paste0("snp",1:nloci))
  snpnames <- colnames(obsP)
  outtxt <- c(outtxt, paste0(snpnames, collapse="\n"))
  for(cpop in 1:npop){
    cgt <- obsG.gp[cpop,,]
    cgt <- apply(cgt, 2, function(x){
      x <- gsub("0","A",x)
      x <- gsub("1","H",x)
      x <- gsub("2","B",x)
      x <- gsub("A","0101",x)
      x <- gsub("H","0102",x)
      x <- gsub("B","0202",x)
    })
    cgt <- apply(cgt, 1, paste0, collapse="\t")
    cind <- paste0("p",cpop,"i",1:nind.gp)
    cgt <- paste0(cind, ", ", cgt)
    cgt <- c("POP",cgt)
    outtxt <- c(outtxt,cgt)
  }
  cat(outtxt, file=gpfile, sep="\n")
  #rm(obsG.gp);gc()
}
```
