## Supplementary Script for "Exploratory analysis of multiple traits co-adaptations in the population history": SimMigrationE1_1way0mig_20211001.R

```
# Copyright (C) Reiichiro Nakamichi
# License: http://www.gnu.org/licenses/gpl.html GPL version 2 or higher


# for main analysis
library(FinePop2)
source("FinePop2Ext20210817.R")
library(gtools)
library(MASS)
library(ape)
library(foreach)
library(doParallel)
source("SimFunction20211001.R")

# for TREEMIX
library(RColorBrewer)
source("TreemixFunction20211001.R")

# for PolyGraph
source("admixturegraphFunction20210927.R")
library(msm)
library(reshape2)
library(pscl)
library(ggplot2)
source("PolygraphFunction20211001.R")

# simulation setting
map.file <- "map_1way_0mig.txt"    # map file of population expansion senario
baseNe <- 1e4                # Ne of each population
scale.ancestor <- 10         # scale of ancestral population (Ne of ancestral pop = [baseNe] x [scale.ancestor])
num.gen <- 260               # number of generations to be simulated �inum.gen must be integer multiplier of [interval.mig] �j
interval.mig <- 10           # population expansion to a vacant deme will be occured every [interval.mig] generations
num.loci.ancestor <- 10000   # observed number of SNPs in the ancestral population
num.loci.new.neutral <- 500  # observed number of newly derived neutral SNPs after [num.gen] generations
num.loci.new.adaptive <- 10  # observed number of newly derived adaptive SNPs after [num.gen] generations for each trait
init.af <- 0.01              # initial allele freq of newly derived SNPs
num.ind <- 50                # number of individuals to be sampled after [num.gen] generations


#####################################
### Simulation ###

### simulation parameters
# R.GE, R.E: ratio of survivability(S) 
# S.G0E1/S.G0E0 = R.E (0 < R.E < 1)
# S.G2E0/S.G0E0 = 1/R.GE  (1 <= R.GE)
# S.G2E1/S.G0E1 = R.GE

## map & capacity
map <- as.matrix(read.table(map.file))   
pops <- sort(unique(as.integer(map[,c(1,2)])))
npop <- length(pops)
connect <- map[,c(1,2)]
mig.rateV <- map[,3]
mig.rate <- rep(0, length(mig.rateV))

Ne <- rep(baseNe, npop) # effectiv pop size
Ne[1] <- baseNe * scale.ancestor
Ne.scale <- Ne/min(Ne)

## environment
nE <- 1                        # number of env factors
obsE <- array(0, c(npop,nE))   # 0=normal or 1=severe
obsE[c(9,15),1] <- 1           # deme 9 and 15 have severe condition
rownames(obsE) <- 1:25
obsE.order <- as.character(1:npop)
obsE.rorder <- as.integer(rownames(obsE))

obsE <- apply(obsE, 2, function(x){(x-min(x))/(max(x)-min(x))})
colnames(obsE) <- paste0("E",1:nE)

### survival parameter
R.E <- 0.5
R.GE <- 1.1   # s=0.1

gammaE <- qnorm(R.E*0.5)
gammaG <- 0.5 * qnorm(1/R.GE * 0.5)
gammaGE <- 0.5 * (qnorm(0.5*R.E*R.GE) - qnorm(R.E*0.5) - qnorm(1/R.GE * 0.5) )

eff.cutoff <- 1e-10
if(abs(gammaG) < eff.cutoff){gammaG <- 0}
if(abs(gammaE) < eff.cutoff){gammaE <- 0}
if(abs(gammaGE) < eff.cutoff){gammaGE <- 0}

## setting for newly derived SNPs
ngen.step <- interval.mig
nstep  <- as.integer(num.gen / interval.mig)

dist.from1 <- rep(0,npop)
for(cpop in 2:npop){
  cpar <- map[map[,2]==cpop,1]
  cdist <- 1
  while(min(cpar, na.rm=T)>1){
    cpar <- map[map[,2]==cpar,1]
    cdist <- cdist + 1
  }
  dist.from1[cpop] <- cdist
}
nchance.new <- sum((nstep - dist.from1) * ngen.step * Ne.scale)
rchance.popE <- sum(obsE)/length(obsE)
nloci.ancestor <- num.loci.ancestor * 4
nloci.1step.adaptive <- as.integer(num.loci.new.adaptive / init.af / nchance.new / rchance.popE) + 5
nloci.1step.neutral <- as.integer(num.loci.new.neutral / init.af / nchance.new) + 10

csim.label <- paste0("RE",sprintf("%.1f",R.E),"RGE",sprintf("%.4f",R.GE))


### Population expansion ###
## ancestral SNPs
q1 <- 1/Ne[1]
q2 <- 1-1/Ne[1]
C <- 1/log((q2*(1-q1))/(q1*(1-q2)))
d <- log(q1/(1-q1))
u <- runif(nloci.ancestor)
dist.af.ancestor <- 1/(1+exp(-(u/C+d)))
rm(q1,q2,C,d,u);gc()
obsPa <- array(NA, c(npop, nloci.ancestor)) # SNP: ancestor
obsPa[1,] <- apply(cbind(dist.af.ancestor,1-dist.af.ancestor), 1, min)

## newly derived SNPs
obsPmGET1 <- NULL # GET, E1
obsPmN <- NULL    # neutral

## generations (!!! several ten minits required !!!)
message(csim.label, " gammaG=", gammaG, " gammaGE=", gammaGE)
cutoff.0 <- 1/baseNe * 0.1
for(cstep in 1:nstep){
  message(rep("\b",10),cstep,"/",nstep,appendLF=F);flush.console()
  pop.exist <- !is.na(obsPa[,1])
  for(cgen in 1:ngen.step){
    cgen.total <- (cstep-1)*ngen.step+cgen
    obsPmGET1 <- cbind(obsPmGET1, mutateP(npop, init.af, nloci.1step.adaptive, Ne.scale, pop.exist, cgen.total))
    obsPmN <- cbind(obsPmN, mutateP(npop, init.af, nloci.1step.neutral, Ne.scale, pop.exist, cgen.total))

    obsPa <- migratePxV(obsPa, Ne, mig.rate, pop.exist, vacant=F)
    obsPmGET1 <- migratePxV(obsPmGET1, Ne, mig.rate, pop.exist, vacant=F)
    obsPmN <- migratePxV(obsPmN, Ne, mig.rate, pop.exist, vacant=F)

    obsPmGET1 <- survivePxm(obsPmGET1, pop.exist, obsE[obsE.order,"E1"])

    obsPa <- makeP.neutral(obsPa, Ne)
    obsPmGET1 <- makeP.neutral(obsPmGET1, Ne)
    obsPmN <- makeP.neutral(obsPmN, Ne)
  }
  obsPa <- migratePxV(obsPa, Ne, mig.rateV, pop.exist, vacant=T)
  obsPmGET1 <- migratePxV(obsPmGET1, Ne, mig.rateV, pop.exist, vacant=T)
  obsPmN <- migratePxV(obsPmN, Ne, mig.rateV, pop.exist, vacant=T)

  obsPa <- af.cut.off(obsPa, af.threshold=cutoff.0)
  obsPmGET1 <- af.cut.off(obsPmGET1, af.threshold=cutoff.0)
  obsPmN <- af.cut.off(obsPmN, af.threshold=cutoff.0)
}
message(" done.")

cutoff.keep <- 1/baseNe
obsPa.cut <- af.cut.off(obsPa, af.threshold=cutoff.keep)
obsPmGET1.cut <- af.cut.off(obsPmGET1, af.threshold=cutoff.keep)
obsPmN.cut <- af.cut.off(obsPmN, af.threshold=cutoff.keep)

colnames(obsPa.cut) <- paste0("A",1:ncol(obsPa.cut))
colnames(obsPmGET1.cut) <- paste0("M1",colnames(obsPmGET1.cut))
colnames(obsPmN.cut) <- paste0("N",colnames(obsPmN.cut))

## test of sim reult
cutoff.use <- 0.01
obsPa.test <- af.cut.off(obsPa.cut, af.threshold=cutoff.use)
obsPmGET1.test <- af.cut.off(obsPmGET1.cut, af.threshold=cutoff.use)
obsPmN.test <- af.cut.off(obsPmN.cut, af.threshold=cutoff.use)
message("A=", ncol(obsPa.test), " N=", ncol(obsPmN.test), " GET1=", ncol(obsPmGET1.test))
rowMeans(obsPa.test)
rowMeans(obsPmN.test)
rowMeans(obsPmGET1.test)

## select SNPs to use
cutoff.use <- 0.01
obsPa.cut <- maf.cut.off(obsPa.cut, af.threshold=cutoff.use)
obsPmGET1.cut <- maf.cut.off(obsPmGET1.cut, af.threshold=cutoff.use)
obsPmN.cut <- maf.cut.off(obsPmN.cut, af.threshold=cutoff.use)
message(csim.label, " A=", ncol(obsPa.cut), " N=", ncol(obsPmN.cut), " GET=", ncol(obsPmGET1.cut))


###################################################
### gene-environment-trait association analysis ###
nind.gp <- 50       # number of sampled individuals per deme

# loci: neutral & analyze
nGETs <- 10
nNs <- 50
nAs <- 500
nAsC <- 5000
nloci <- nGETs*nE + nNs + nAs
nlociC <- nGETs*nE + nNs + nAsC

selectP <- function(obsP,n.select){obsP[,sort(sample(1:ncol(obsP),n.select))]}
selectPN <- function(obsP,n.select,loc){
  locs <- unlist(strsplit(colnames(obsP), split="p"))[c(F,T)]
  locs <- as.integer(unlist(strsplit(locs, split="s"))[c(T,F)])
  locs.hit <- sample(which((loc-1)<=locs & locs<=(loc+1)), n.select)
  obsP[,sort(locs.hit)]
}
obsPmGET1.use <- cbind(selectPN(obsPmGET1.cut,nGETs/2,9), selectPN(obsPmGET1.cut,nGETs/2,15))
obsPmN.use <- selectP(obsPmN.cut,nNs)
CobsPa.use <- selectP(obsPa.cut,nAsC)
obsPa.use <- CobsPa.use[,1:nAs]
colnames(obsPmGET1.use)

# all SNPs
AobsP <- cbind(obsPmGET1.use, obsPmN.use, CobsPa.use)
AobsG.gp <- makeG(AobsP, nind.gp)
Agpfile <- paste0("A_GETsim_SNP", nlociC, "_", csim.label, "_g", ngen.step, "E1d_0mig_genepop.txt")
make_genepop(AobsP, AobsG.gp, Agpfile)
popdataA <- read.GENEPOP(Agpfile) 

# neutral SNPs
CobsP <- cbind(obsPmN.use, CobsPa.use)
CobsG.gp <- makeG(CobsP, nind.gp)
Cgpfile <- paste0("C_GETsim_SNP", nlociC, "_", csim.label, "_g", ngen.step, "E1d_0mig_genepop.txt")
make_genepop(CobsP, CobsG.gp, Cgpfile)
popdataC <- read.GENEPOP(Cgpfile) 

# SNPs to analyse
obsP <- cbind(obsPmGET1.use, obsPmN.use, obsPa.use)
obsG.gp <- makeG(obsP, nind.gp)
gpfile <- paste0("GETsim_SNP", nloci, "_", csim.label, "_g", ngen.step, "E1d_0mig_genepop.txt")
make_genepop(obsP, obsG.gp, gpfile)
popdata <- read.GENEPOP(gpfile) 

# trait
obsT1.gp <- makeTm(CobsG.gp[,,1:(nGETs/2),drop=F], obsE[obsE.order,"E1"])
obsT2.gp <- makeTm(CobsG.gp[,,(1:(nGETs/2))+nGETs/2,drop=F], obsE[obsE.order,"E1"])
obsT1.gp <- apply(obsT1.gp, c(1,2), sum)
obsT2.gp <- apply(obsT2.gp, c(1,2), sum)
obsT.gp <- array(0, c(dim(obsT1.gp),2), dimnames=list(rownames(obsT1.gp), colnames(obsT1.gp), c("T1","T2")))
obsT.gp[,,"T1"] <- obsT1.gp
obsT.gp[,,"T2"] <- obsT2.gp
obsT1.pop <- apply(obsT1.gp, 1, mean)
obsT2.pop <- apply(obsT2.gp, 1, mean)
obsT.pop <- data.frame(T1=obsT1.pop, T2=obsT2.pop)
rownames(obsT.pop) <- 1:npop
nT <- ncol(obsT.pop)


## pop-specific FST using neutral SNPs
res.psfst <- pop_specificFST(popdataC, cov=F)
psfst <- res.psfst$fst[,"FST"]
names(psfst) <- popdata$pop_names

x11()
x <- barplot(psfst[obsE.rorder], las=3, ylim=c(0,0.2))
y.psfst <- psfst[obsE.rorder]
se.psfst <- res.psfst$fst[obsE.rorder,"SE"]
segments(x0=x, x1=x, y0=y.psfst-se.psfst*1.96, y1=y.psfst+se.psfst*1.96)

## GWAS for crrespondence analysis
cov.pop.moment <- cov.pop.mafW3(popdataA)
n.pop <- popdata$num_pop
data.get <- GET.data(popdata, obsE[obsE.order,,drop=F], obsT.pop)
result.gls <- GET.GLS(data.get, cov.pop.moment$cov.maf)
result.fdr <- GET.FDR(result.gls, fdr=0.05)

## correspondence analysis
result.crp <- GET.corresp(data.get)
data.get.attr <- GET.data.attribute(data.get)
data.get.attr[data.get.attr[,"type"]=="environment","col"] <- "green"
data.get.attr[data.get.attr[,"type"]=="trait","col"] <- "orange"
data.get.attr[grep("M",rownames(data.get.attr)),"col"] <- "purple"
data.get.attr[grep("A",rownames(data.get.attr)),"col"] <- "gray"
data.get.attr[grep("N",rownames(data.get.attr)),"col"] <- "gray"

col.pop <- (psfst-min(psfst))/(max(psfst)-min(psfst))
col.pop <- rgb(1-col.pop,0,col.pop)

lim.xy <- rbind(result.crp$rscore,result.crp$cscore)
lim.xy <- rbind(apply(lim.xy, 2, min),apply(lim.xy, 2, max))
lim.xy <- lim.xy*1.1
dimnames(lim.xy) <- list(c("min","max"),c("x","y"))

node.fdr <- unique(as.character(unlist(result.fdr[,c("environment","trait_gene")])))

attr <- data.get.attr[node.fdr,]
node.fdr.env <- unique(as.character(result.fdr[,c("environment")]))
node.fdr.gt <- unique(as.character(result.fdr[,c("trait_gene")]))
attr.e <- attr[node.fdr.env,]
attr.ne <- attr.e
attr.e[,"label"] <- paste0("+", attr.e[,"label"])
rownames(attr.ne) <- paste0("-",rownames(attr.ne))
attr.ne[,"label"] <- paste0("-", attr.ne[,"label"])
attr.gt <- attr[node.fdr.gt,]
attr <- rbind(attr.e, attr.ne, attr.gt)
font.e <- rep(2, nrow(attr.e))
font.ne <- rep(4, nrow(attr.ne))
font.gt <- rep(3, nrow(attr.gt))
font.gt[attr.gt[,"type"]=="trait"] <- 2
attr <- cbind(attr, font=c(font.e, font.ne, font.gt))
node.draw <- rownames(attr)
node.draw.et <- node.draw[attr[,"type"]=="environment"|attr[,"type"]=="trait"]

x11();par(mar=c(2.5, 2.5, 0.5, 0.5), mgp=c(1.5,0.5,0))
plot(result.crp$cscore, xlim=lim.xy[,"x"], ylim=lim.xy[,"y"], col="gray", pch=19,
     cex=min(attr[,"size"]), xlab="1st axis",ylab="2nd axis")
points(0,0, pch=3, cex=5)
for(cedge.i in 1:nrow(result.fdr)){
  cedge <- result.fdr[cedge.i,]
  cgt <- as.character(cedge[,"trait_gene"])
  cenv <- as.character(cedge[,"environment"])
  if(cedge[,"z"]<0){cenv <- paste0("-",cenv)}
  cxy <- result.crp$cscore[c(cenv,cgt),]
  lines(x=cxy[,1], y=cxy[,2], col="gray")
}
points(result.crp$cscore[node.draw,], col=as.character(attr[,"col"]),
       pch=19, cex=attr[,"size"])
text(result.crp$cscore[node.draw.et,], labels=as.character(attr[node.draw.et,"label"]),
     font=as.integer(attr[node.draw.et,"font"]))
text(result.crp$rscore, labels=1:n.pop, col=col.pop)


### TREEMIX ###  
# Preclaculated result of TREEMIX is in "treemix_C_GETsim_SNP5060_RE0.5RGE1.1000_g10E1d_0mig" folder.

Ctreemixfile <- paste0("C_GETsim_SNP", nlociC, "_", csim.label, "_g", ngen.step, "E1d_0mig_treemix.txt")
treemixdir <- paste0("treemix_C_GETsim_SNP", nlociC, "_", csim.label, "_g", ngen.step, "E1d_0mig")

make_treemixfile(popdataC, Ctreemixfile)

### Gzip "Ctreemixfile", set the zipped file in "treemixdir", and apply TREEMIX. ###
# treemix -i $zipped_treemixfile -root pop1 -o tm_mig0


### PolyGraph ###
## topology data: treemix to polygraph 
x11()
tm.stem <- paste0(treemixdir, "/tm_mig0")
tm.tree <- plot_tree(tm.stem, arrow=0.1, arrow.lwd=2, mbar=F, disp=0.0005)

pg.tree <- treemix2polygraph(tm.tree)

## GWAS for PolyGraph
# Precalculated result of GWAS is in "gwas_C_GETsim_SNP5060_RE0.5RGE1.1000_g10E1d_0mig" folder.

# folder to save gwas result in polygraph format
gwasfile.dir <- paste0("gwas_C_GETsim_SNP", nlociC, "_", csim.label, "_g", ngen.step, "E1d_0mig")

lociALL <- popdataA$loci_names
lociGET.i <- grep("M", lociALL)
lociGET <- lociALL[lociGET.i]
dimnames(AobsG.gp) <- list(NULL,NULL,lociALL)
gwas.effect <- array(NA, c(length(lociALL), ncol(obsT.pop)), dimnames=list(lociALL, colnames(obsT.pop)))
for(ct in colnames(gwas.effect)){
  cy <- as.numeric(obsT.gp[,,ct])
for(cg in rownames(gwas.effect)){
  cx <- as.numeric(AobsG.gp[,,cg])
  clm <- summary(lm(cy~cx))
  gwas.effect[cg, ct] <- clm$coefficients["cx","t value"]
}}

gwas.sig <- list()
gwas.sig[["T1"]] <- lociGET[1:5]
gwas.sig[["T2"]] <- lociGET[6:10]

for(cti in 1:length(gwas.sig)){
  ct <- names(gwas.sig)[cti]
  ct.gene <- gwas.sig[[ct]]
  cgwasfile <- paste0(gwasfile.dir, "/gwas_", ct, ".txt")
  outtxt <- paste(c("CHROM", "POS", "SNPID", "EFFECT", popdataC$pop_names), collapse="\t")
  for(cg in ct.gene){
    cgi <- which(popdataA$loci_names==cg)
    cg.count <- popdataA$allele_count[[cgi]]
    cg.count <- apply(cg.count, 2, function(x){paste(x,collapse=",")})
    ceff <- gwas.effect[cg, ct]
    outtxt <- c(outtxt, paste(c(NA,NA,cg,ceff,cg.count),collapse="\t"))
  }
  cat(outtxt, file=cgwasfile, sep="\n")

  cneutfile <- paste0(gwasfile.dir, "/neut_", ct, ".txt")
  outtxt <- paste(c("CHROM", "POS", "SNPID", "EFFECT", popdata$pop_names), collapse="\t") 
  for(cg in setdiff(lociALL, ct.gene)){
    cgi <- which(popdataA$loci_names==cg)
    cg.count <- popdataA$allele_count[[cgi]]
    cg.count <- apply(cg.count, 2, function(x){paste(x,collapse=",")})
    ceff <- gwas.effect[cg, ct]
    outtxt <- rbind(outtxt, paste(c(NA,NA,cg,ceff,cg.count),collapse="\t"))
  }
  cat(outtxt, file=cneutfile, sep="\n")
}

### PolyGraph MCMC ###
# Precalculated result of PolyGraph is in "polygraph_C_GETsim_SNP5060_RE0.5RGE1.1000_g10E1d_0mig" folder.

# folder to save polygraph result
polygraph.dir <- paste0("polygraph_C_GETsim_SNP", nlociC, "_", csim.label, "_g", ngen.step, "E1d_0mig")

# polygraph parameters
numsteps <- 30000
numsample <- 300
innerfreqs_proposize <- 0.1
alpha_prior_stdev <- 0.1
alpha_proposize <- 0.02
ssfactor <- 25
qbfactor <- 3
pvalcutoff <- 0.05
rootprior <- "unif"
runmode <- "alphas"
branchcandidate <- NaN
pvaltotal <- 0.05 

# num thread for MCMC calculation
nCPU <- 2

# select traits with significantly associated SNPs
trait.use.list.pg <- names(gwas.sig)
for(cti in 1:length(trait.use.list.pg)){
  target_trait <- trait.use.list.pg[cti]
  gwasfile <- paste0(gwasfile.dir, "/gwas_", target_trait, ".txt")
  cgwas <- scan(gwasfile, character(), sep="\n")
  if(length(cgwas)<=2){trait.use.list.pg[cti] <- NA}
}
trait.use.list.pg <- trait.use.list.pg[!is.na(trait.use.list.pg)]

#�@qstat file of PolyGraph
for(cti in 1:length(trait.use.list.pg)){
  target_trait <- trait.use.list.pg[cti]
  gwasfile <- paste0(gwasfile.dir, "/gwas_", target_trait, ".txt")
  neutfile <- paste0(gwasfile.dir, "/neut_", target_trait, ".txt")
  qfile <- paste0(polygraph.dir, "/qfile_", target_trait, ".txt")
  calc_qstat(gwasfile, neutfile, qfile, pg.tree)
}

#�@MCMC of PolyGraph (!!! several minits required !!!)
cl <- makeCluster(nCPU)
registerDoParallel(cl)
test.par <- foreach(cti=1:length(trait.use.list.pg), .packages="msm") %dopar% {
  target_trait <- trait.use.list.pg[cti]
  gwasfile <- paste0(gwasfile.dir, "/gwas_", target_trait, ".txt")
  neutfile <- paste0(gwasfile.dir, "/neut_", target_trait, ".txt")
  mcmcfile <- paste0(polygraph.dir, "/MCMC_trace_", target_trait, ".txt")

  leaves <- pg.tree$supergraph[[1]]$leaves
  data <- LoadCounts(gwasfile, leaves)
  leaves_counts <- as.data.frame(data[,seq(5,dim(data)[2])])
  effects <- as.numeric(data[,4])
  data <- LoadCounts(neutfile, leaves)
  neut_leaves_counts <- as.data.frame(data[,seq(5,dim(data)[2])])

  test.polygraph <- PolyGraph(mcmcfile,
    leaves_counts, neut_leaves_counts, effects, runmode, branchcandidate, numsteps, numsample,
    innerfreqs_proposize, alpha_prior_stdev, alpha_proposize, ssfactor, qbfactor, rootprior, pg.tree=pg.tree)
}
stopCluster(cl)

# polygraph alpha post distribution
alpha.list <- list()
for(cti in 1:length(trait.use.list.pg)){
  message(cti, " ", appendLF=F);flush.console()
  target_trait <- trait.use.list.pg[cti]
  mcmcfile <- paste0(polygraph.dir, "/MCMC_trace_", target_trait, ".txt")
  mcmc.trace <- read.table(mcmcfile,header=TRUE)
  mcmc.trace <- mcmc.trace[seq(1,dim(mcmc.trace)[1]-1,1),]
  alphacols <- grep("alpha_",colnames(mcmc.trace))
  calpha <- mcmc.trace[,alphacols]
  alpha.list[[target_trait]] <- calpha
  selcoefs <- apply(calpha,2,function(x){mean(x)})
}
alpha.mean <- NULL
for(cti in 1:length(trait.use.list.pg)){
  target_trait <- trait.use.list.pg[cti]
  calpha <- alpha.list[[target_trait]]
  calpha <- calpha[-(1:10),]  # remove burn-in
  alpha.mean <- cbind(alpha.mean, colMeans(calpha, na.rm=T))
}
colnames(alpha.mean) <- trait.use.list.pg

# alpha boxplot
pdffile <- paste0(polygraph.dir, "/polygraph_alpha_postdist.pdf")
pdf(pdffile, width=8, height=8)
for(cti in 1:length(trait.use.list.pg)){
  target_trait <- trait.use.list.pg[cti]
  calpha <- alpha.list[[target_trait]] 
  calpha <- calpha[-(1:10),]  # remove burn-in
  titleboxplot <- paste0("Posterior distributions of alpha: ", target_trait)
  alphacols <- which(sapply(colnames(calpha),function(x){grep("alpha_",x)}) == 1)
  data.alpha <- c()
  for(i in alphacols){data.alpha <- cbind(data.alpha,jitter(calpha[,i],0.1))}
  colnames(data.alpha) <- colnames(calpha)[alphacols]
  colnames(data.alpha) <- sapply(colnames(data.alpha),function(x){gsub("alpha_","",x)})
  data.alpha <- melt(as.data.frame(data.alpha))
  colnames(data.alpha) <- c("branch","alpha")
  plot_alpha <- ggplot(data.alpha, aes(branch, alpha)) +
                 geom_boxplot(width=0.4,fill="light blue") +
                 geom_hline(yintercept=0,colour="black") +
                 ggtitle(titleboxplot) +
                 theme(plot.title = element_text(hjust=0.5)) +
                 theme(axis.text.x = element_text(angle=90, hjust=1, vjust=0.5))
  #x11()
  plot(plot_alpha)
}
dev.off()


### PolyGraph alpha on TREEMIX
#pdffile <- paste0("polygraph_alpha_treemix_E1d_0mig.pdf")
#pdf(pdffile, width=8, height=8)
x11():par(mfrow=c(1,2))
for(cti in 1:length(trait.use.list.pg)){
  target_trait <- trait.use.list.pg[cti]
  #x11()
  plot_tree2(tm.stem, arrow=0.2, arrow.lwd=3, scale=T, mbar=F, lwd=3, title=target_trait, disp=0.001,
             alpha.color=alpha2col(x=alpha.mean[,target_trait], edge=pg.tree$edge))
}
#dev.off()


### Polygraph alpha PCA (TREEMIX all node)
# cor env vs trait
env.pop <- obsE[obsE.order,"E1"]
cordata.env <- array(rep(env.pop, nind.gp), dim(obsT.gp[,,"T1"]))
cordata.trait <- obsT.gp
cordata.env <- data.frame(E1=as.numeric(cordata.env))
cordata.trait <- data.frame(T1=as.numeric(obsT.gp[,,"T1"]), T2=as.numeric(obsT.gp[,,"T2"]))
cor.mat.et <- array(NA, c(ncol(cordata.env),ncol(cordata.trait)),
                   dimnames=list(colnames(cordata.env),colnames(cordata.trait)))
for(ct in colnames(cordata.trait)){
for(ce in colnames(cordata.env)){
  ctv <- cordata.trait[,ct]
  cev <- cordata.env[,ce]
  cor.mat.et[ce,ct] <- cor(ctv,cev)
}}

## pca alpha vs trait
pcadata.alpha <- alpha.mean
pcares.alpha <- prcomp(pcadata.alpha, rank=2)
x11();plot.pca.alpha(pcadata.alpha, pcares.alpha, title="Polygraph alpha vs trait", pop)
summary(pcares.alpha)

## pca alpha-env vs trait
pcadata.alphaenv <- rbind(alpha.mean, cor.mat.et[,colnames(alpha.mean),drop=F])
pcares.alphaenv <- prcomp(pcadata.alphaenv)
x11();plot.pca.alpha(pcadata.alphaenv, pcares.alphaenv, title="Polygraph alpha-environment vs trait")
summary(pcares.alphaenv)


### Polygraph alpha PCA (TREEMIX pop node) !!! used in article !!!
pop.names <- popdata$pop_names
env.pop <- obsE
rownames(env.pop) <- pop.names
alpha.pop <- array(NA, dim=c(length(pop.names),ncol(alpha.mean)))
dimnames(alpha.pop) <- list(pop.names, colnames(alpha.mean))
for(cp in pop.names){
  calpha <- alpha.mean[grep(paste0(cp,"_"), rownames(alpha.mean)),,drop=F]
  alpha.pop[cp,] <- colMeans(calpha)
}

pcadata.alphaenv.pop <- scale(cbind(alpha.pop,env.pop))
col.alphaenv <- c(rep("red", ncol(alpha.pop)), rep("darkgreen", ncol(env.pop)))
pcares.alphaenv.pop <- prcomp(pcadata.alphaenv.pop)
x11();plot.pca.pop(pcadata.alphaenv.pop, pcares.alphaenv.pop, col.score=col.alphaenv, PropVar=T)
summary(pcares.alphaenv.pop)


### Polygraph alpha PCA on treemix
# alpha (all node)
pca.score.alpha <- pcares.alpha$x
x11();par(mfrow=c(1,2))
for(i in 1:2){
  plot_tree2(tm.stem, arrow=0.1, arrow.lwd=3, lwd=3, title=paste0("PCA", i), mbar=F, scale=F, disp=0.001,
             alpha.color=alpha2col(x=pca.score.alpha[,i], edge=pg.tree$edge))
}

# alpha-env (all node)
pca.score.alphaenv <- pcares.alphaenv$x
x11();par(mfrow=c(1,2))
for(i in 1:2){
  plot_tree2(tm.stem, arrow=0.1, arrow.lwd=3, lwd=3, title=paste0("PCA", i), mbar=F, scale=F, disp=0.001,
             alpha.color=alpha2col(x=pca.score.alphaenv[,i], edge=pg.tree$edge))
}

# alpha (pop node) !!! used in article !!!
pca.score.alpha.pop <- pcares.alpha$x
is.pop <- rep(F, nrow(pca.score.alpha.pop))
for(cp in pop.names){
  is.pop[grep(paste0(cp,"_"), rownames(pca.score.alpha.pop))] <- T
}
pca.score.alpha.pop[!is.pop,] <- 0
x11();par(mfrow=c(1,2))
for(i in 1:2){
  plot_tree2(tm.stem, arrow=0.1, arrow.lwd=3, lwd=3, title=paste0("PCA", i), mbar=F, scale=F, disp=0.001,
             alpha.color=alpha2col(x=pca.score.alpha.pop[,i], edge=pg.tree$edge))
}
```
